# β-Hairpin Fold Peptide Brush Polymers

**DOI:** 10.64898/2026.10.02.756272

**Authors:** Julia Oktawiec, Hassan Nadeem, Omar M. Ebrahim, Yu Chen, Clara Barbut, James Byrnes, Matthew P. Thompson, Diwakar Shukla, Nathan C. Gianneschi

## Abstract

Peptide-brush polymers prepared via graft-through polymerization of peptide-based olefinic monomers form Protein-Like Polymers (PLPs) with demonstrated utility as chemical biology tools for modulating intracellular protein–protein interactions and are in development as a emerging class of therapeutic. This favorable bioactivity has been achieved in part by mimicking some of the structural and functional features of proteins, such as their dynamic, globular solution-phase structures, ability to insert into lipid membranes and enter cells, and tendency to enforce α-helicity in peptide side chains. However, because structure leads to function, insight into the structure of these proteomimetic polymers, particularly their overall tertiary structure and the extent to which this influences the brush peptide structure, is incomplete. To further this understanding, we prepared a set of brush polymers incorporating the trpzip2 peptide sequence, commonly used as a model system for protein folding. We examine these polymers via circular dichroism spectroscopy, infrared spectroscopy, and small-angle X-ray scattering, coupled with insights from molecular dynamics simulations. Here, a loss of the β-hairpin structure of the monomer is observed regardless of polymer composition, and simulations corroborate that the polymer backbone destabilizes local secondary structure, restricting peptide motion and altering the accessible free-energy landscape. This study provides insight into design rules governing these synthetic polymer-based proteomimetics.

## Introduction

Peptide brush polymers, or Protein-Like Polymers (PLPs), prepared via graft-through polymerization of peptides coupled to polymerizable moieties are emerging as useful proteomimetics for the development of intracellular tools for modulating cellular processes and as potential therapeutics.^1–3^ PLPs collapse into globular structures that resist proteolysis giving rise to excellent pharmacokinetic and pharmacodynamic properties and can passively penetrate cells, while also incorporating bioactive peptides for multivalent, potent and selective target protein engagement.^4,5^ PLPs are in development as therapeutics for a range of conditions spanning cancer, neurodegenerative disease, and cardiovascular disease.^6–11^

While these systems effectively engage with proteins of interest, a number of questions remain in understanding their structure for proteomimetic design more broadly. For example, can such polymer backbones support, enforce, or disrupt secondary structure in peptide side chains, including α-helices and β-sheets? Toward this, we have demonstrated that α-helices are maintained when incorporated as brushes, and indeed, their helicity is enhanced by densely grafting into peptide brush polymers.^12^ Here, we examine how β-sheet-like hydrogen bonding and β-hairpin folds behave when conjugated as dense polymer brushes. The β-hairpin fold allows the peptide backbone to fold on itself to form intramolecular hydrogen bonds in a β-sheet-like arrangement, and many proteins contain such folds, making them a critical class of secondary structure.

As a test case, we employed peptides whose β-hairpin fold is stabilized by tryptophan pairs, which arrange to form aromatic interactions, called tryptophan zippers, or “trpzips.”^13^ These π–π stacking interactions help stabilize the hairpin fold in aqueous solution. Additionally, the aromatic interactions yield a characteristic signal in circular dichroism (CD) spectroscopy, due to the excited states of the tryptophan pairs interacting to form a delocalized excited state, or exciton.^14^ Accordingly, such trpzip peptides have been used as models in folding experiments^15^ and simulations.^16–18^ Such studies on the folding mechanism of trpzip peptides have yielded two competing models: one termed the “zipping model” where the β-turn forms first, the hydrophobic core of the peptide then collapses, and then hydrogen bonds form; or the “hydrophobic-core-centric” mechanism, where the hydrophobic core collapses first with some hydrogen bonding present, before more of the hydrogen bonding forms as the configuration is finalized.^18^ These studies provide insight into the complex interplay between amino acid sequence and its resulting impacts on water solvation, hydrophobic collapse, and hydrogen bonding that influence secondary structure within peptides, proteins, and proteomimetics. Further demonstrating the utility of the trpzip sequence, such motifs have also served as scaffolds for antimicrobial, self-healing hydrogels^19^ and for antibody mimetics optimized via phage display for targeted cancer therapeutics.^20^ Additionally, they have been found to support copper and terbium ions as metalloprotein models.^21,22^

Here, we prepared the trpzip2 sequence as a polymerizable monomer to generate homopolymers and copolymers of peptide monomers. We found that regardless of polymer configuration and copolymer composition, the β-hairpin fold is significantly disrupted, with loss of β-sheet hydrogen bonding and a consistent drop in CD intensity, indicative of loss of the tryptophan zipper configuration upon polymerization. These results are further corroborated by infrared spectroscopy studies and are consistent with small-angle X-ray scattering (SAXS) confirming the globular collapse of the polymers and nuclear magnetic resonance (NMR) spectroscopy that likewise suggests loss of the hairpin configuration. To elucidate the impact of polymerization on the conformational stability of the trpzip2 β-hairpin at an atomistic level, we conducted extensive molecular dynamics simulations on both the monomeric and polymer-conjugated forms of the peptide. These simulations enabled a direct comparison of the folding landscapes in isolation versus within the context of the polymer. Consistent with previous structural studies, the monomer readily adopts a well-defined β-hairpin conformation, with a pronounced free-energy minimum corresponding to the experimentally determined native structure. In contrast, polymer conjugation substantially alters the folding behavior, favoring extended or misfolded states over the native fold. This shift arises from the collapse of the polymer into a compact globular architecture, which creates steric crowding and geometric constraints around the conjugated peptides. Although this confinement reduces the range of conformations accessible to each peptide, it preferentially restricts the backbone arrangements required to form the native hydrogen-bond network and correctly align the stabilizing tryptophan pairs. Consequently, the polymer-bound peptides exhibit reduced conformational heterogeneity, diminished native-contact formation, and lower structural similarity to the reference hairpin. These results show that the brush polymers do not merely restrict peptide motion but this architecture reshapes the accessible free-energy landscape in favor of conformations that are incompatible with the native β-hairpin, providing a mechanistic explanation for the experimentally observed loss of β-hairpin character upon polymerization. This study has important implications for recapitulating peptide folding within the proteomimetic brush polymer scaffold and suggests that such polymers may be useful where repression of β-sheet-like folding is desired.

## Experimental Section

### Circular Dichroism Spectroscopy

Circular dichroism (CD) spectra were obtained on a JASCO J-815 spectrometer equipped with a Peltier sample holder for temperature control in the Keck Biophysics Facility at Northwestern University. Samples were dissolved in aqueous buffer (50 mM sodium acetate buffer, pH 4) at approximately 100 μM with respect to peptide monomer concentration and placed into a 1 mm quartz cuvette (Hellma). Measurements were obtained using a scan speed of 10 nm/min, a 1 nm bandwidth, and 4 s response time. Background spectra of solvent were obtained prior to sample measurements and subtracted from the sample spectra using JASCO Spectra Analysis. Spectra were not smoothed. Sample concentration was corrected using calculated extinction coefficients^23^ and absorbance data measured at 205 nm. For variable temperature scans, the sample temperature was varied between 5 and 90 °C at a rate of 1 °C/min while monitoring ellipticity at 229 nm. At each end point of the temperature ramps (20 and 90 °C), full CD spectra were obtained.

## Infrared Spectroscopy

Samples of peptide monomers and polymers were dissolved in 10 mM HCl in H_2_O and lyophilized three times to remove trace trifluoroacetic acid that might interfere with analysis of the amide IR bands.^24^ After lyophilization, samples were dissolved in D_2_O (ca. 5 to 10 mg/mL) and then their Fourier transform infrared spectra were measured on a Thermo Scientific Nicolet iS50 infrared spectrometer. Solutions of samples (20 μL) were placed between two calcium fluoride plates (32 mm × 3 mm, Pike Technologies) with a 0.5 mm Teflon spacer, which were then put into a transmission cell and measured using 256 scans. Sample spectra were background-subtracted from a scan of D_2_O (20 μL) in the same setup.

## Molecular Dynamics Simulations

The structure of the trpzip2 monomer with norbornene dicarboximide linked to aminohexanoic acid (NorAha) was drawn using the Molefacture plugin in VMD^25^. The 15-mer (polymer) structures were also constructed in the same way. CHARMM36m^26^ was used to parametrize the standard protein residues, whereas CHARMM General Force Field^27^ was used for parameterization of NorAha and the interface bond/dihedrals. The peptide–NorAha linkage as well as polymer backbone bonds were implemented using patches in the PSFGEN package in VMD. Molecular dynamics simulations were performed using OpenMM^28^ at the distributed computing platform of Folding@home.^29^ Details of parameterization protocol as well as molecular dynamics simulation parameters are outlined in the Supporting Information.

## Markov State Modeling

The molecular dynamics trajectories were analyzed using Markov state models (MSMs). First, the trajectories were featurized into relevant physical descriptors using MDTraj^30^. These features were then projected into a lower-dimensional space using time-lagged independent component analysis (tICA). The resulting tICA coordinates were clustered using KMeans, and the cluster assignments were used to build the Markov state models. All steps involving tICA, clustering, and MSM construction were carried out using the Deeptime library^31^. Therefore, all free energy plots shown in Figures 6, 7, and 8 are weighted with MSM weights. The details of model construction and hyperparameter selection are provided in the Supporting Information (Section 7: Simulations), along with implied time-scale plots (**Figures S31–S33**) and Chapman– Kolmogorov tests (**Figures S34–S36**).

## Results and Discussion

### Design, synthesis, and characterization of peptide monomers and polymers

Several monomers were prepared to test various configurations and arrangements of β-hairpin structures for incorporation into brush polymers (**Figure 1**, **Table 1**). Firstly, the trpzip2 monomer peptide was prepared as a norbornene dicarboximide linked to aminohexanoic acid (NorAha) via solid-phase peptide synthesis at the N-terminal end (NorAha-SWTWENGKWTWK, “beta”; Figures S1-S4, S5 and S10); the trpzip2 sequence where asparagine (N6) was functionalized with Nor-diaminohexane (SWTWE**N**(NorImAmine)GKWTWK, “N-beta”; Figures S6 and S11); and finally, the trpzip2 sequence ending in norbornene carboxyamide linked to AEEA (NorAmOEG-SWTWENGKWTWK, “NorAmOEG-beta”; Figures S7 and S12) was prepared. Additionally, two monomers were prepared where one of the two tryptophan pairs was replaced with valine – specifically NorAha-S**W**T**V**ENGK**V**T**W**K (“WVVW-beta”; Figures S8 and S13) and NorAha-S**V**T**W**ENGK**V**T**W**K (“VWWV-beta”; Figures S9 and S14). These mutated sequences were synthesized to examine the effect of purposefully removing the tryptophan pairs on the resulting polymer properties.^32^ The successful synthesis and purity of these five monomers were confirmed by HPLC and ESI-MS (SI Section 3).

**Figure 1.**
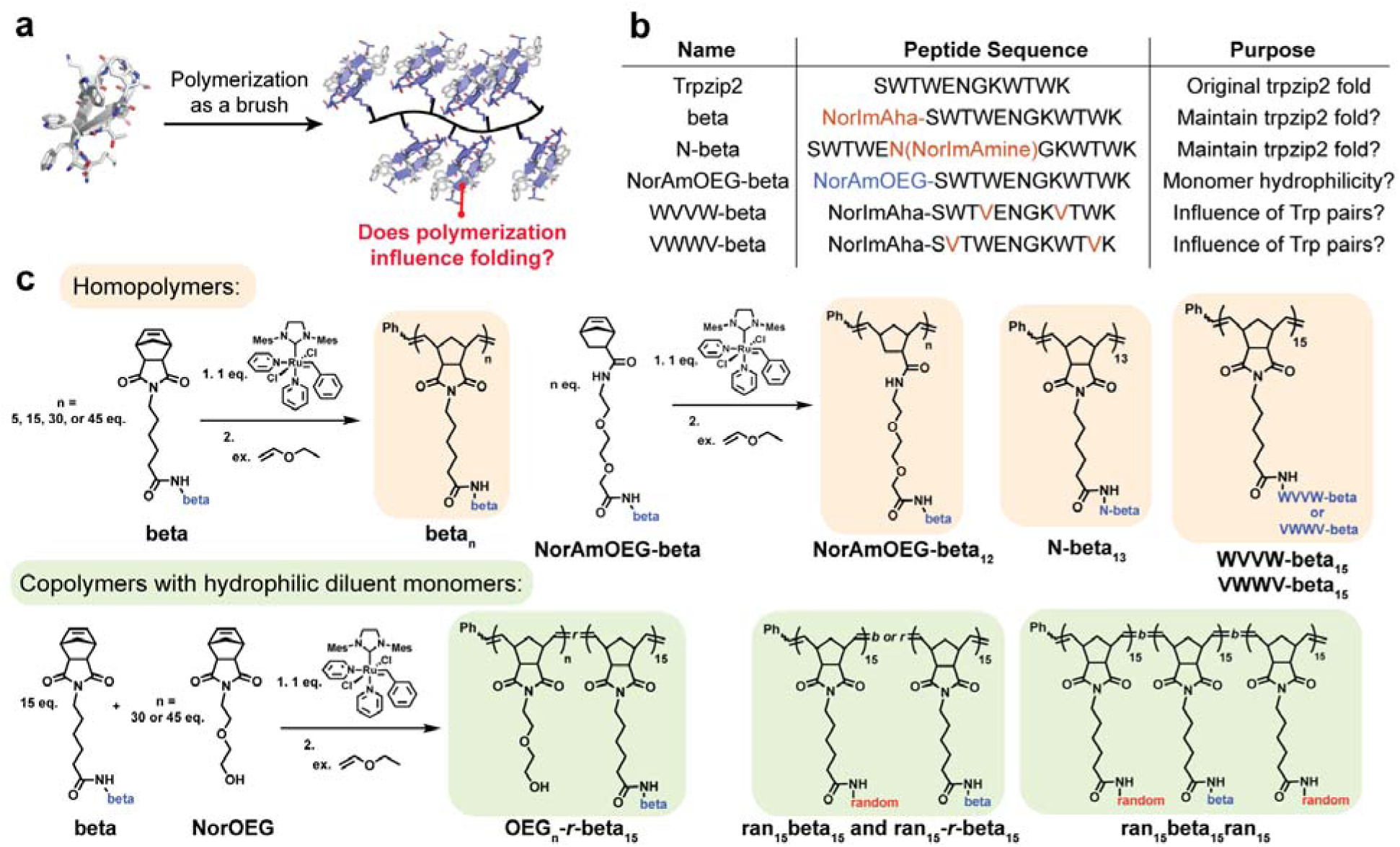
a) Peptide brush polymer design, with a hypothetical cartoon of a brush polymer of the trpzip2 β-hairpin peptide. **b)** Peptide sequences synthesized and polymerized, the acronyms for the sequences, and the questions driving their synthesis and study. **c)** Homopolymer and copolymer synthesis schemes.

**Table 1.**
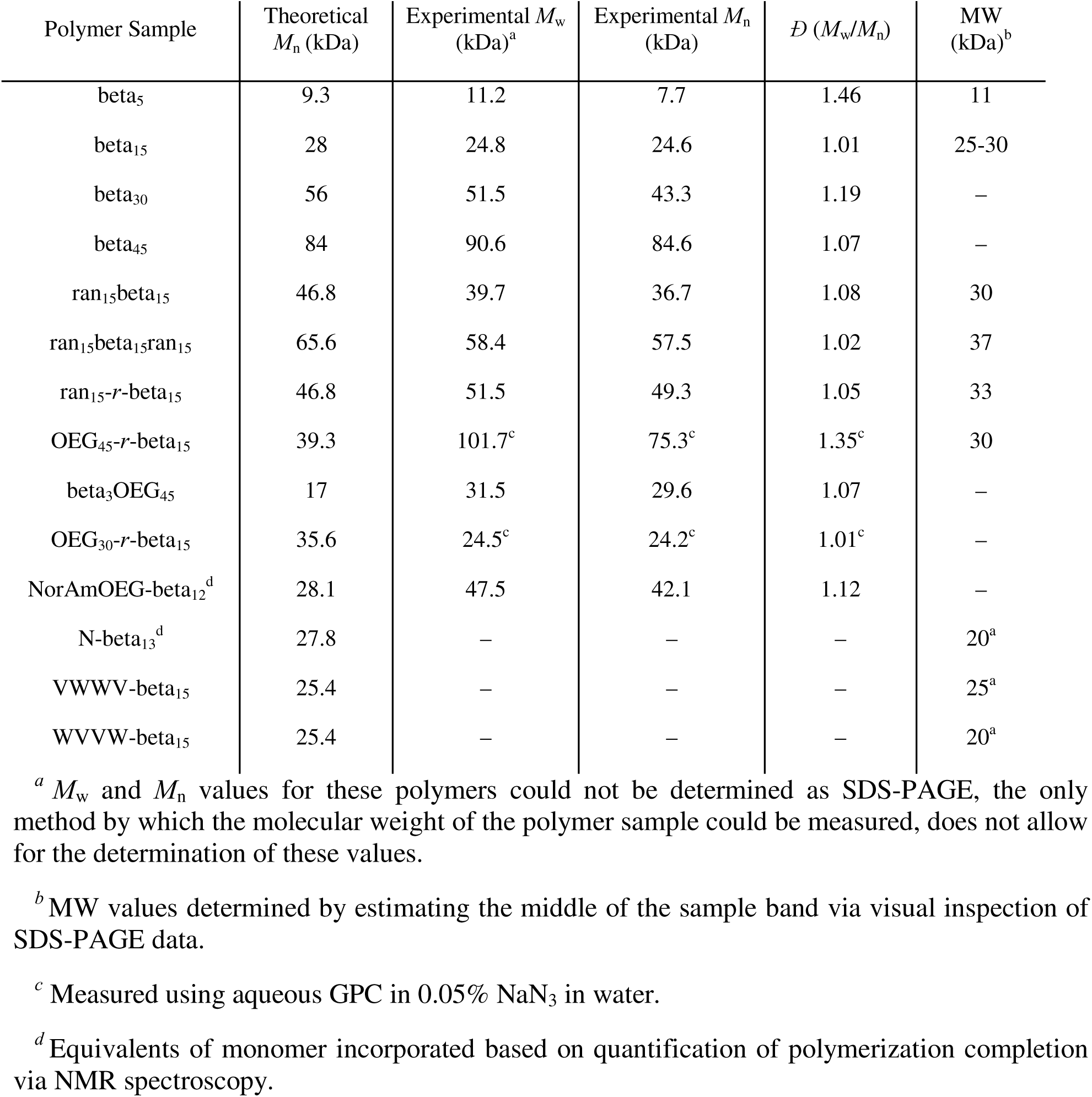
Molecular weight characterization of polymers

These monomers were then used to construct a library of polymers, including homopolymers of the beta monomer approximately 5, 15, 30, and 45 units in length (beta_5_, beta_15_, beta_30_, and beta_45_); 15-mer homopolymers of the N-beta, NorAmOEG-beta, WVVW-beta, and VWWV-beta (N-beta_13_, NorAmOEG-beta_12_, WVVW-beta_15_, and VWWV-beta_15_); copolymers of the beta monomer with a random coil sequence^12^ (ran_15_beta_15_, ran_15_beta_15_ran_15_, and ran_15_-*r*-beta_15_); and finally, copolymers of the beta monomer with an OEG-based monomer with no peptide (“NorOEG”; OEG_30_-*r*-beta_15_, OEG_45_-*r*-beta_15_, and beta_3_OEG_45_). Here, we sought to understand whether different linkages of the peptide to the polymer influenced trpzip2 folding (N-beta_13_) and whether monomer hydrophilicity impacted trpzip2 folding (NorAmOEG-beta_12_). We also aimed to test which tryptophan pairs of the trpzip2 sequence (WVVW-beta_15_ and VWWV-beta_15_) were more destabilized. Additionally, we hypothesized that copolymerization of the trpzip2 sequence with hydrophilic unfolded (where ran = NorImAhaGSGSGRGSGSGE; ran_15_beta_15_, ran_15_beta_15_ran_15_, and ran_15_-*r*-beta_15_) or unhindered monomers (OEG_30_-*r*-beta_15_, OEG_45_-*r*-beta_15_, and beta_3_OEG_45_) would allow the trpzip2 moiety to fold as for the unpolymerized peptide sequence. All polymerizations were conducted using ring-opening metathesis polymerization (ROMP) via a modified third-generation Grubbs’ catalyst, according to previously published protocols,^33–35^ while monitoring the polymerizations using nuclear magnetic resonance (NMR) spectroscopy (**Figures S15–S18**). Polymers were purified by chromatography and characterized by size-exclusion chromatography coupled with multi-angle light scattering (SEC-MALS) and sodium dodecyl sulfate-polyacrylamide gel electrophoresis (SDS-PAGE) to confirm that all samples had the targeted molecular weight (Table 1 and SI Section 4, **Figures S19-20**). We note that although we can resolve monomer features with NMR spectroscopy, particularly using 2D NMR spectroscopy, the polymers in aqueous buffer had broadened spectra that did not allow for further analysis.

## CD spectroscopy reveals loss of Trp–Trp exciton upon polymerization

Folding of the trpzip2 sequence before and after polymerization, could be examined using circular dichroism (CD) and infrared (IR) spectroscopy (**Figure 2; SI** Sections 5 and 6). This class of peptides is an appealing test case because studies have demonstrated that modifications to the originally reported trpzip sequences can influence their stability, related to turn sequence, charged amino acids, and tryptophan residues contributing to the stability of the β-hairpin fold in trpzip peptides.^14,32,36,37^ Critically, the characteristic exciton signal in the CD spectra of tryptophan zippers is influenced by the strength of the interactions between the tryptophan pairs, and substitution for other aromatic or aliphatic amino acids causes significant drops in the CD intensity as well as weakening of the fold stability.^14,32,36,37^ It is found that mutating the tryptophan pair closest to the β-turn has an impact on turn stability, indicating that these amino acids might have a strong influence. In addition to affecting the intensity of the CD signal, such mutations caused red-shifting of the maxima of the peaks in the CD spectra (e.g., from 226.8 nm to 227.4 nm) and calculations suggested a loss of intramolecular hydrogen bonding with increasing substitutions.

**Figure 2.**
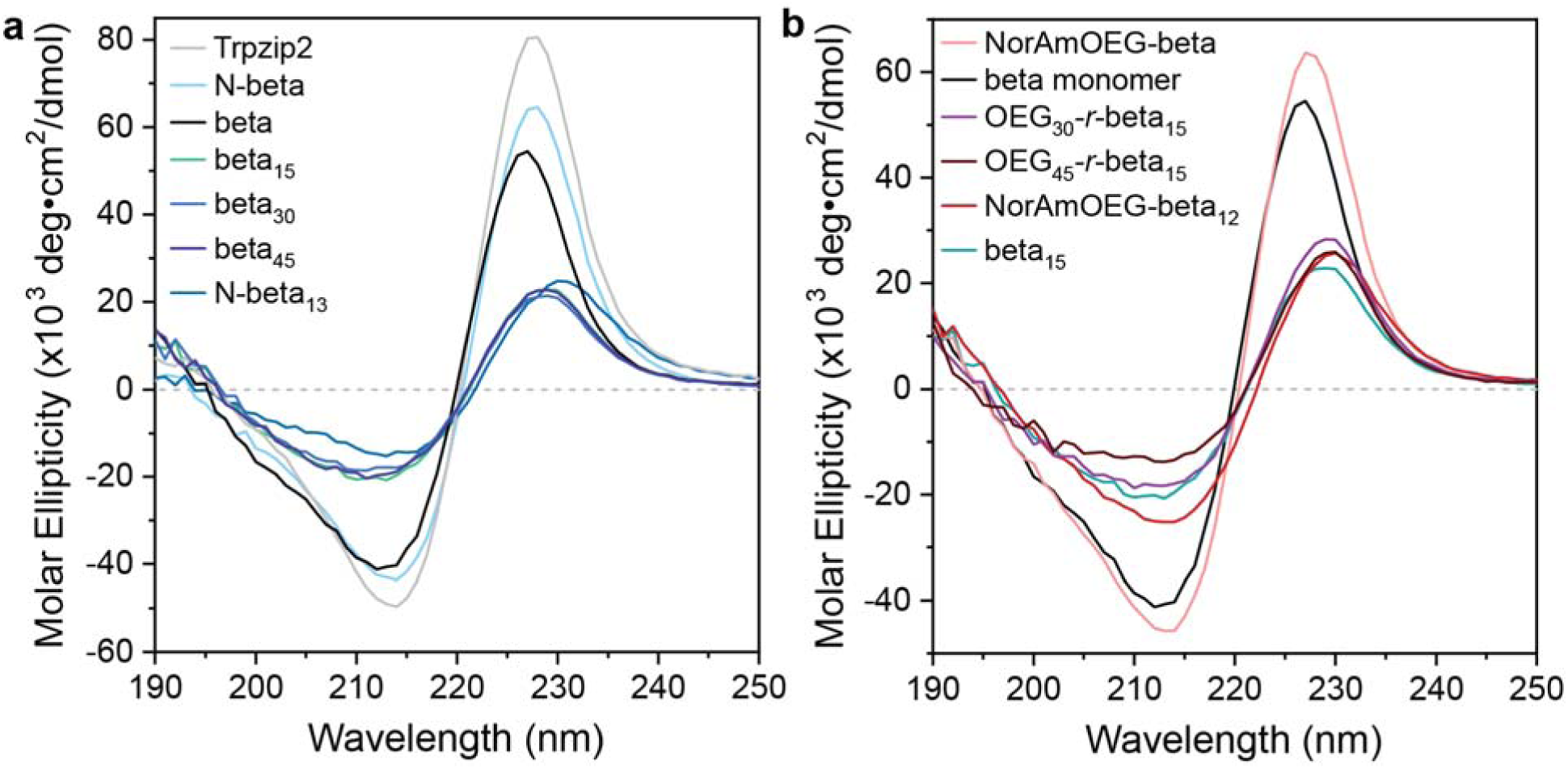
a) CD spectra of the original trpzip2 sequence (light gray trace), the N-beta monomer (light blue trace), the beta monomer (black trace), beta_15_ (teal trace), beta_30_ (blue trace), beta_45_ (navy blue trace), and N-beta_13_ (dark teal trace). **b)** CD spectra of the co-polymers OEG_30_-*r*-beta_15_ (light purple line) and OEG_45_-*r*-beta_15_ (dark purple line), as well as the beta monomer (black line), the NorAmOEG-beta monomer (pink line), the beta_15_ polymer (green line), and the NorAmOEG-beta_12_ polymer (red line).

All peptide monomers with the exact trpzip2 sequence (beta, N-beta, and NorAmOEG-beta) reproduce the original trpzip2 CD spectra, indicating that the presence of the norbornene linker units does not perturb the native trpzip2 fold (**Figure 2**). In particular, the features are present in the same positions as the published work, including the additional feature at approximately 300 nm, from tryptophan absorption. However, in the CD spectrum of N-beta, there is a slight redshift in the maximum position from 227 nm (from the original trpzip2 peptide) to 228 nm (**Figure 2a**).

All polymers reveal a preservation of features that are found in the CD spectra of their respective monomers but show a dramatic drop in CD intensity (**Figure 2**, **Figures S23-26**). For almost all polymers, this drop corresponds to approximately 40% of the original beta monomer intensity, when adjusted for molar ellipticity and trpzip2 sequence percentage (in the case of copolymers). The polymer intensity is greater than that observed of a typical unfolded control, the equimolar combination of the half peptides SWTWEN and GKWTWK (**Figure S25**).^13^ This drop is particularly noteworthy in the case of the beta homopolymers, which almost overlay with each other, indicating that the drop in intensity might be intrinsic to the polymer (**Figure 2a, Figure S23**). In other words, every monomer on the polymer brush appears to experience the same destabilization of hairpin folding, irrespective of polymer length, as polymers approximately 5 units in length experience the same degree of folding as polymers approximately 45 units in length. Similar drops in intensity are observed in the copolymers with the random coil sequence as well as, surprisingly, copolymers with the OEG-based monomer, when the spectra intensity is scaled for percentage of trpzip2 content (**Figure 2b, Figure S26**). In the random coil-based copolymers, a clear increase in random coil content is observed that is consistent with the increase in random coil monomer content, as evidenced by an increase in the growth of the negative peak at 200 nm, which is particularly prominent in the ran_15_beta_15_ran_15_ polymer. The loss of CD intensity of samples of the beta monomer randomly copolymerized with the Nor-OEG monomer is striking, as it supports that steric hindrance alone does not account for the loss in CD intensity. Additionally, the results from the NorOEG copolymers and the NorAmOEG-beta_12_ polymer suggest that relatively hydrophilic polymer environments are also not able to recover trpzip2 folding.

Additionally, many spectra of polymerized trpzip2 contain a redshift in peak positions. For example, in beta_15_, the peak maximum shifts from 227 to 229 nm; in N-beta_15_, the maximum shifts from 228 to 230 nm. These shifts appear to be related to changes in the π–π interactions with the Trp pairs and also potentially weakening of the interaction due to increased distances between the two ring systems.^32^ We note that while variable-temperature melting curves of beta_15_ relative to the beta monomer were obtained using CD spectroscopy, only minor changes were observed between the two samples using this method (**Figure S27**).

## IR spectroscopy confirms loss of features found in original trpzip2

Infrared spectroscopy provides additional insight into secondary structure, complementary to CD spectroscopy, and has been used on trpzip2-based peptides to further elucidate their structural features.^32,37–41^ Our studies of monomer and polymer samples using Fourier transform infrared (FT-IR) spectroscopy reveal that the beta monomer closely matches the data of the original trpzip2 peptide, with peak positions at 1633.8 and 1675 cm^−1^. These peak positions match those identified to correspond to β-sheet content, specifically 1624–1637 cm^−1^ and 1675 cm^−1^ in D_2_O (**Figure 3a**).^42^ Upon polymerization to a 15-mer, the peak at 1675 cm^−1^ develops a prominent, blueshifted shoulder at 1680–1690 cm^−1^, which indicates an increase in β-turn content. This shift is consistent with both the CD data and simulation data described below, which indicates a significant disruption of the noncovalent interactions between tryptophan pairs and a resulting destabilization of β-sheet-like hydrogen bonding.

**Figure 3.**
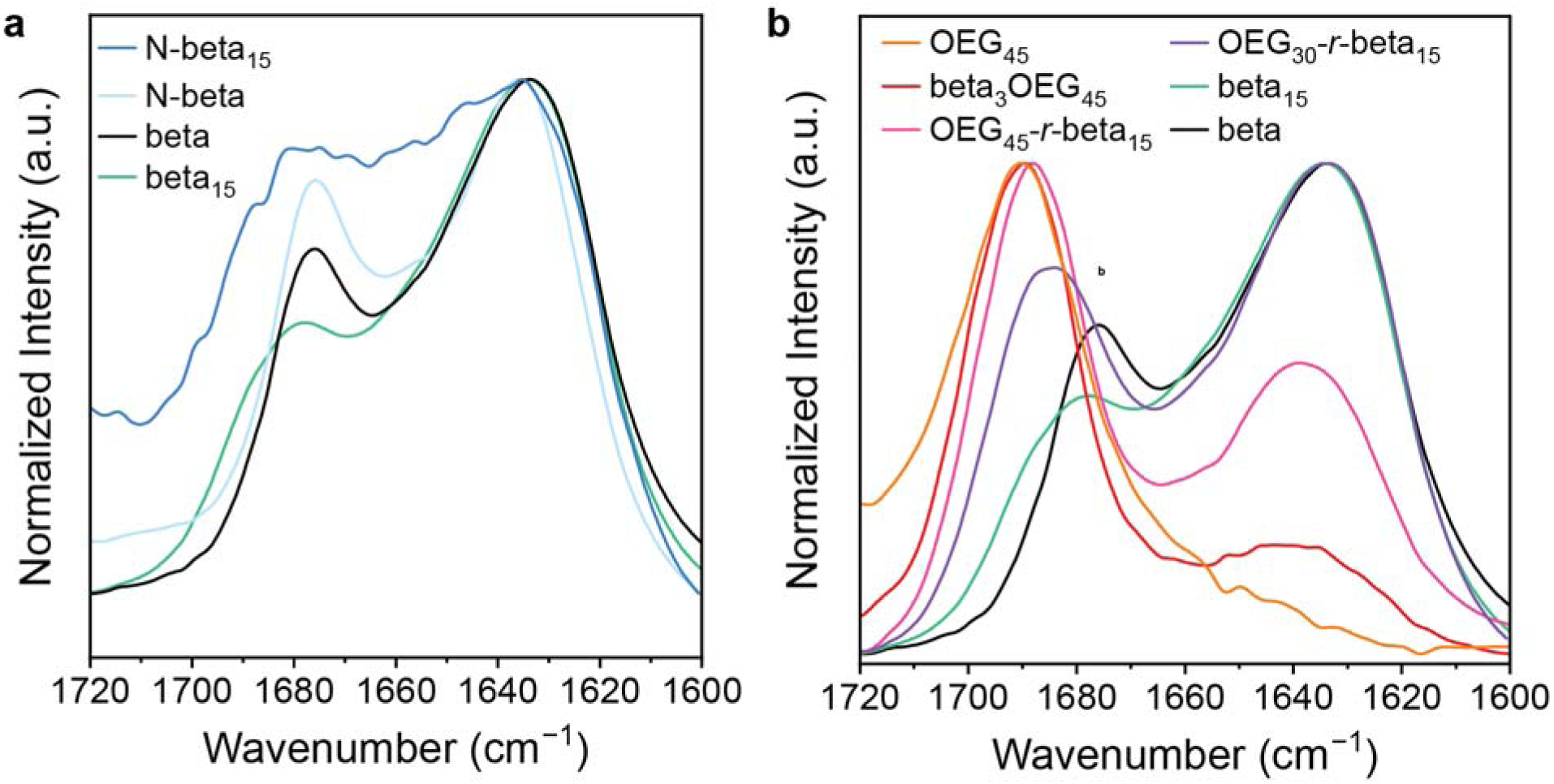
a) Transmission FT-IR spectra of the monomers beta (black line) and N-beta (light blue line), as well as the polymers N-beta_15_ (blue line) and beta_15_ (green line). **b**) Transmission FT-IR spectra of NorOEG_45_ (orange line), beta_3_OEG_45_ (red line), OEG_45_-*r*-beta_15_ (pink line), OEG_30_-*r*-beta_15_ (purple line), beta_15_ (green line), and the beta monomer (black line) for comparison.

In copolymers containing the random coil monomer, there is a clear shift in the maximum at 1633.8 to 1646 cm^−1^ (ran_15_beta_15_ran_15_; **Figure S29**), which is in line with the greater amount of random coil monomer content attributed to appear at approximately 1645 cm^−1^.^42^ The copolymers retain the left peak maximum at 1678 cm^−1^ that is observed in beta_15_, as well as a shoulder at 1685 cm^−1^. In contrast, copolymers with the OEG-based monomer (**Figure 3b**) have a decrease in relative intensities of β-sheet features at 1633 and 1675 cm^−1^, with a contribution from the NorOEG monomer that is observed at 1689 cm^−1^ that grows in with increasing NorOEG content. As the amount of beta monomer content drops, the rightmost peak maximum shifts from 1633.8 to 1640 cm^−1^ (beta_3_OEG_45_), indicating a likely increase in random coil content. This shift is in contrast to that observed with NorAmOEG-beta, which has its maximum shift from 1637 to 1633 cm^−1^ upon polymerization to NorAmOEG-beta_12_ (**Figure S30**). The N-beta monomer and N-beta_15_ sample also show development of a shoulder at approximately 1690 cm^−1^ that appears in the polymerized sample, and a shift from 1633.8 cm^−1^ in the monomer to 1640 cm^−1^ in the polymer, indicative of a loss of the β-hairpin fold of the monomer upon polymerization (**Figure 3a**). Overall, the FTIR spectra confirm the loss of β-sheet hydrogen bonding upon polymerization of the trpzip2 sequence into peptide brush polymers.

## Mutations of trpzip2 sequence confirm destabilization of β-hairpin fold

Prior work on the trpzip2 sequence showed that mutation of the tryptophan pairs allowed for additional insight into the stability of the β-hairpin fold. In particular, mutation of the tryptophan residues closest to the β-turn valines (W4 and W9) have been shown to contribute to destabilization of the hairpin fold, as assessed by CD and FTIR spectroscopy.^32^ In our observations of monomers and polymers of similarly altered trpzip2 sequences, we have also found that mutating the tryptophan pair structures weakens the trpzip2 fold stability to an even greater extent. Notably, both WVVW-beta and VWWV-beta monomers show significant drops in CD spectra intensity relative to published data on comparable sequences without norbornene (**Figure 4a**), as well as significant redshifts in their peak maxima relative to the parent trpzip2 sequence (228 and 229 nm for VWWV-beta and WVVW-beta respectively, relative to the beta monomer, 227 nm; **Figure S24**).^32^ We hypothesize that this may be due to additional destabilizing effects from the hydrophobic norbornene linker on the monomer, which, while not influencing the native trpzip2 fold, may preferentially destabilize the mutated sequence by altering the hydrophobicity that helps to favor the pairing of the remaining tryptophans. This drop in the CD intensity of the WVVW-beta and VWWV-beta monomers persists even when the samples are cooled to 5 °C (**Figure S28**).

**Figure 4.**
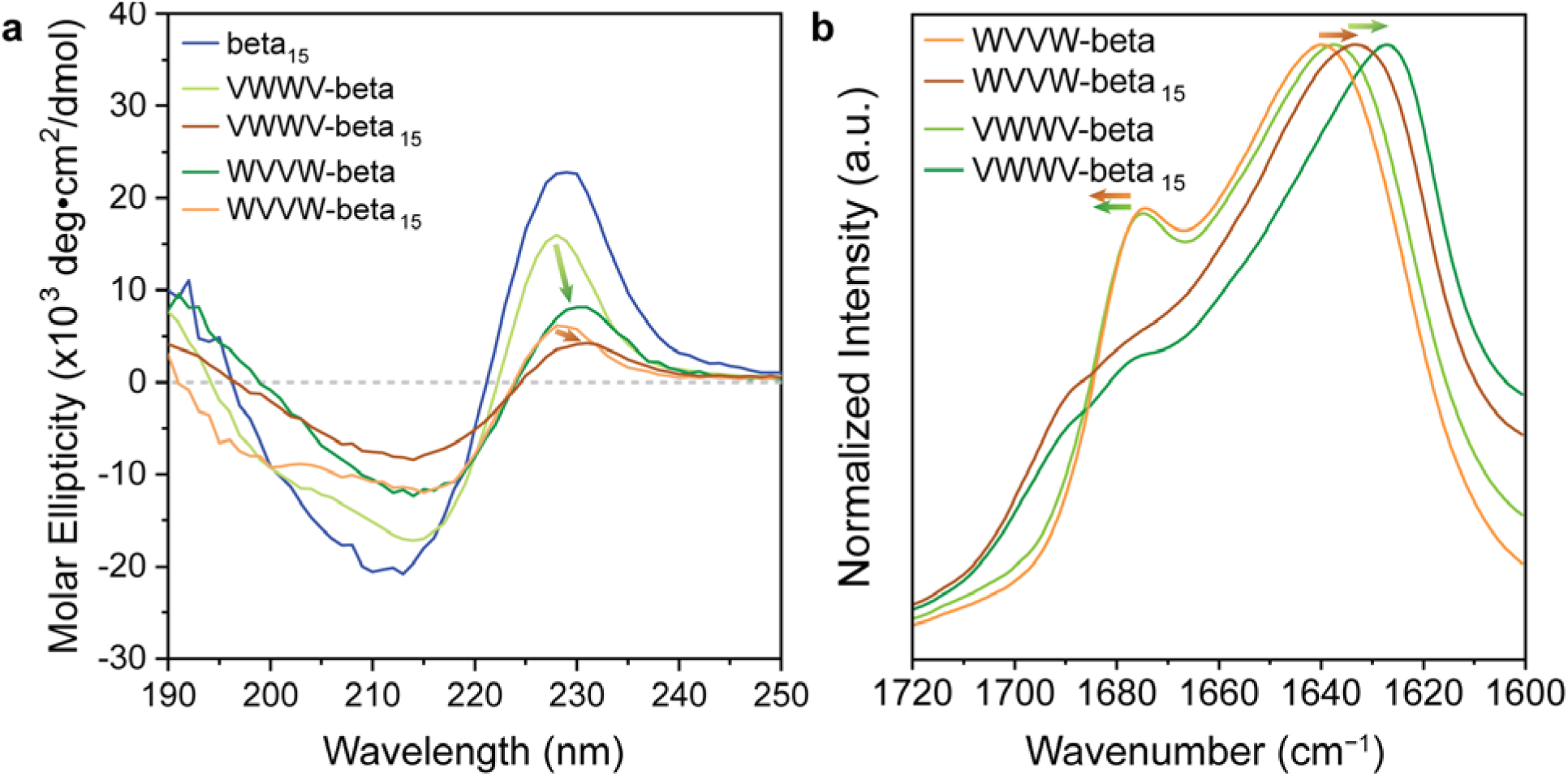
a) CD spectra of the VWWV-beta monomer (light green trace), VWWV-beta_15_ polymer (green trace), WVVW-beta monomer (orange trace), WVVW-beta_15_ polymer (brown trace), and beta_15_ polymer (blue trace). Colored arrows indicate the transformation of the CD peak maxima upon polymerization. **b)** Transmission FT-IR spectra of the VWWV-beta monomer (light green trace), VWWV-beta_15_ polymer (green trace), WVVW-beta monomer (orange trace), and WVVW-beta_15_ polymer (brown trace). Colored arrows indicate the transformation of the FT-IR peak maxima upon polymerization.

Likewise, polymers of the mutated monomers display almost a 50% drop in CD intensity as well as red-shifts in the position of the maximum relative to the spectra of the monomers (228 to 230 nm for VWWV-beta to VWWV-beta_15_, and 229 to 231 nm for WVVW-beta to WVVW-beta_15_; **Figure 4a**). This combination of effects indicates that the β-hairpin folds in the mutated polymers are significantly destabilized and reveals that while the original sequence shows loss of the β-hairpin fold, it is not a complete loss. It is also noteworthy that the VWWV-beta system, which preserves a key tryptophan pair near the β-turn, shows greater evidence of folding in both the monomer and polymer than the WVVW-beta sequence, which mutates this tryptophan pair. This result is consistent with previous reports showing the particular importance of these two tryptophan positions to the stability of the hairpin fold.^32,43^

In comparison to the CD results, infrared spectra of the VWWV-beta and WVVW-beta monomers display similar spectra to that of the beta monomer, with prominent bands at 1640 and 1675 cm^−1^ for WVVW-beta, and 1637 and 1675 cm^−1^ for VWWV-beta (**Figure 4b**). In the polymers, there are shifts in these bands to 1633, 1679, and 1688 cm^−1^ for WVVW-beta_15_; and 1627, 1679, and 1688 cm^−1^ for VWWV-beta_15_. These trends generally match those observed with the beta_15_ polymer, with an increase in apparent β-turn content and loss of some β-sheet character. We note one key difference between the two sets of spectra – a smaller shift in the peak position of 1640 to 1633 cm^−1^ is observed for the WVVW-beta monomer to the polymer WVVW-beta_15_, in contrast to the greater shift observed for VWWV-beta to VWWV-beta_15_, 1637 to 1627 cm^−1^. This increase in the shift of the IR band may be related to the fact that the WVVW-beta monomer shows greater initial destabilization of the hairpin fold than the VWWV-beta monomer.

## SAXS data corroborate globular collapse in aqueous solution

Small-angle X-ray scattering data provide insight into the solution-phase conformation of polymers at the nanoscale.^5,44,45^ In particular, we were curious whether these polymers collapse into globules, or whether the β-hairpin folds would resist globular collapse and cause the polymer to remain extended in solution. To explore this question, SAXS data were collected on the polymer samples in aqueous buffer (**Figures 5** **and S38-40, SI** Section 8). In almost all cases, data consistent with monodisperse, globular systems were observed (**Figure 5**). Performing Guinier analyses yielded radii of gyration (*R*_g_) ranging from 2.4 to 3.0 nm (**Table S1**), significantly smaller than the value of *R*_g_ anticipated for a fully extended polymer backbone (∼4.5 nm for a 15-mer). These values indicate that the polymers collapse in solution, consistent with the normalized Kratky plots that demonstrate that the samples approximate the features of compact spheres. The Kratky plots contain bell-shaped curves centered close to values predicted for compact spheres (indicated by gray dashed lines at values of *qR*_g_ = 1.73205 and (*qR*_g_)^2^*I*_(q)_/*I*_0_ = 1.104; **Figure 5b** and **Figure S41**). This qualitative assessment is corroborated by power law and Guinier–Porod fits performed on the data that yield parameters consistent with compact globular objects (**Tables S2 and S3; Figures S42–S43**). Specifically, through the Guinier–Porod fit, all data have been fit to values of the dimension variable *s* that are 0.26 or less.^46,47^ Additionally, the Porod exponent is generally greater than 3. Data also showed reasonable fits to the flexible cylinder model (**Table S4, Figures S44–S45**), with radii 2 nm or less in diameter and lengths that approximately increase as the polymer chain is extended, similar to those obtained in other peptide brush polymers.^12^

**Figure 5.**
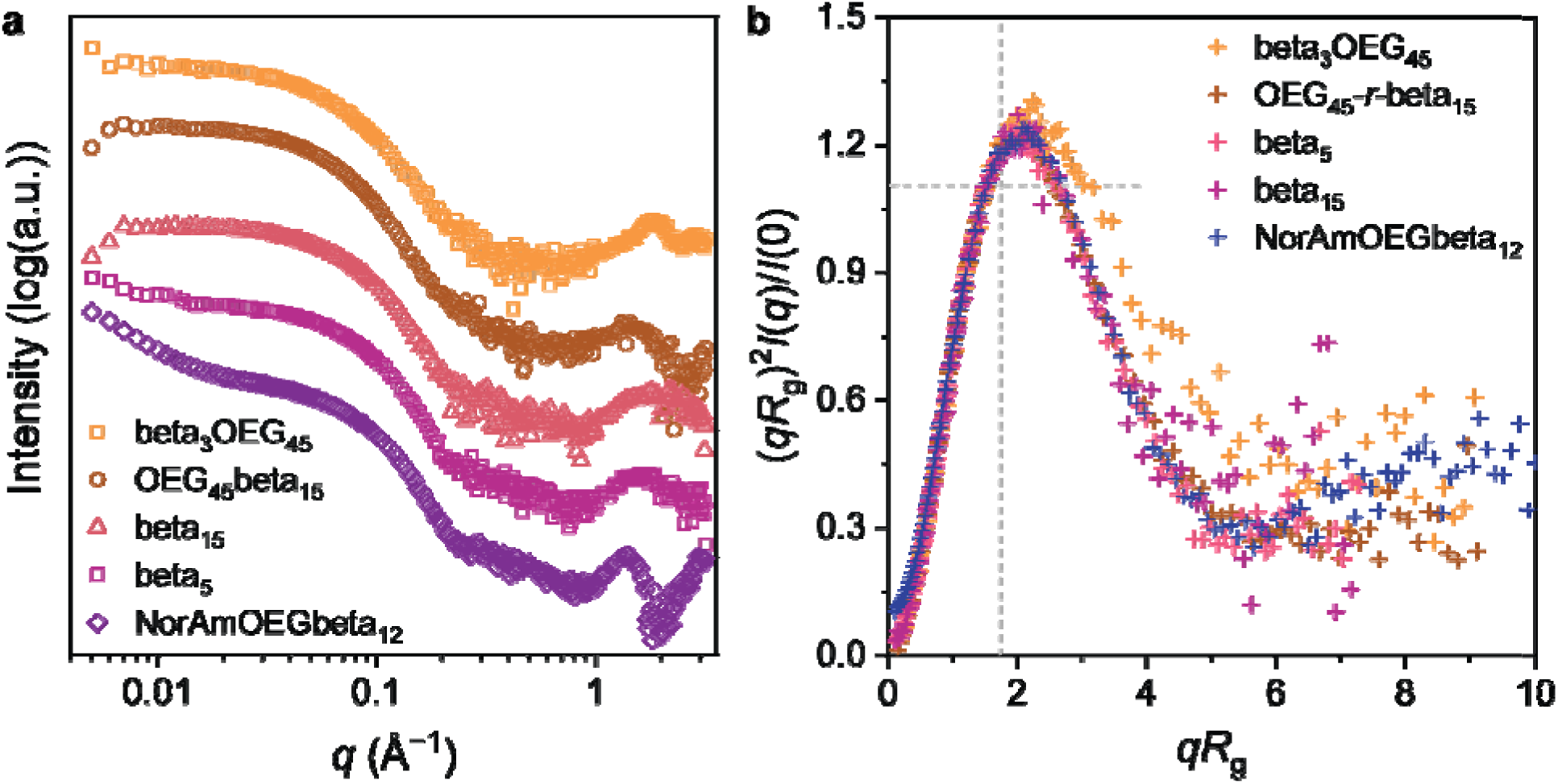
a) Raw SAXS data, plotted in double-logarithmic scale, of beta_3_OEG_45_ (yellow symbols), OEG_45_-r-beta_15_ (orange symbols), beta_15_ (red symbols), beta_5_ (magenta symbols), and NorAmOEG-beta_12_ (purple symbols). Patterns are offset horizontally for clarity. The wavelength of measurement is listed in Table S1. **b)** Normalized Kratky plots of beta_3_OEG_45_ (yellow dashes), OEG_45_-r-beta_15_ (orange dashes), beta_15_ (red dashes), beta_5_ (magenta dashes), and NorAmOEG-beta_12_ (purple dashes). Values for an ideal compact sphere are indicated by gray dashed lines at q*R*_g_ = 1.73205 and (*qR*_g_)^2^*I*_(q)_/*I*_0_ = 1.104.

The observed globular collapse in trpzip2-based peptide brush polymers is consistent with observations in similar peptide brush polymer systems.^5,12^ Previous results obtained with brush polymers containing α-helices showed that globular collapse did not impede helical folding. However, with these systems, it is observed that the fold of the β-hairpin monomer is significantly disrupted when globular conformations are observed. Additionally, we have performed fits of the scattering data to molecular dynamics simulation data, which are described further below and which further validate that the globular collapse of the polymer is observed alongside loss of the native trpzip2 folding pattern.

## In silico investigation of trpzip2 monomer illustrates the sustained stability of the hairpin conformation

Large-scale unbiased molecular dynamics simulations were performed for the trpzip2 monomer unit to elucidate the folding mechanism and to serve as a benchmark for comparison to the polymer system (**Figure 6**). The cumulative simulation time for the monomer simulations was 1.736 ms, with a representative replicate in the simulation box shown (**Figure 6a**). To analyze the high-dimensional trajectories, the simulation data were featurized into the following features: root-mean-square deviation of the peptide structure from the NMR structure,^13^ fraction of native contacts,^48^ Cα distances for all residues separated by two or more neighbors, native hydrogen bond distances,^49^ and the sum of radii of gyration for the two tryptophan pairs. All of these features quantify the extent of folding of the peptide with reference to the published NMR structure and the degree of compactness achieved by the native hydrogen bonds, as well as the aromatic interactions from the tryptophan zippers. For further dimensionality reduction, these structural features were used as input to time-lagged independent component analysis (tICA).^50^ This technique, tICA, is a dimensionality reduction technique commonly applied to molecular dynamics simulations^51^ to identify the slowest, most relevant collective motions in the system. By projecting the high-dimensional feature space onto a lower-dimensional subspace defined by these slowest dynamical modes, tICA enables a more interpretable representation of the peptide’s conformational landscape, emphasizing transitions relevant to folding and stability.

The projection of the simulation data is illustrated as a free-energy landscape governed by distinct metastable basins, defining both on- and off-pathway conformational states (**Figure 6b**). Interestingly, the landscape also captures a high-energy, off-pathway intermediate characterized by a helical conformation (**Figure 6c**). The global free energy minimum corresponds to a highly stable, completely folded conformation (**Figure 6d**). This stability is driven primarily by the precise alignment and aromatic stacking of the tryptophan pairs, which creates a well-defined hydrophobic core necessary to stabilize the structure. The transition from the unfolded coil to the native folded state proceeds through a distinct on-pathway intermediate conformation (**Figure 6e**). This intermediate state adopts a semi-folded topology stabilized by partial backbone hydrogen bonding networks. However, this lacks the alignment and aromatic stacking of the critical tryptophan pairs. The absence of these hydrophobic interactions renders the intermediate structure unstable, and it transitions between the completely folded and unfolded conformations. Previous experimental and computational works^17,19,49^ have identified a similar folding pathway. Conversely, a broader, higher-energy basin represents a fully unfolded, disordered coiled conformation (**Figure 6f**).

## In silico investigation of polymer system confirms diminishing prevalence of folded hairpin conformation

To assess the conformational space of the polymer-bound trpzip2 units and find the molecular origins of the secondary structure variations observed experimentally, we performed extensive molecular dynamics (MD) simulations of a 15-mer polymer system yielding an aggregate simulation time of 100 μs. Visual inspection of the trajectories reveals a clear sequence of events: upon initiation, the extended polymer chain undergoes a rapid structural collapse, ultimately transitioning into a dense, globular shape (**Figure 7a**). This globular collapse^5^ is driven by a cooperative collapse of the polymer backbone and sidechains, a physical phenomenon strongly correlated with the exceptional proteolytic resistance characteristic of these peptide brush architectures.^1^ To quantitatively track this compaction, we calculated the surface area of the 3D convex hull^52^ encompassing the coordinates of all polymer atoms over the course of the trajectory. A temporal reduction in this metric signifies a transition from extended, solvent-exposed states to a compact, globular state. The calculated mean convex hull area across 100 independent replicates decreases steadily over the simulation timeline, confirming a robust and reproducible collapse into a highly compact geometry (**Figure 7b)**.

**Figure 6.**
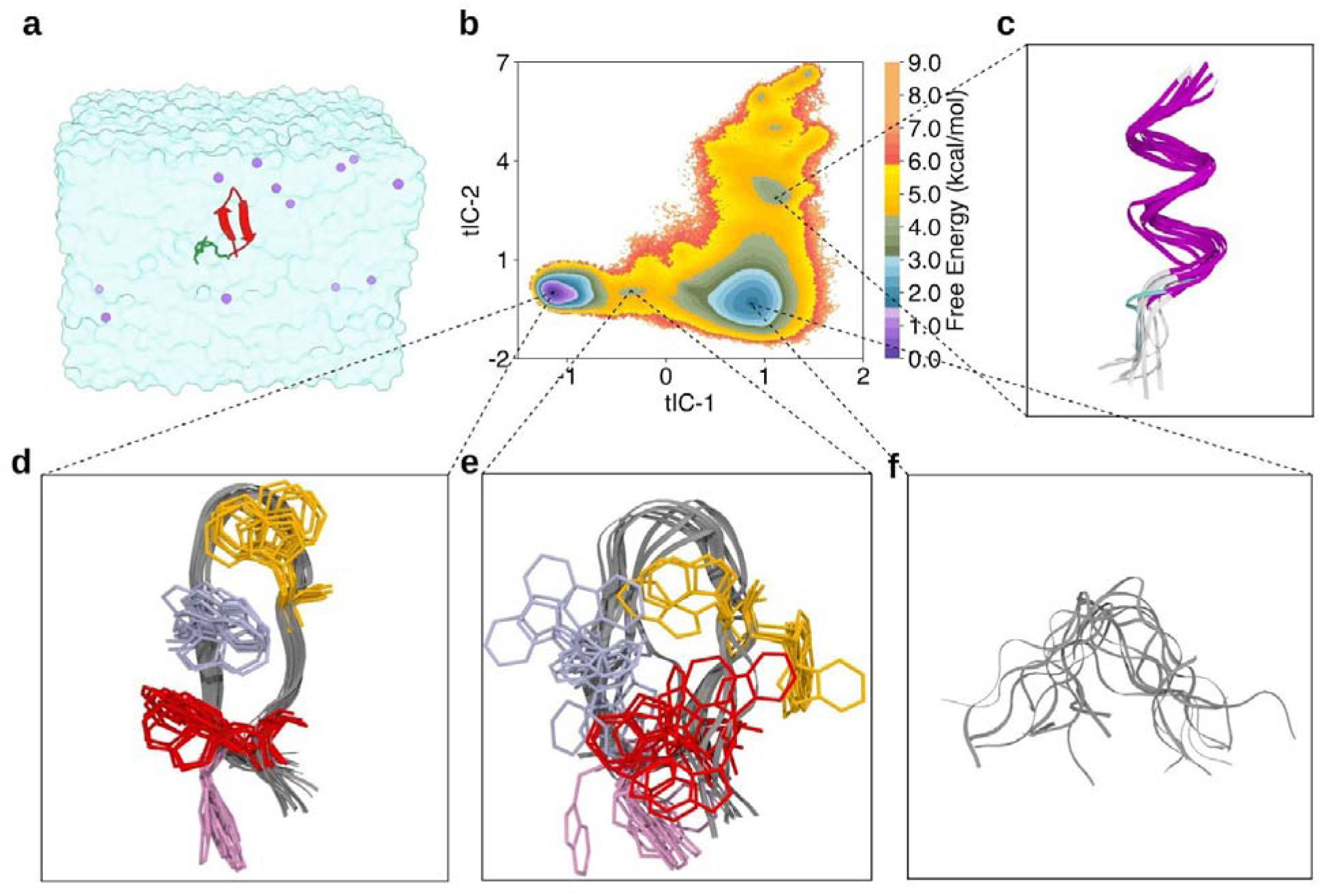
Simulations of the β-monomer. **(a)** A representative replicate within the simulation box, where the NorAha unit is shown in green and the trpzip2 structure is represented by the red ribbon diagram. **(b)** The free energy landscape of the monomer system projected onto tICA dimensions. **(c)** An off-pathway α-helical conformation of the β-monomer. **(d)** The lowest-energy conformation of the β-monomer closely matches the published trpzip2 structure, with the peptide backbone shown in gray and the tryptophan sidechains occupying highly consistent positions indicated by yellow, light purple, red, and pink lines. **(e)** An intermediate-energy conformation exhibiting a more open turn structure and highly disordered tryptophan orientations. **(f)** Unfolded conformation displaying no defined secondary structure.

**Figure 7.**
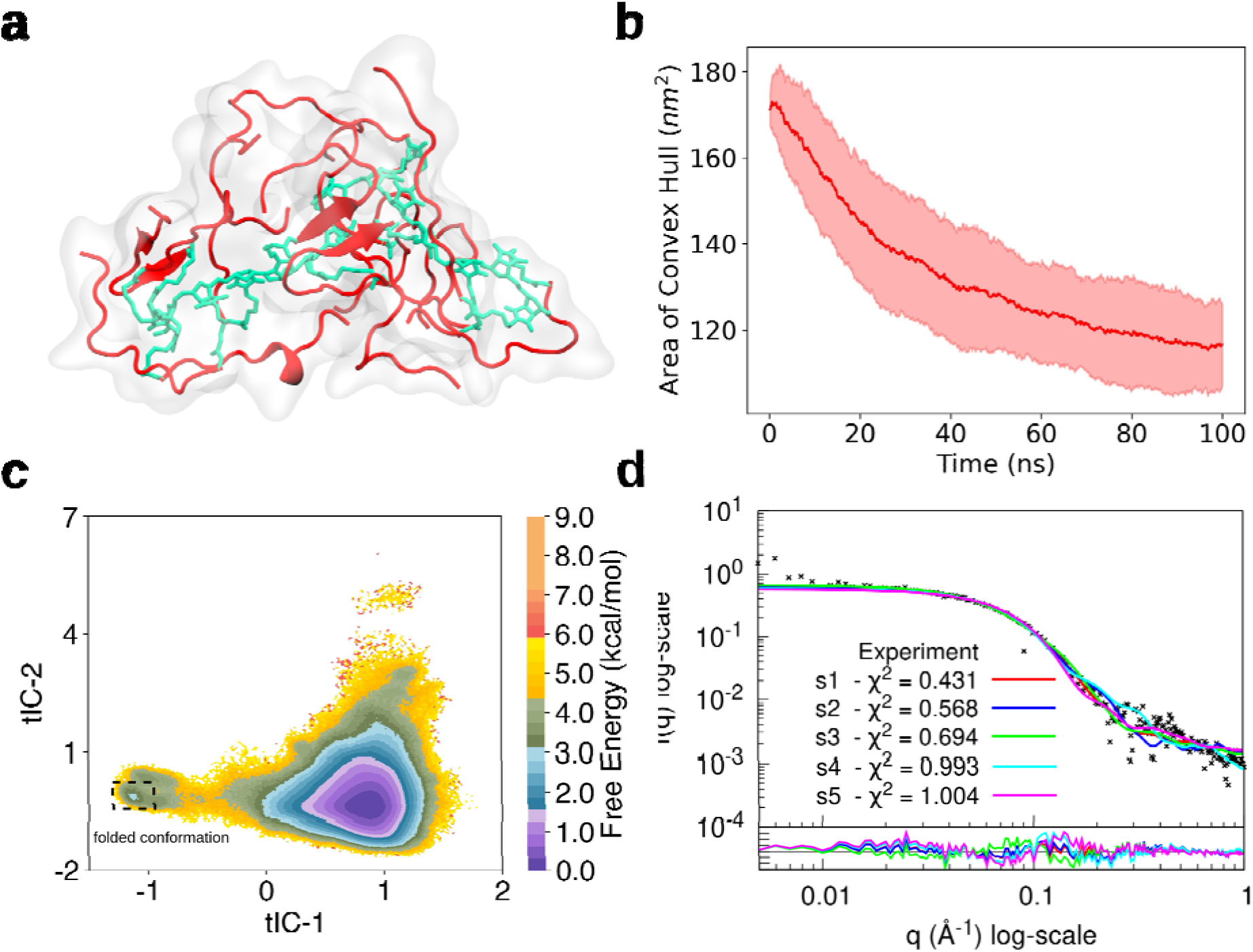
Simulations of the Ser-linked β polymer. **(a)** A representative snapshot from MD simulations of the 15-mer polymer system, illustrating the globular collapse of the polymer structure. The polymer backbone is shown in teal, while the linked peptides are highlighted in red. **(b)** The free energy landscape of the 15-mer system projected onto tICA dimensions, which were fitted using monomer simulation data. **(c)** Comparison between the experimental SAXS profile and computationally calculated SAXS profiles for five randomly selected polymer structures from the MD simulations. **(d)** Evidence of globular collapse represented by a monotonic decrease in the area of the convex hull over time. The solid red trace indicates mean value of area from 100 randomly selected trajectories, whereas the shaded area represents data values within one standard deviation.

To compare the folding dynamics of the polymer-mounted peptides against the isolated monomer, each trpzip2 unit was treated as an independent instance, effectively extending the analyzed sampling dataset to 1.5 ms (15 monomer units × 100 μs). Applying the same geometric feature set used for the monomer, we projected the polymer trajectories onto the monomer-fitted time-lagged independent component analysis (tICA) space. The resulting free energy landscape exhibits significant topological differences from the monomeric baseline (**Figure 7c**). Notably, the fully folded native conformation is no longer the global thermodynamic minimum; instead, the lowest free energy state shifts decidedly toward the unfolded ensemble. This behavior likely results from steric crowding arising from the globular collapse (**Figure 7b**) of the polymer, which imposes geometric constraints that disfavor formation of the ordered β-hairpin structure. This prevents the native hydrogen bonds supported by the folded conformation from forming in the polymer configuration (**Figure S37**). In addition, the free energy barriers between the folded and unfolded states remain relatively unchanged. It is also interesting to observe that the polymer landscape exhibits a restricted sampling profile and lowered conformational entropy compared to the monomer baseline (**Figure 6b**).

We also performed a comparison of molecular dynamics (MD) ensemble with experimental structural data; for which we randomly sampled MD snapshots and computed SAXS profiles using a command-line version of FoXS.^53^ These computational profiles were then compared to the experimental SAXS profile of the 15-mer polymer (**Figure 7d**). The computed and experimental profiles display strong agreement, particularly within the intermediate scattering vector range, confirming that the simulated ensembles accurately capture the solution-state polymer dimensions.

For investigating the mechanistic drivers of this structural shift, we evaluated the free energy landscapes using explicit physical descriptors. The radius of gyration of the core tryptophan pairs (*R_g_*), the cumulative distance of the five native hydrogen bonds (*L*), the fraction of native contacts (*q*), and the backbone root-mean-square deviation (RMSD) relative to the reference NMR structure were computed (**Figure 8**). For the monomer system, the projection across *R_g_* and *L* maps a deep, highly localized free energy minimum corresponding to the folded state at low *R_g_* and low *L* values, underscoring the stabilizing role of the native hydrogen bond network (**Figure 8a**). Conversely, the landscape for the polymer-bound units shows a thermodynamic inversion, with the global minimum shifting to high *L* and *R_g_*coordinates, indicative of an unfolded or highly extended ensemble (**Figure 8c**). This inversion occurs alongside a noticeable reduction in the free energy barrier separating the states. We note that in these plots the dashed line represents *L* = 17.5 Å, which is the upper limit for the value of *L*, when all five hydrogen bond distances are within 3.5 Å.

**Figure 8.**
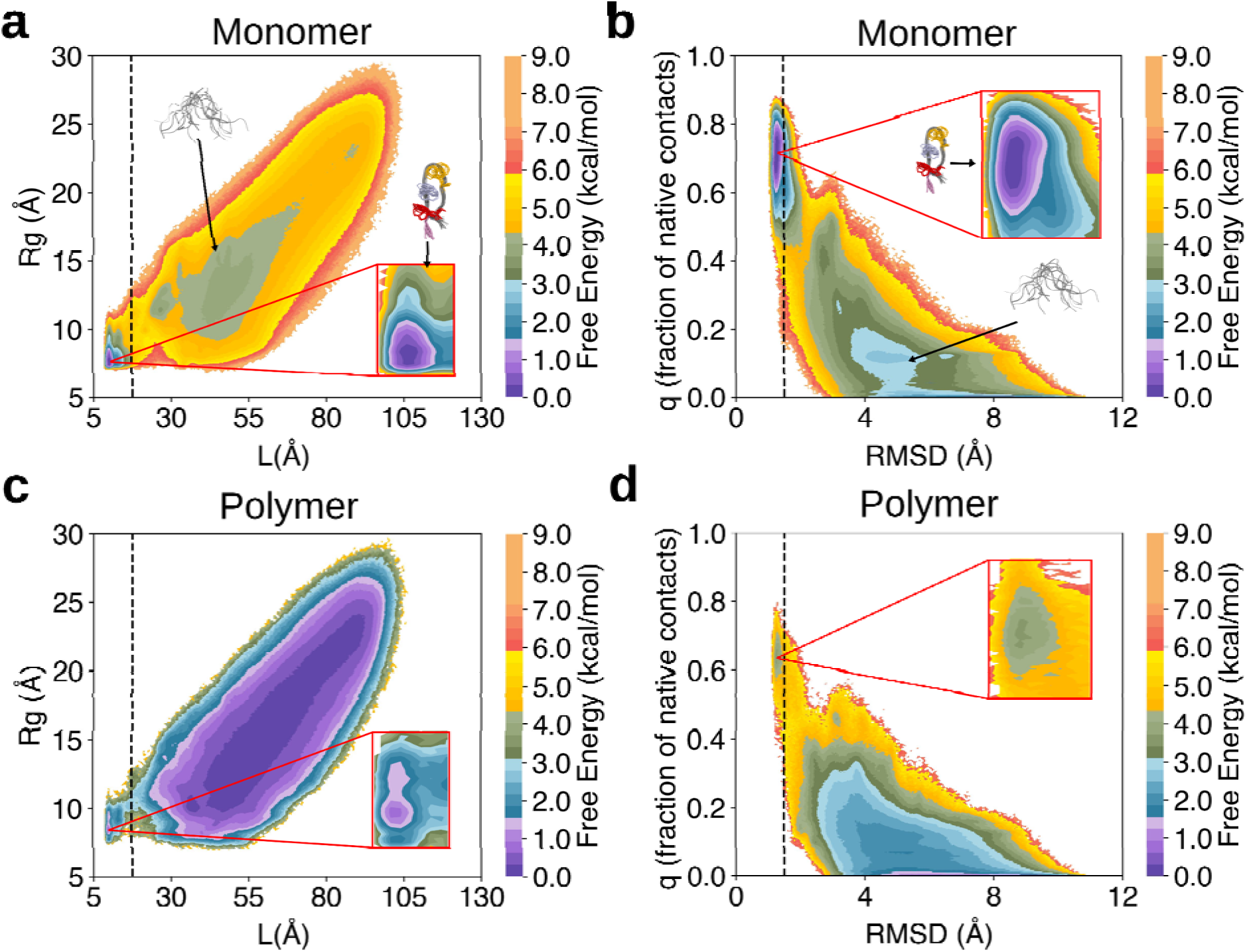
Structural comparison of simulation data for monomer and Ser-linked β-hairpin polymer systems. **(a)** Free energy landscape of the β-monomer projected onto the radius of gyration (sum of radii of the two tryptophan pairs) (*y*-axis) and sum of native hydrogen bonds (*x*-axis). The vertical dashed line indicates the folding threshold at 17.5 Å. **(b)** Free energy landscape of the β-monomer projected onto root mean square deviation (RMSD) from the reference NMR structure (*x*-axis) and fraction of native contacts (*y*-axis). The vertical dashed line marks the RMSD threshold at 1.5 Å. **(c)** Free energy landscape of the 15-mer polymer system projected onto the radius of gyration (*y*-axis) and sum of native hydrogen bonds (*x*-axis). **(d)** Free energy landscape of the 15-mer polymer system projected onto RMSD from the reference NMR structure (*x*-axis) and fraction of native contacts (*y*-axis).

This structural divergence from the native fold is further highlighted when examining similarity metrics relative to the reference NMR structure. The monomer exhibits a well-populated, stable minimum centered near a high native contact fraction (*q* ∼ 0.7) and within a native RMSD threshold of 1.5 Å (**Figure 8b**). In contrast, the corresponding region on the polymer landscape exhibits a high free energy penalty, with the populated ensemble shifting entirely to disordered conformations characterized by low native contacts (*q* < 0.2) and expanded RMSD values (**Figure 8d**). The suppression of this minimum demonstrates that polymer-mounted peptides face severe energetic barriers when attempting to approach the native reference conformation.

Collectively, these thermodynamic and structural analyses demonstrate that the stability of the folded β-hairpin conformation is drastically altered on the polymer scaffold, providing a molecular rationale for the significant fold perturbations observed via circular dichroism and infrared spectroscopy.

## Conclusions

Herein, we aimed to understand how polymerization of the trpzip2 motif into a brush polymer configuration would influence the folding of the individual trpzip2-based brushes. Generally, we observed a significant loss of CD intensity upon polymerization. As the intensity of the trpzip2 CD spectra are attributed to the strong excitonic interaction between the two pairs of tryptophan edge-to-face interactions within the structure, the significant loss in intensity suggests that polymerization disrupts these interactions. Additionally, there are shifts in CD peak position that are suggestive of weakened tryptophan pair interactions. These results were corroborated by transmission FT-IR spectroscopy studies that are consistent with loss of β-sheet hydrogen bonding content, and concurrent increases in random coil and β-turn content. The disruption of these tryptophan pair interactions and spectroscopic data suggest that the β-hairpin motif is significantly destabilized upon polymerization, and the degree of polymerization-induced unfolding is consistent across all samples and polymer lengths tested.

Simulation data provided mechanistic insight into the destabilization of the β-hairpin motif upon incorporation of trpzip2 into a polymeric scaffold. In contrast to the monomeric system, which exhibits a stable and well-populated native state with favorable thermodynamic and structural properties, the polymer-linked peptides display a marked shift in conformational preference toward unfolded or partially folded states. This shift is accompanied by an increase in root-mean-square deviation from the reference structure, a reduction in native contact formation, and a flattening of the free energy barriers separating metastable states. Furthermore, the emergence of a compact, globular conformation of the polymer backbone imposes additional steric constraints that limit the structural accessibility of the native fold. Collectively, these findings demonstrate that polymerization significantly reshapes the conformational free energy landscape of trpzip2, thereby destabilizing its native secondary structure and rationalizing the experimental findings. These results can be compared to those found in similar peptide brush polymers of α-helical monomers, which yielded polymers of significantly enhanced helicity upon polymerization.^12^ In the case of helicity, it is hypothesized that the locally hydrophobic environment of the polymer promotes folding, and the sterically crowded nature of the polymer helps to stabilize the α-helical fold by shielding water from the interior, in contrast to the destabilization observed in the case of trpzip2. These two systems illustrate that the folding of peptides appended as dense brushes on these polymers can be influenced differently by the scaffold they are conjugated to, providing guidelines for designing functional brush polymer architectures where sidechain peptide conformation is desirable, or not, for the function of the material. Finally, the inhibition of β-sheet character observed in peptide brush polymers in this report may thus be useful to impede the progression of harmful protein folding, for example in tau-derived fibrillization processes relevant in therapeutics for neurodegenerative disease.^54^

## ASSOCIATED CONTENT

### Supporting Information

The following files are available free of charge.

Supporting information containing additional detailed methods, synthetic procedures, and characterization of monomers and polymers. (PDF)

## Author Contributions

The manuscript was written through contributions of all authors. All authors have given approval to the final version of the manuscript.

## Notes

The views and conclusions contained in this document are those of the authors and should not be interpreted as representing the official policies, either expressed or implied, of the Army Research Office or the U.S. Government. The U.S. Government is authorized to reproduce and distribute reprints for Government purposes notwithstanding any copyright notation herein. The authors declare no competing financial interest.

## ACKNOWLEDGMENTS

J.O. acknowledges support through an NIH NRSA F32 fellowship from the National Institute of General Medical Sciences (NIGMS; F32GM143925). O.M.E. acknowledges support from the NSF Graduate Research Fellowship (Grant No. 2234667). We thank Dr. Nathan Rosenmann, Dr. Shirish Chodankar, Dr. Xinqi Chen, Hannah Palomino, and Jozsef Toth for helpful discussions and experimental assistance in the course of this study. This work made use of the IMSERC MS facility at Northwestern University, which has received support from the Soft and Hybrid Nanotechnology Experimental (SHyNE) Resource (NSF ECCS-2025633), and Northwestern University. This work also made use of the IMSERC NMR facility at Northwestern University, which has received support from the Soft and Hybrid Nanotechnology Experimental (SHyNE) Resource (NSF ECCS-2025633), NIH 1S10OD012016-01 / 1S10RR019071-01A1, and Northwestern University. This work made use of the Keck-II facility of Northwestern University’s NUANCE Center, which has received support from the SHyNE Resource (NSF ECCS-2025633), the IIN, and Northwestern’s MRSEC program (NSF DMR-2308691). Additionally, this work used instruments in the Keck Biophysics Facility, a shared resource of the Robert H. Lurie Comprehensive Cancer Center of Northwestern University supported in part by the NCI Cancer Center Support Grant P30 CA060553.

SAXS data were measured at Beamline 16-ID of the National Synchrotron Light Source II, part of the Center for BioMolecular Structure (CBMS), which is primarily supported by the National Institutes of Health, National Institute of General Medical Sciences (NIGMS) through a P30 Grant (P30GM133893), and by the DOE Office of Biological and Environmental Research (KP1605010). LiX also received additional support from NIH Grant S10 OD012331. As part of NSLS-II, a national user facility at Brookhaven National Laboratory, work performed at the CBMS is supported in part by the U.S. Department of Energy, Office of Science, Office of Basic Energy Sciences Program under contract number DE-SC0012704. We thank the citizen-scientists of Folding@home for donating their computing resources that enabled the distributed simulations (projects 12113-12118). This work benefited from the use of the SasView application, originally developed under NSF award DMR-0520547. SasView contains code developed with funding from the European Union’s Horizon 2020 research and innovation program under the SINE2020 project, grant agreement No 654000.

## ABBREVIATIONS

CD, circular dichroism; NMR, nuclear magnetic resonance; RMSD, root-mean-square deviation; SAXS, small-angle X-ray scattering; HPLC, high-performance liquid chromatography; ESI-MS, electrospray ionization-mass spectrometry; ROMP, ring-opening metathesis polymerization; FT-IR, Fourier transform infrared spectroscopy; IR, infrared; MD, molecular dynamics; MSM, Markov state model; OEG, oligo(ethylene glycol); SDS-PAGE, sodium dodecyl sulfate– polyacrylamide gel electrophoresis; SEC-MALS, size-exclusion chromatography coupled with multi-angle light scattering; SPPS, solid-phase peptide synthesis; tICA, time-lagged independent component analysis.

