## Supporting Information for "β-Hairpin Fold Peptide Brush Polymers"

1. Materials

All amino acids used to prepare peptides by solid-phase peptide synthesis (SPPS) were obtained from commercial sources. Lithium chloride (99%), Fmoc-Asp-O^t^Bu, 6-(Fmoc-amino)hexanoic acid (97%) were purchased from commercial sources and used without purification. N-(hexanoic acid)-cis-5-norbornene-exo-dicarboximide and the olefin metathesis initiator (IMesH_2_)(C_5_H_5_N)_2_(Cl)_2_Ru=CHPh were synthesized according to published procedures.^1–3^

2. Synthesis of Fmoc-Asn(NorIm)-OH

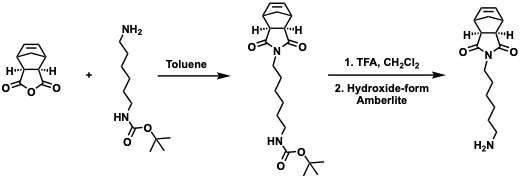

**Boc-Protected Norbornene-C6 Amine**

A procedure similar to one previously published^4^ was pursued. Specifically, to a stirred solution of *cis*-5-norbornene-*exo*-2,3-dicarboxylic anhydride (1.00 g, 6.09 mmol) in toluene (40 mL) was added N-Boc-1,6-hexanediamine (1.45 g, 6.70 mmol, 1.1 equiv). The reaction was heated to reflux with a Dean Stark trap in place for 48 h. Upon cooling to room temperature, the reaction mixture was washed with 1 M HCl (×3), the organic layer was dried over Na_2_SO_4_, filtered and concentrated to dryness to give a pale yellow/white solid, 2.0 g (91%). ^1^H NMR (CDCl_3_): δ (ppm) 1.17 (d, 1H, 1 × CH2, J = 9.9 Hz), 1.27 (m, 4H, 2 × CH2), 1.38–1.60 (m, 14H, 3 × CH3, 2 × CH2, CH), 2.62 (d, 2H, CH2), 3.04 (m, 2H, CH2), 3.22 (m, 2H, CH2), 3.40 (dd, 2H, CH2, J = 8.3, 6.6 Hz), 4.57 (bs, 1H NH), 6.24, (t, 2H, HC=CH, J = 1.9 Hz). ^13^C{1H} NMR (CDCl_3_): δ (ppm) 26.27, 26.59, 27.68, 28.46, 29.89, 38.55, 40.44, 42.75, 42.86, 45.05, 45.18, 47.62, 47.82, 78.98, 137.85, 156.00, 178.09. ESI-MS: 385.31 [M+Na]^+^

**Norbornene-C6 Amine**

The protected amine (0.60 g, 1.98 mmol) was dissolved in a 1:1 TFA : CH_2_Cl_2_ solution (20 mL) and stirred at room temperature for 6 h. The reaction was concentrated to dryness to give a brown residue that was precipitated by the addition of ether. Removal of the solvent gave the free amine as a yellow oil.

After synthesis, NorIm (4 g) was dissolved in water (ca. 10 mL) and then treated with hydroxide-form Amberlite™ resin (IRN78, ca. 10 mL) until the pH of the solution above the resin was basic. The resin was filtered off and washed with water three times, and the combined filtrate was collected and lyophilized to yield a yellow oil. ^1^H NMR (CDCl_3_) δ 1.19 (2, 1H, CH2), δ 1.29 (m, 2H, CH2), δ 1.38 (m, 2H, CH2), δ 1.52 (m, 3H, CH2), δ 1.63 (m, 2H, CH2), δ 2.66 (d, 2H, CH2, J = 1.33 Hz), δ 2.87 (t, 2H, CH2, J = 7.53 Hz), δ 3.25 (t, 2H, CH2, J = 1.7 Hz), δ 3.44 (q, 2H, CH2, J = 6.58 Hz), δ 4.90 (s, 5H, NH2), δ 6.27 (t, 2H, HC=CH, J = 1.86 Hz). ^13^C{1H} NMR (CDCl_3_): δ (ppm) 25.90, 26.33, 27.48, 28.91, 38.43, 40.32, 42.77, 45.20, 47.87, 113.15, 115.48, 117.81, 120.14, 137.88, 161.73, 162.01, 162.29, 162.57, 178.31.

**Fmoc-Asn(NorIm)-O^t^Bu**

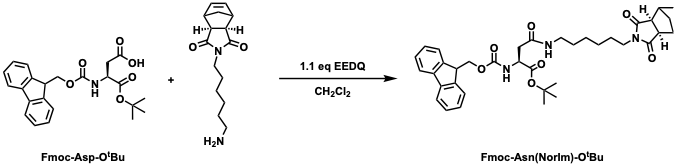

A solution of Fmoc-Asp-O^t^Bu (1.5 g, 3.6 mmol, 1 equiv) was dissolved in dichloromethane (25 mL) in a round-bottom flask, and EEDQ (0.99 g, 4 mmol, 1.1 equiv) was added while stirring. The solution was left to stir at room temperature for 20 min before adding NorIm (0.96 g, 3.6 mmol, 1 equiv) dissolved in dichloromethane (5 mL) dropwise. The yellow solution was left to stir overnight at room temperature, and then was dried using a rotary evaporator before being purified using column chromatography (Sfär Silica D Duo 60 μm 25 g, 20% EtOAc in hexanes to 100% EtOAc over 20 column volumes). The fractions containing the majority product were dried using a rotary evaporator and then repurified using column chromatography (Sfär Silica D Duo 60 μm 25 g, 20% EtOAc in hexanes to 100% EtOAc over 20 column volumes) to remove remaining trace Fmoc-Asp-O^t^Bu, to yield a yellow oil (1.2 g, 51% yield).

**Fmoc-Asn(NorIm)-OH**

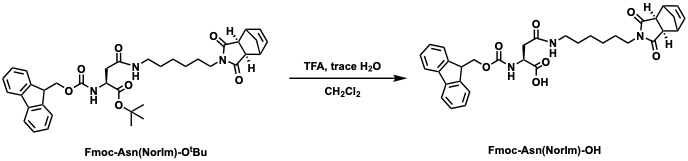

The compound Fmoc-Asn(NorIm)-O^t^Bu (1.2 g, 1.8 mmol, 1 equiv) was dissolved in dichloromethane (30 mL) and trifluoroacetic acid (18 mL) as well as trace water (60 μL) were added to a round-bottom flask. The solution was stirred for four hours before being concentrated using a rotary evaporator. The reaction mixture was then resuspended in water and concentrated again, three times, to remove trace protecting group side products, and then redissolved in methanol and dried three times to remove trace trifluoroacetic acid and isolate the final product, an off-white powder (0.735 g, 1.2 mmol, 67% yield).

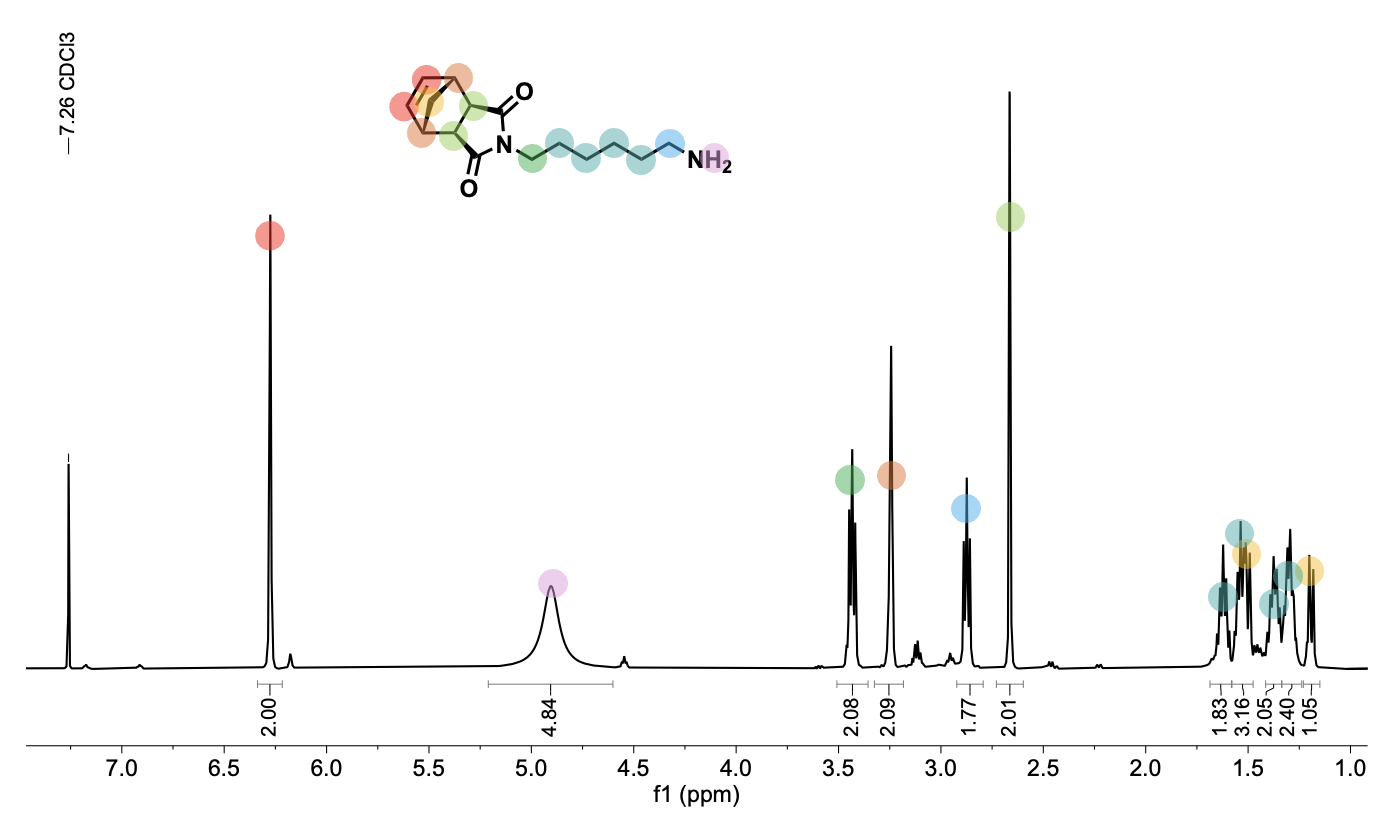

Figure S1. ^1^H NMR spectrum of NorIm-amine after being treated with Amberlite^™^ resin dissolved in CDCl_3_, obtained on a 500 MHz spectrometer at 25 °C.

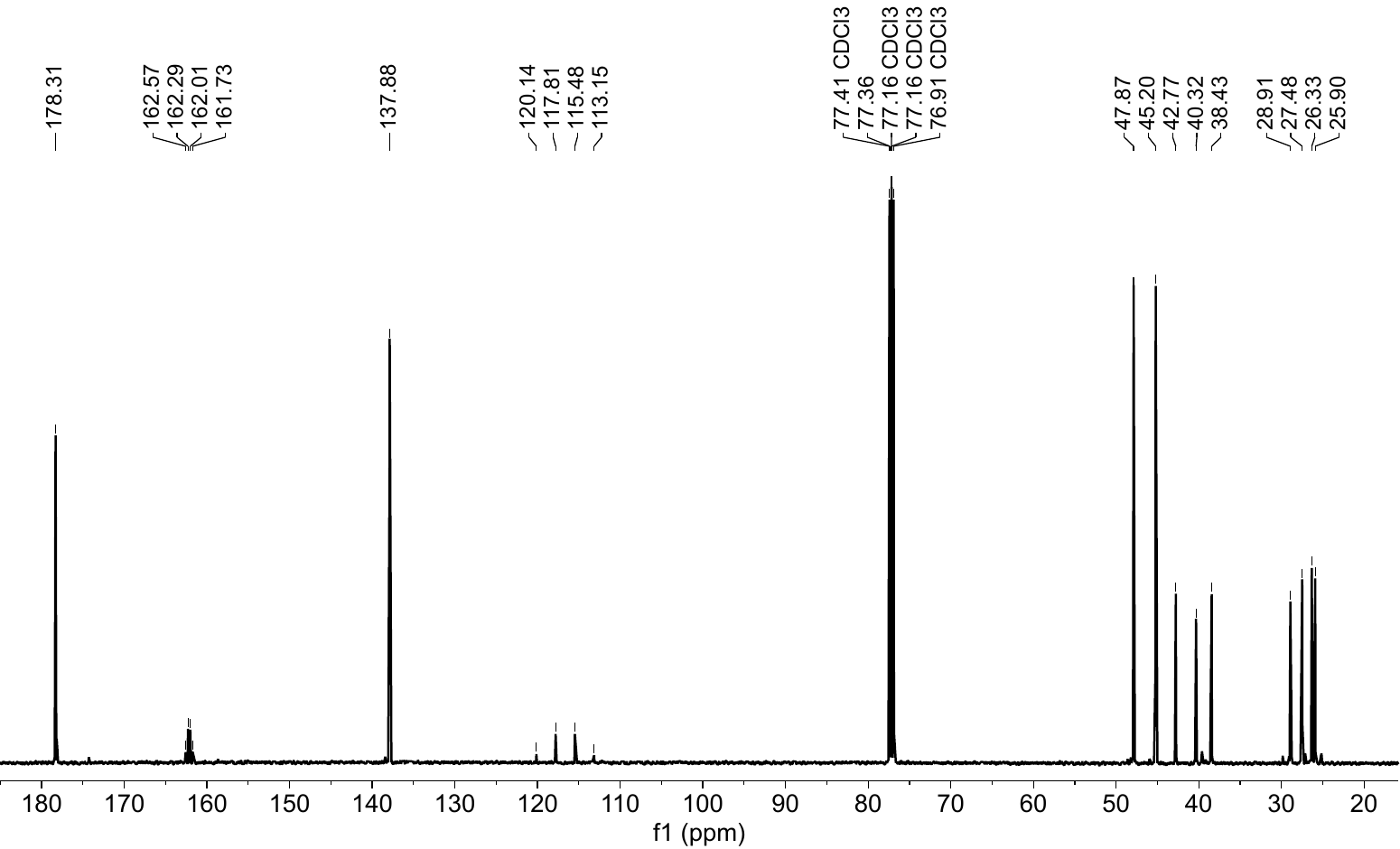

Figure S2. ^13^C{H} NMR spectrum of NorIm-amine after being treated with Amberlite^™^ resin dissolved in CDCl_3_, obtained on a 500 MHz spectrometer at 25 °C.

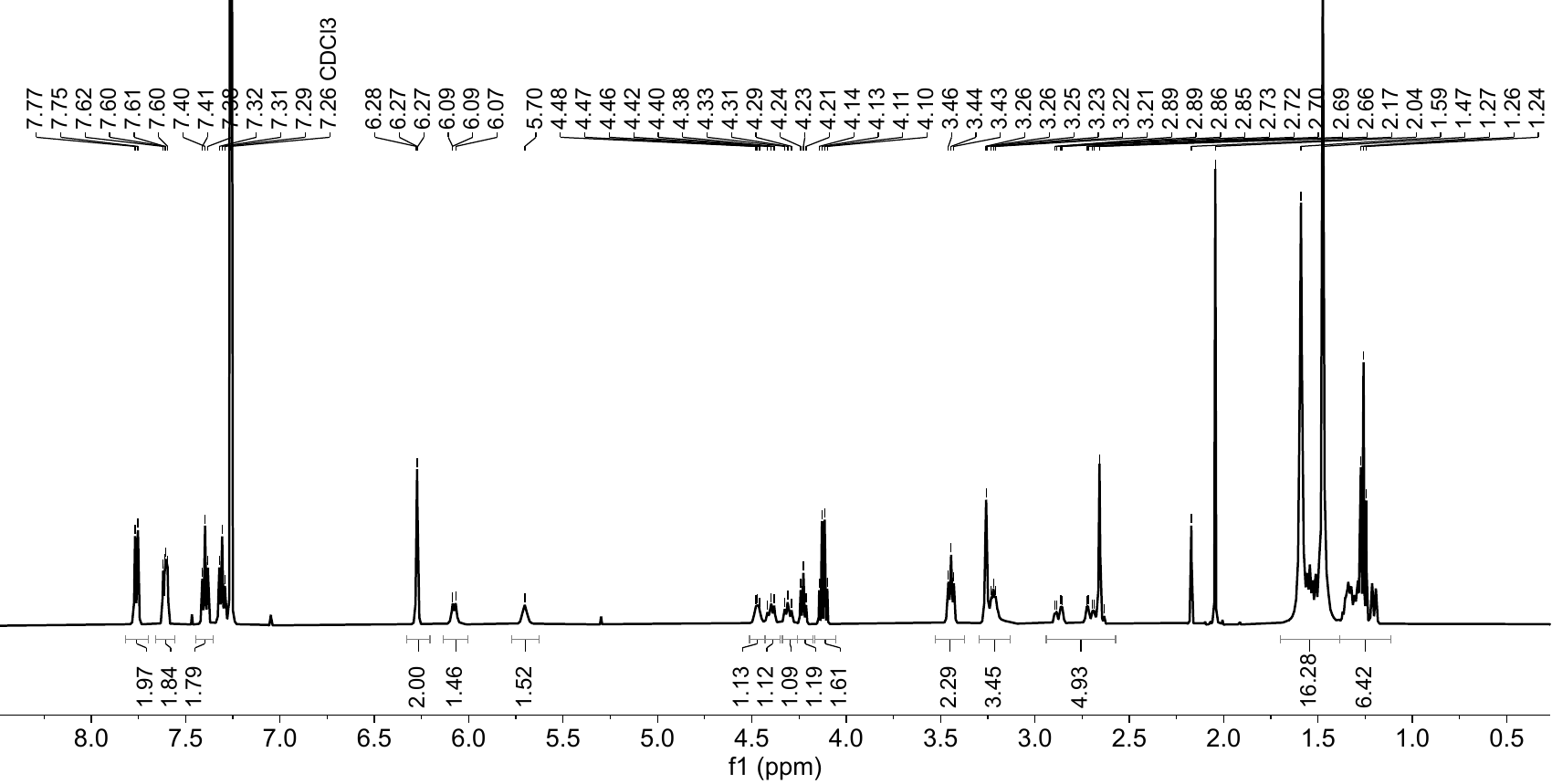

Figure S3. ^1^H NMR spectrum of Fmoc-Asn(NorIm)-OtBu dissolved in CDCl_3_, obtained on a 500 MHz spectrometer at 25 °C.

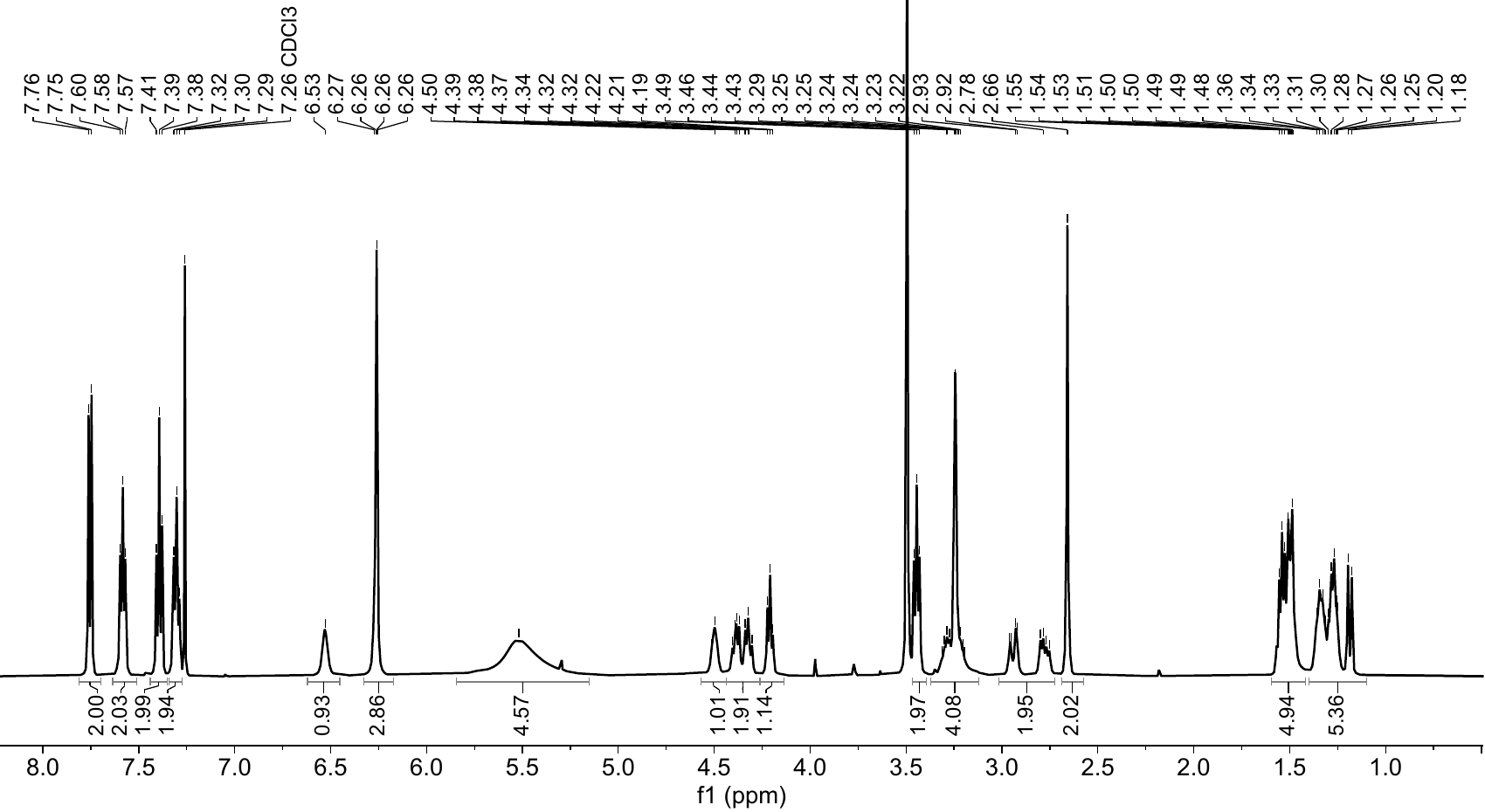

Figure S4. ^1^H NMR spectrum of Fmoc-Asn(NorIm)-OH dissolved in CDCl_3_, obtained on a 500 MHz spectrometer at 25 °C.

3. Synthesis and Purification of Peptide Monomers

Peptide Synthesis: Peptides were synthesized using a Liberty Blue (CEM) automated microwave peptide synthesizer, utilizing Fmoc-protected amino acids and standard solid-phase peptide synthesis (SPPS) protocols. All peptides were generated on Rink amide MBHA resin to give C-terminal amidated peptides. Peptide monomers, unless otherwise indicated, were capped with N-(hexanoic acid)-*cis*-5-norbornene-*exo*-2,3-dicarboximide (“NorAha”) using SPPS. The random coil monomer, “ran,” the sequence of which is NorImAha-GSGSGRGSGSGE-NH_2_, was made as previously described.^5^

Cleavage from the resin was performed in 95 : 2.5 : 2.5 (% v/v) trifluoroacetic acid (TFA), water, and triisopropylsilane, respectively, for three and a half hours. After filtration, the resin was washed three times with fresh trifluoroacetic acid and dichloromethane, before all fractions were concentrated using a rotary evaporator and precipitated in cold diethyl ether. The precipitated product was dried under vacuum to yield solid crude peptide that was purified as detailed below. Crude samples that retained protecting groups, commonly trace Boc on the tryptophan residues and could be detected as shoulders on HPLC traces, were treated with 5% TFA water/acetonitrile solutions for an additional two hours at room temperature before being lyophilized and purified as detailed.

Analytical High-Performance Liquid Chromatography (HPLC): Analytical HPLC analysis of peptides was performed on a Jupiter 4μ Proteo 90Å Phenomenex column (150 × 4.60 mm) using a Hitachi-Elite LaChrom L-2130 pump equipped with UV-Vis detector (Hitachi-Elite LaChrom L2420). The solvent system consists of (A) 0.1% v/v TFA in water and (B) 0.1% v/v TFA in acetonitrile. After one column volume was monitored (3 min), the solvent gradient was allowed to run (30 min) before a wash step (not shown).

Preparative Flash Chromatography: All peptides were purified using a Biotage Selekt automated flash chromatography system. The solvent system consisted of (A) 0.1% TFA in water and (B) 0.1% TFA in acetonitrile, and Biotage Sfär C18 D Duo 100 Å, 30 μm columns were used for purification. Samples were dissolved in minimal solvent (~10 mg/mL), filtered through a 0.22 μm PES filter, and run using solvent gradients optimized by examination of analytical HPLC data.

Preparative High-Performance Liquid Chromatography (HPLC): Additional purification was performed using either a Gilson PLC 2050 purification system or a BUCHI C-380 Prep purification system. The solvent system consisted of (A) 0.1% TFA in water and (B) 0.1% TFA in acetonitrile. Samples were dissolved (<10 mg/mL), filtered through a 0.22 μm PES filter, and run using solvent gradients optimized by examination of analytical HPLC data. For the beta peptide monomer, the optimized gradient was 20–50% B.

Electrospray Ionization Mass Spectrometry (ESI-MS): ESI-MS spectra of peptides were collected using a Bruker amaZon SL mass spectrometer configured with an ESI source in both negative and positive ionization modes, using a mobile phase of 50% water and 50% acetonitrile.

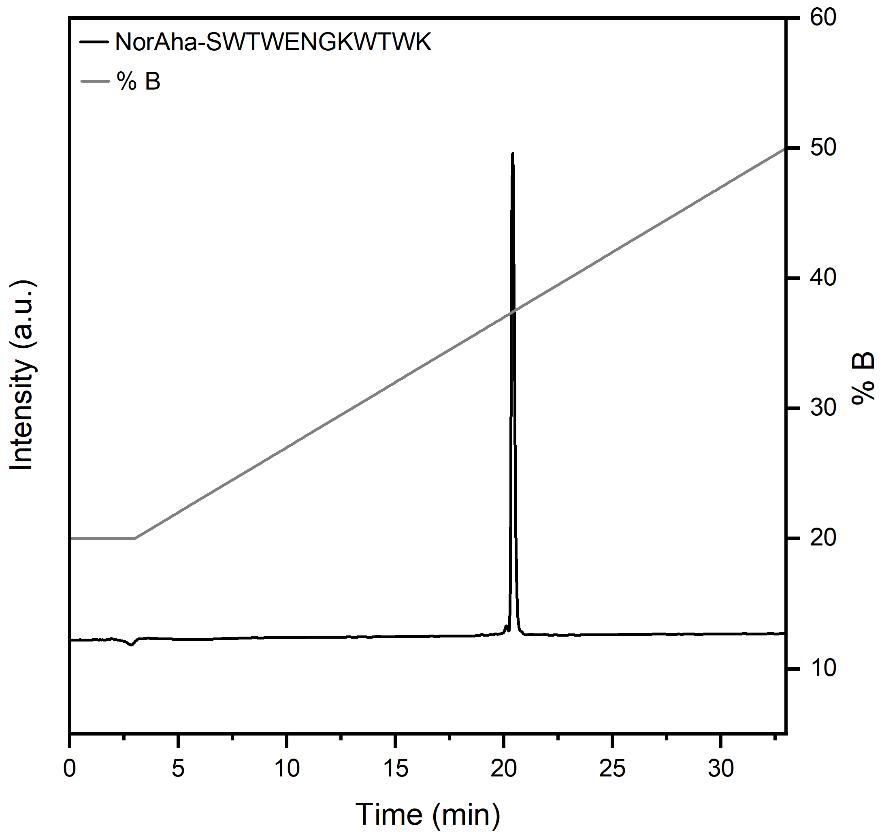

Figure S5. Analytical HPLC trace (obtained at 214 nm) of the trpzip2-based peptide monomer, NorAha-SWTWENGKWTWK-NH_2_, obtained after purification using preparative HPLC, using a solvent gradient of 20 to 50% B.

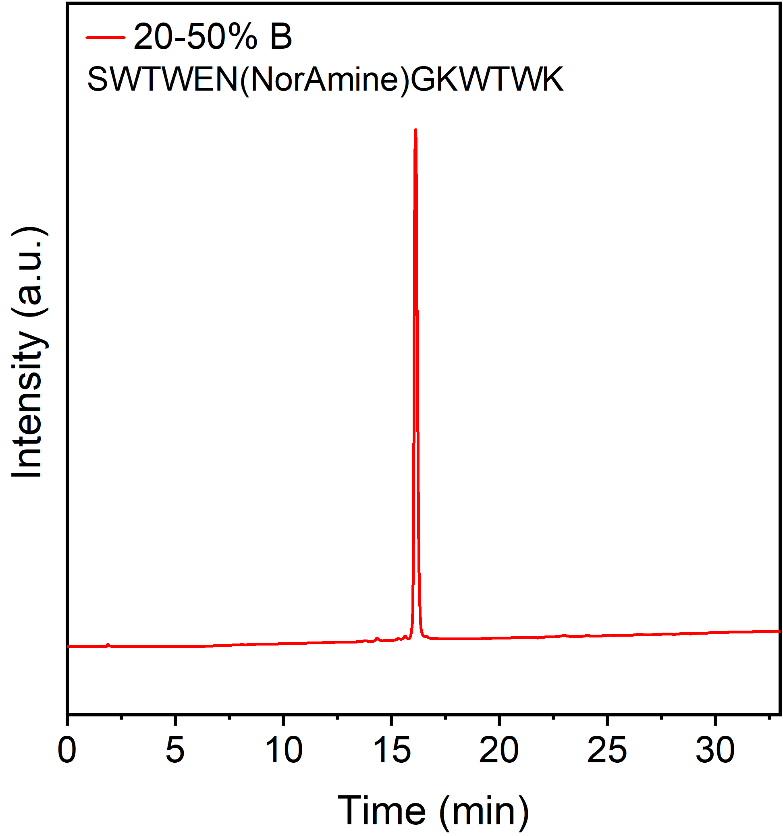

Figure S6. Analytical HPLC trace (obtained at 214 nm) of the modified trpzip2 monomer, N-beta or SWTWEN(NorAmine)GKWTWK-NH_2_, obtained after purification using preparative HPLC, using a solvent gradient of 20 to 50% B.

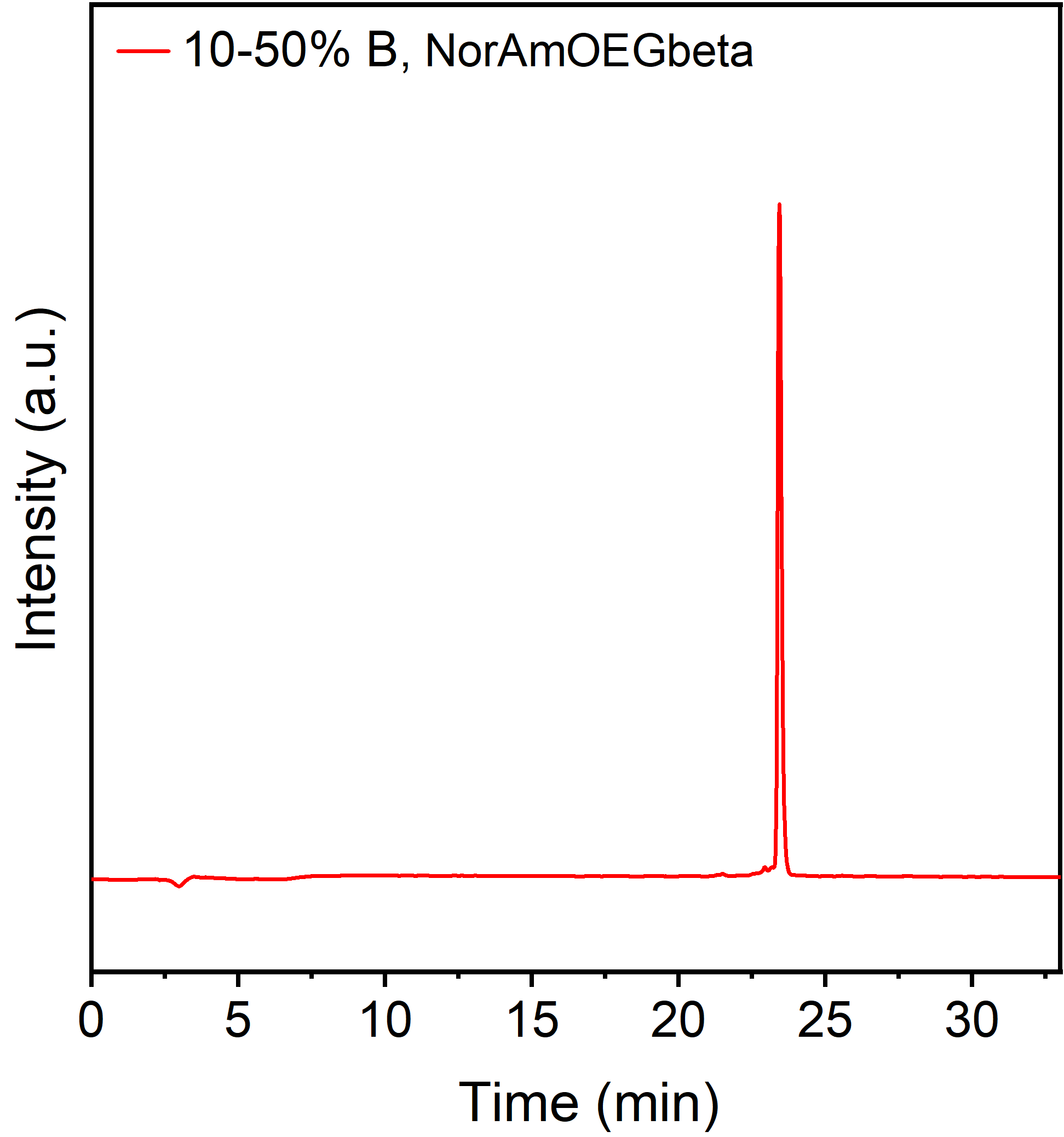

Figure S7. Analytical HPLC trace (obtained at 214 nm) of the modified trpzip2 monomer, NorAmOEG-beta or NorAmOEG-SWTWENGKWTWK-NH_2_, obtained after purification using preparative HPLC, using a solvent gradient of 10 to 50% B.

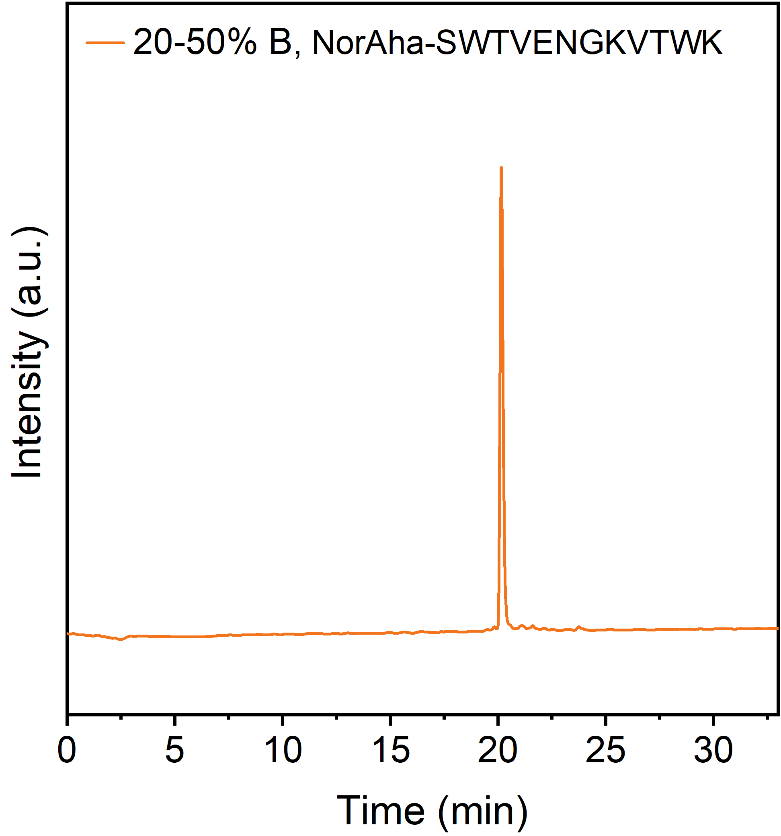

Figure S8. Analytical HPLC trace (obtained at 214 nm) of the modified trpzip2 monomer, WVVW-beta or NorAha-SWTVENGKVTWK-NH_2_, obtained after purification using preparative HPLC, using a solvent gradient of 20 to 50% B.

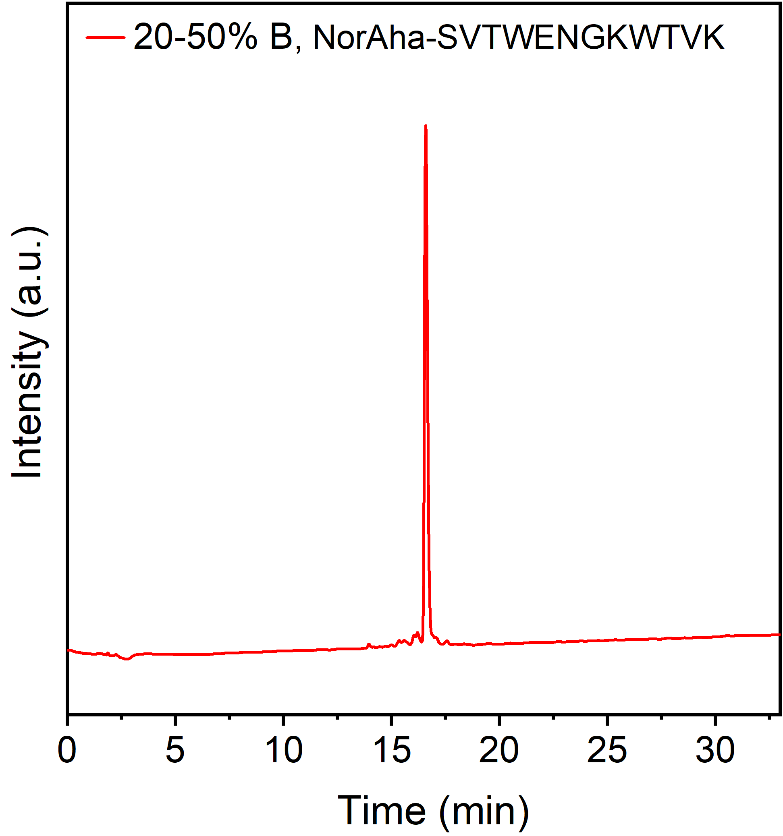

Figure S9. Analytical HPLC trace (obtained at 214 nm) of the modified trpzip2 monomer, VWWV-beta or NorAha-SVTWENGKWTVK-NH_2_, obtained after purification using preparative HPLC, using a solvent gradient of 20 to 50% B.

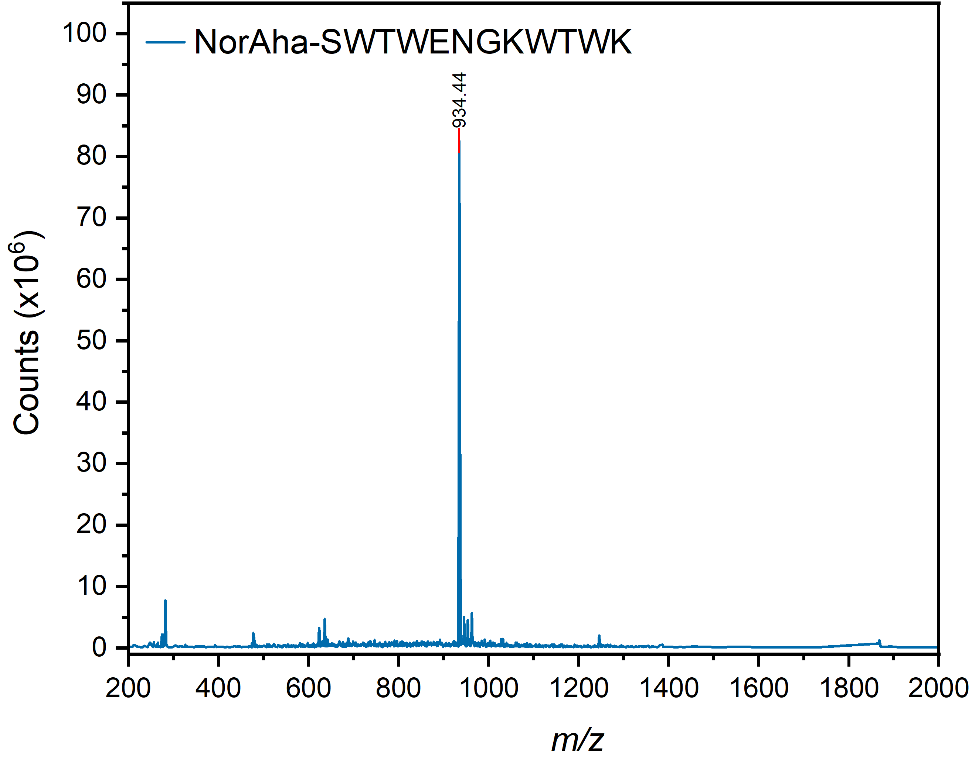

Figure S10. Plot of ESI-MS results of beta, NorAha-SWTWENGKWTWK-NH_2_, after purification, confirming the successful synthesis and purification of the expected peptide. Expected *m*/*z* of [M+H] = 1867.90, [M+2H] = 934.45.

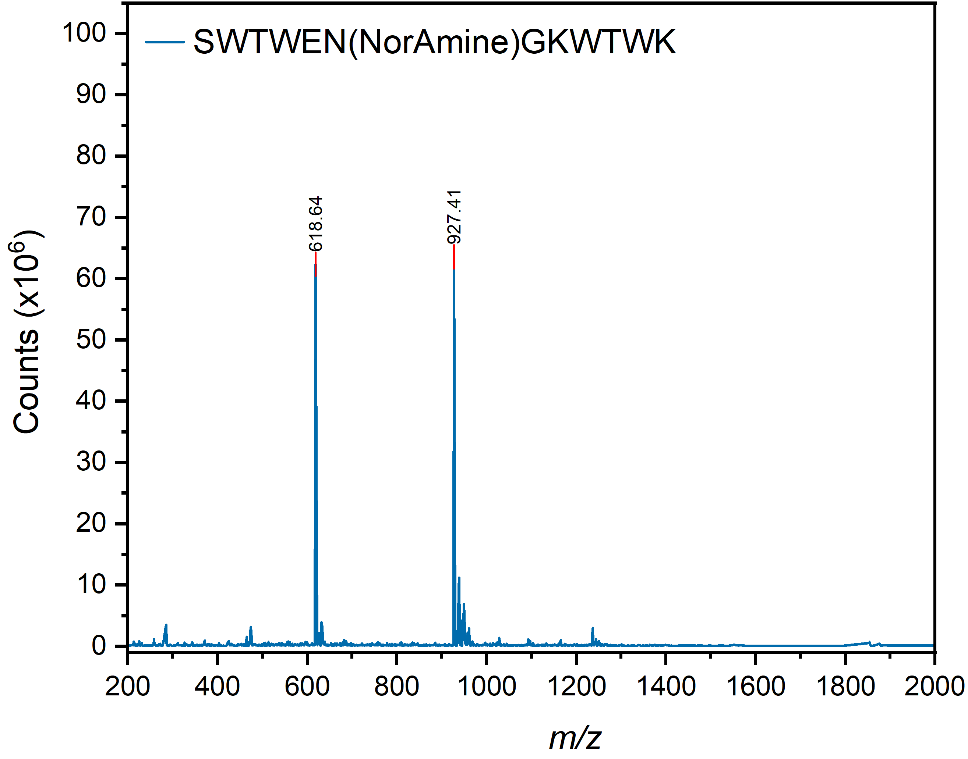

Figure S11. Plot of ESI-MS results of N-beta, SWTWEN(NorAmine)GKWTWK-NH_2_, after purification, confirming the successful synthesis and purification of the expected peptide. Expected *m*/*z* of [M+H] = 1853.92, [M+2H] = 927.46, [M+3H] = 618.64.

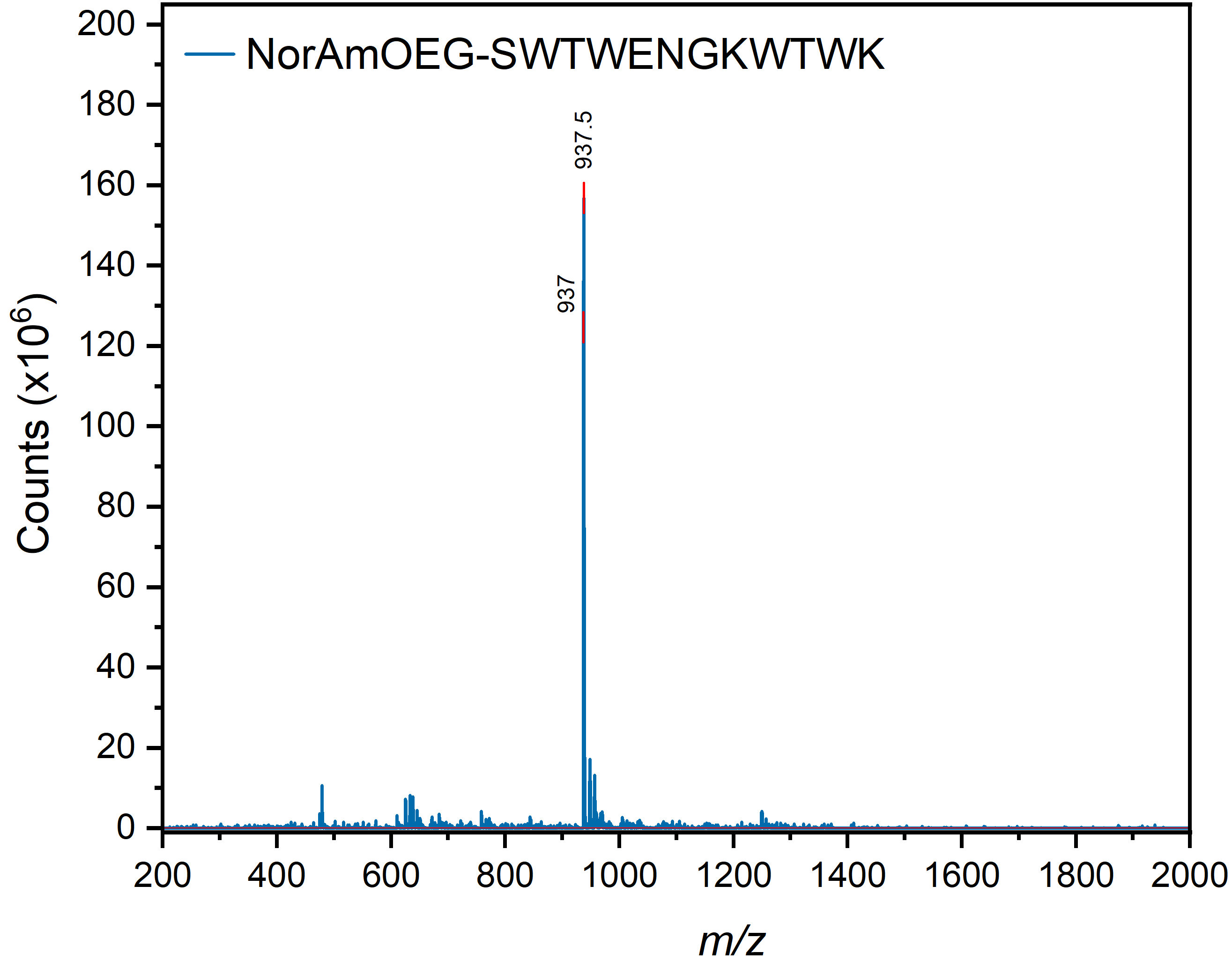

Figure S12. Plot of ESI-MS results of NorAmOEG-beta, NorAmOEG-SWTWENGKWTWK-NH_2_, after purification, confirming the successful synthesis and purification of the expected peptide. Expected *m*/*z* of [M+H] = 1873.91, 1872.91, [M+2H] = 937.46, 936.96.

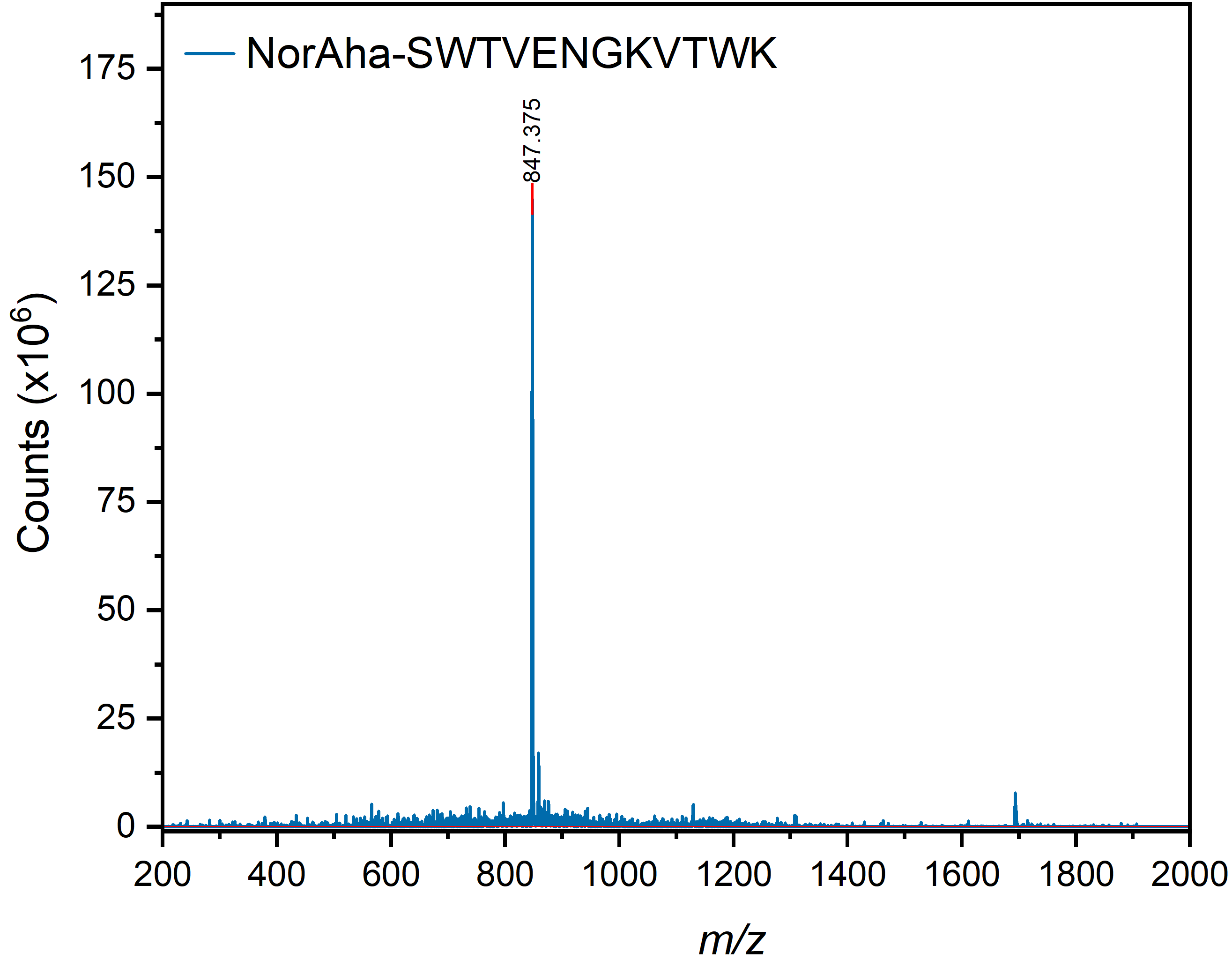

Figure S13. Plot of ESI-MS results of VWWV-beta, NorAha-SVTWENGKWTVK-NH_2_, after purification, confirming the successful synthesis and purification of the expected peptide. Expected *m*/*z* of [M+H] = 1692.88, 1693.88; [M+2H] = 846.94, 847.44.

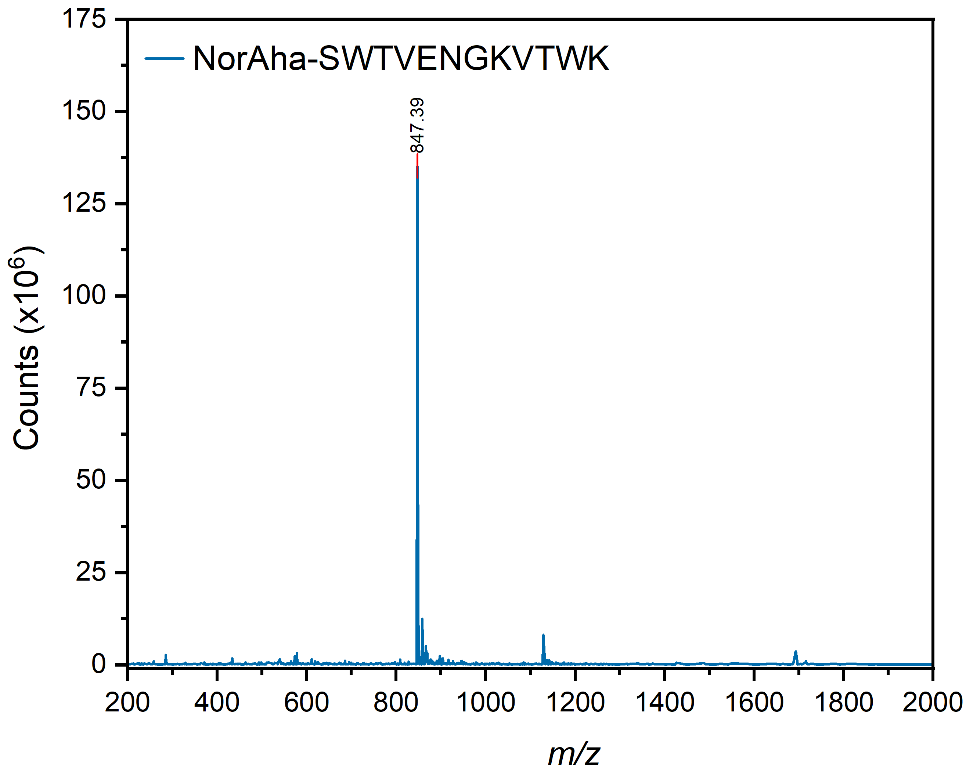

Figure S14. Plot of ESI-MS results of WVVW-beta, NorAha-SWTVENGKVTWK-NH_2_, after purification, confirming the successful synthesis and purification of the expected peptide. Expected *m*/*z* of [M+H] = 1692.88, 1693.88; [M+2H] = 846.94, 847.44.

4. Polymerization and Characterization of Polymers

Polymerization reactions were carried out in a purge box under a nitrogen atmosphere, similar to procedures previously described.^6–9^ Solvent (*N,N*-dimethylformamide) was sparged with nitrogen gas immediately before use. Small-scale reactions were performed and monitored using ^1^H NMR spectroscopy to assess the polymerization of the monomer before polymerizations with larger batches (>0.05 g) of peptide monomer were attempted. ^1^H NMR spectra were measured on a Bruker Avance III spectrometer (400 MHz).

In a typical NMR-scale experiment, peptide monomer (15 to 45 mg) was dissolved in dry *N,N*-dimethylformamide (DMF, 0.5 mL) with or without lithium chloride (1 M; depending on solubility of polymer) and an internal standard (hexamethyldisilane, HMDS). The monomer solution was transferred to a J. Young NMR tube, sealed, and the NMR spectrum of the solution was measured. The integration of the norbornene alkene peak (6.3 ppm) was assessed relative to the integration of the hexamethyl peak (~0 ppm). Subsequently, the tube was brought into the purge box, a solution of Grubbs third-generation catalyst in DMF (1 equiv) was added, and the reaction was mixed. The NMR tube was sealed and then monitored regularly to observe the progress of the polymerization by observing the disappearance of the norbornene alkene peak. Typically, 60–150 min was required for complete conversion. The polymers were terminated with ethyl vinyl ether (>20 equiv) for 20 min at room temperature, precipitated, washed with cold diethyl ether 3 times, and collected by centrifugation.

The molecular weight and polydispersity of all polymers were determined by organic phase or aqueous phase gel permeation chromatography, or SDS-PAGE when polymers were found incompatible with gel permeation chromatography.

We note that the SDS-PAGE values consistently underestimate the approximate molecular weight of polymer samples, likely due to the fact the polymer topology might hinder the binding of sodium dodecyl sulfate in comparison to a protein. This has previously been observed in earlier work,^7^ where *M*_w_ values were obtained very close to the anticipated theoretical values via SEC-MALS, but MW values determined from SDS-PAGE significantly underestimated the weight of the polymer the longer the degree of polymerization. We also note the underestimation of dispersity values determined using SEC-MALS, which is a limitation of this technique.^10^

Organic Phase Gel Permeation Chromatography (Organic GPC): Organic phase GPC measurements were performed on a set of Phenomenex Phenogel 5μ, 1K-75K, 300 × 7.80 mm in series with a Phenomex Phenogel 5μ, 10K-1000K, 300 × 7.80 mm columns with HPLC grade solvents as eluents: dimethylformamide (DMF) with 0.05 M of LiBr at 60 °C. Detection consisted of a Hitachi UV-Vis Detector L-2420, a Wyatt Optilab T-rEX refractive index detector operating at 658 nm and a Wyatt DAWN® HELEOS® II light scattering detector operating at 659 nm.

Aqueous Phase Gel Permeation Chromatography (Aqueous GPC): Aqueous phase GPC measurements were performed with a Tosoh Bioscience TSK-GEL^®^PWxl-CP column 7.8 mm ID × 30 cm, 10 μm, using 0.05% NaN_3_ as the mobile phase. Detection consisted of a Wyatt Optilab refractive index detector operating at 658 nm and a Wyatt DAWN® light scattering detector operating at 659 nm.

Dynamic Light Scattering (DLS): DLS analyses were conducted at room temperature on a Zetasizer Nano-ZS (Malvern) operating with a 633 nm laser. Samples were prepared as 2.5 mg/mL solutions and measured at 25 °C with an equilibration time of 60 s prior to measurement.

SDS-PAGE: Samples were dissolved to a concentration of 1 mg/mL in aqueous buffer or water, before being combined with Laemmli sample buffer (30 μL sample, 20 μL buffer solution; Bio-Rad) and were heated at 95 °C for 10 min. The cooled samples were then loaded (20 μL) onto Mini-PROTEAN TGX Precast Protein Gel (Bio-Rad) with an acrylamide gel gradient of 4 to 15% and using a Tris/Gly/SDS running buffer (Bio-Rad). Molecular weight standards were utilized from Bio-Rad (Precision Plus Protein Dual Xtra). The gels were run at a voltage of 120 V for 1 h at room temperature, after which the gels were stained with Coomassie Brilliant Blue G dye solution (containing 40% v/v methanol and 10% v/v acetic acid) for 1 h. The gels were then destained (using solutions of 40% v/v methanol and 10% v/v acetic acid) until the gel background was clear while retaining stain in the sample and reference bands, then imaged using an Azure Biosystems Azure 600 Imaging System in the Keck Biophysics Facility at Northwestern University.

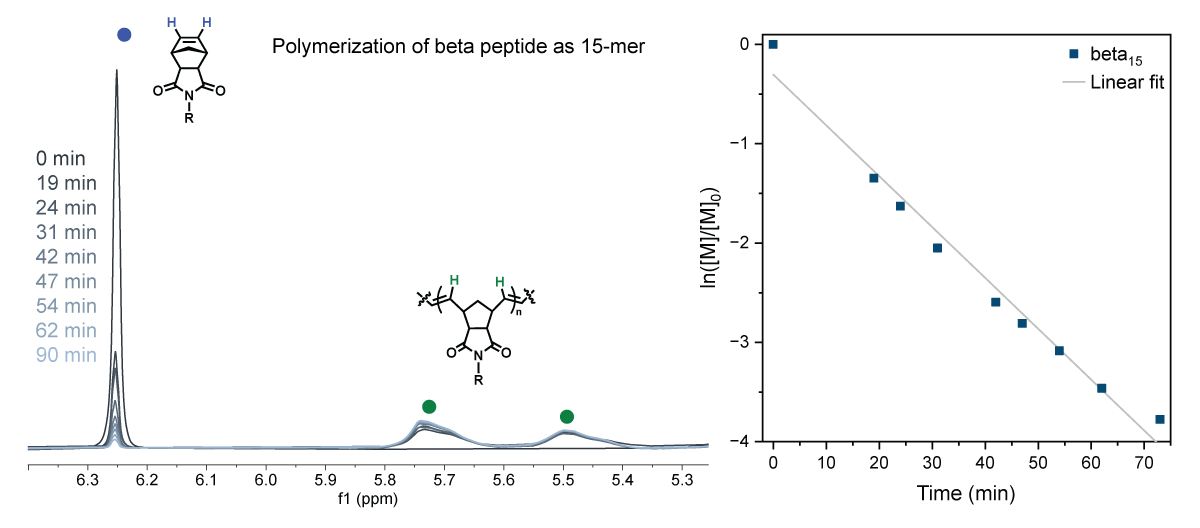

Figure S15. Representative polymerization of the trpzip2 peptide monomer, NorAha-SWTWENGKWTWK, using the Grubbs third-generation catalyst and targeting a 15-mer in length, monitored using ^1^H NMR spectroscopy. Left: a close-up of the alkene region of the monomer (peak indicated with a blue circle) and polymer (broad peaks indicated with green circles). Right: pseudo-first-order kinetics plot of the polymerization, generated by integrating the monomer alkene peak relative to the integration of an HMDS standard peak. The plots were fit to a linear line with a slope of −0.05118 and R^2^ = 97.6%.

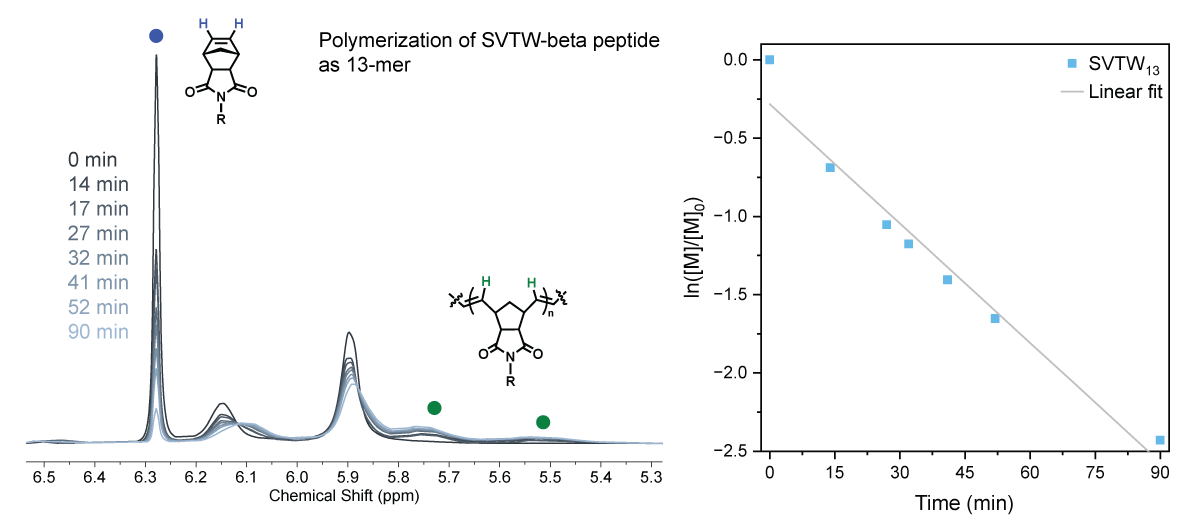

Figure S16. Representative polymerization of the SVTW-variant of trpzip2 peptide monomer, NorAha-SVTWENGKWTVK (“VWWV-beta”), using the Grubbs third-generation catalyst and targeting a 15-mer in length, monitored using ^1^H NMR spectroscopy. Left: a close-up of the alkene region of the monomer (peak indicated with a blue circle) and polymer (broad peaks indicated with green circles). Right: pseudo-first-order kinetics plot of the polymerization, generated by integrating the monomer alkene peak relative to the integration of an HMDS standard peak. The plots were fit to a linear line with a slope of −0.02542 and R^2^ = 95.8%.

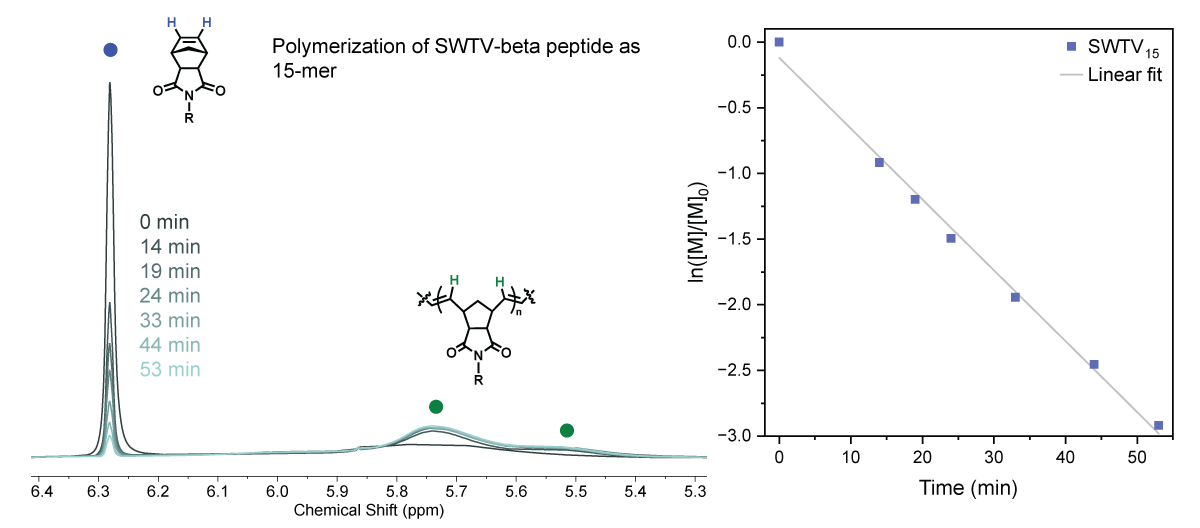

Figure S17. Representative polymerization of the SWTV-variant of trpzip2 peptide monomer, NorAha-SWTVENGKVTWK (“WVVW-beta”), using the Grubbs third-generation catalyst and targeting a 15-mer in length, monitored using ^1^H NMR spectroscopy. Left: a close-up of the alkene region of the monomer (peak indicated with a blue circle) and polymer (broad peaks indicated with green circles). Right: pseudo-first-order kinetics plot of the polymerization, generated by integrating the monomer alkene peak relative to the integration of an HMDS standard peak. The plots were fit to a linear line with a slope of −0.05398 and R^2^ = 99.3%.

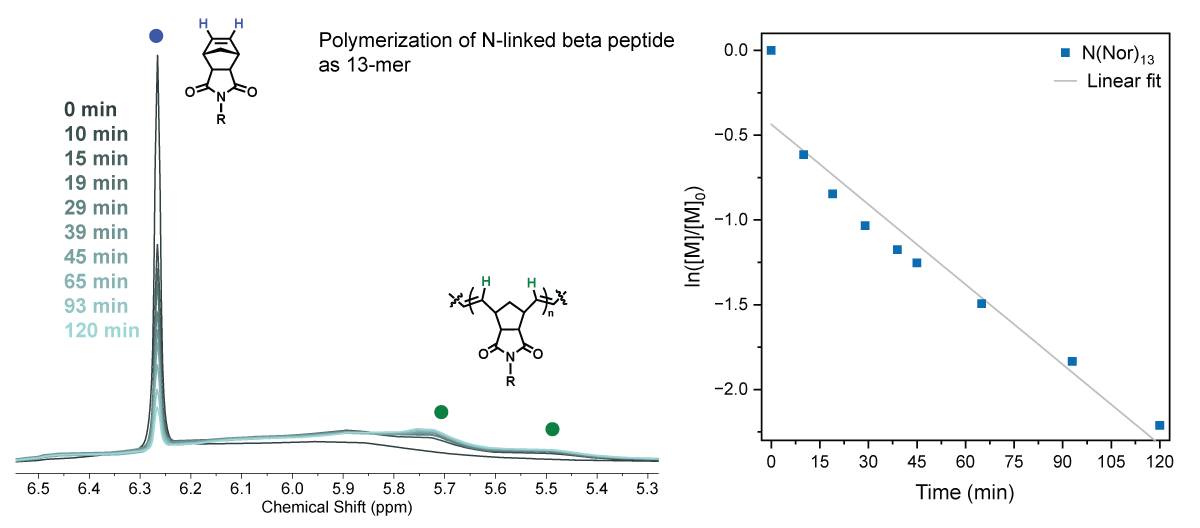

Figure S18. Representative polymerization of the N-linked variant of trpzip2 peptide monomer, SWTWEN(Nor)GKWTWK, using the Grubbs third-generation catalyst and targeting a 15-mer in length, monitored using ^1^H NMR spectroscopy. Left: a close-up of the alkene region of the monomer (peak indicated with a blue circle) and polymer (broad peaks indicated with green circles). Right: pseudo-first-order kinetics plot of the polymerization, generated by integrating the monomer alkene peak relative to the integration of an HMDS standard peak. The plots were fit to a linear line with a slope of −0.01573 and R^2^ = 91.4%.

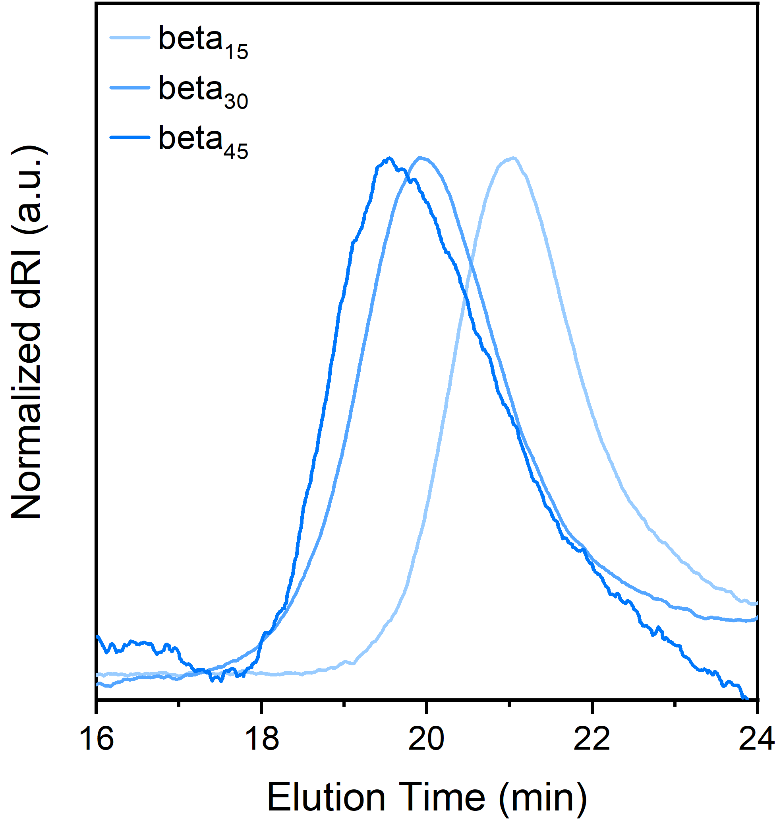

Figure S19. Organic SEC-MALS of polymers, beta_15_ (light blue trace), beta (blue trace), and beta_45_ (purple trace), run using 0.05 M LiBr in DMF. The differential refractive index was normalized to the highest and lowest values of the run’s data to allow for a more facile comparison.

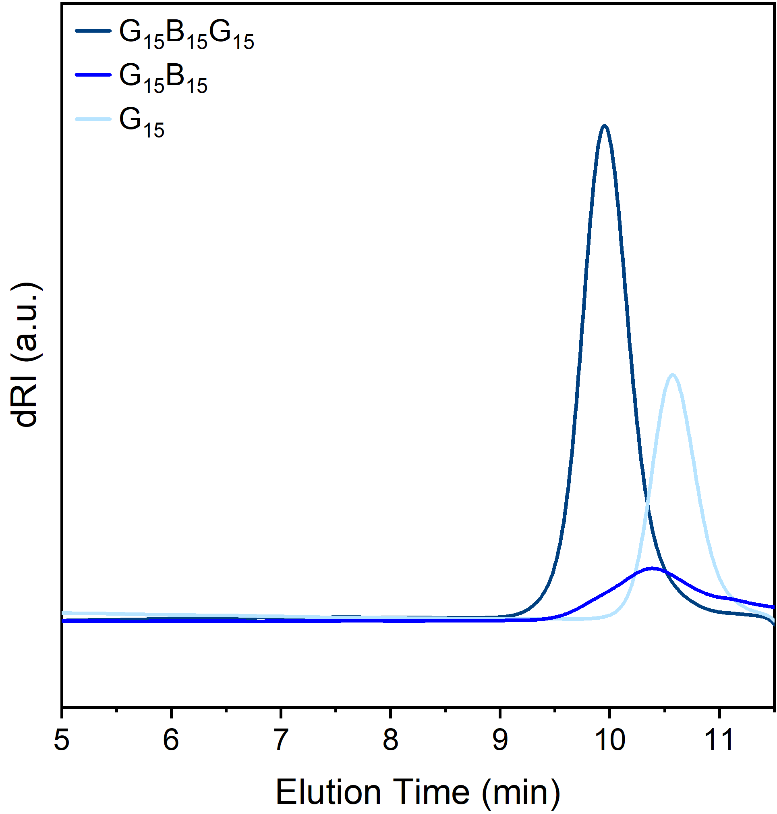

Figure S20. Organic SEC-MALS of polymers, ran_15_ (light blue trace), ran_15_beta_15_ (blue trace), and ran15beta15ran_15_ (navy trace), run using 0.05 M LiBr in DMF. The differential refractive index was not normalized.

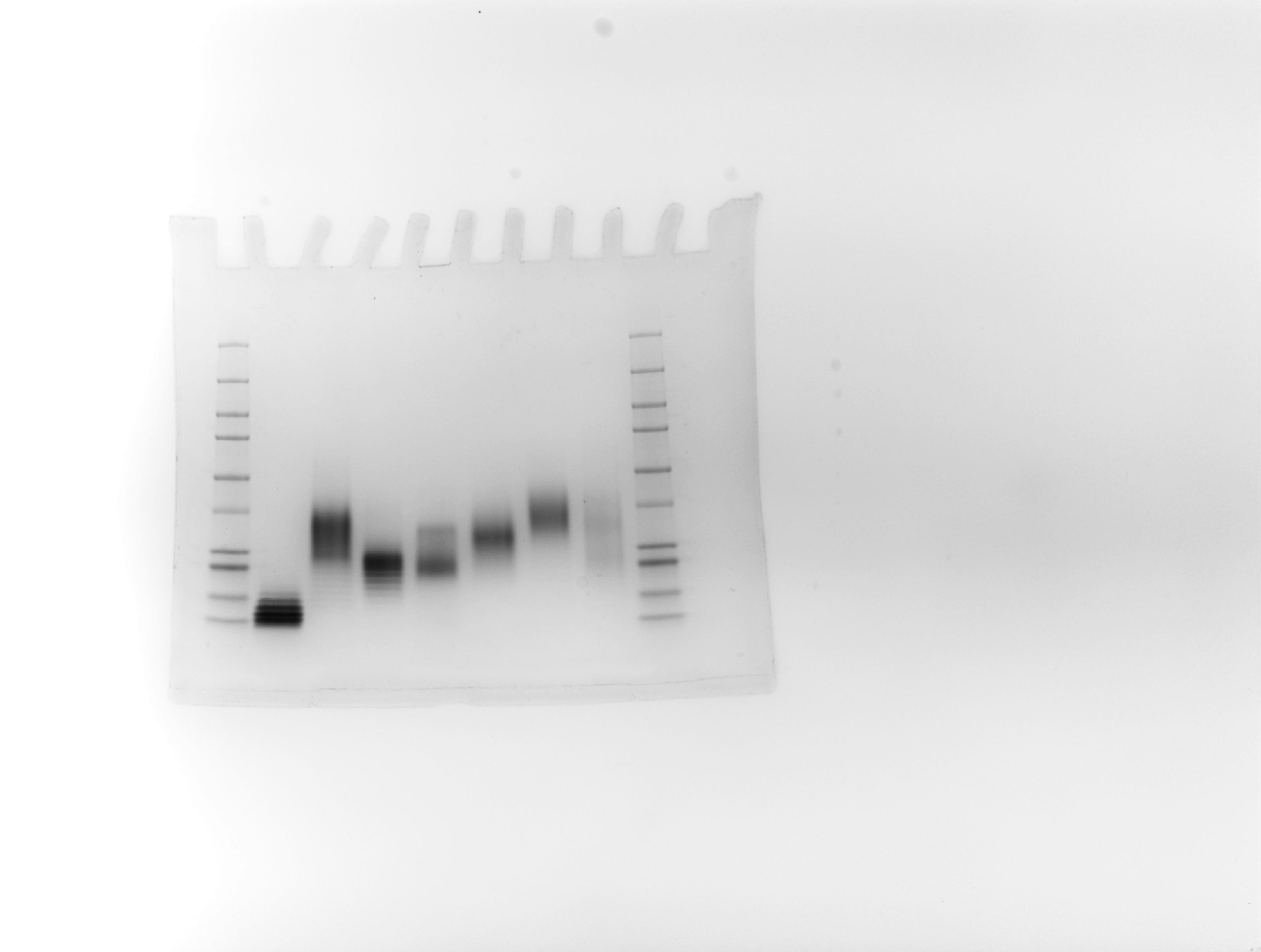

**beta_5_**

**beta_15_**

**10 kDa**

**15 kDa**

**20 kDa**

**25 kDa**

**37 kDa**

**50 kDa**

**75 kDa**

**100 kDa**

**150 kDa**

**250 kDa**

**G_15_beta_15_**

***ran-G_15_beta_15_***

**OEG_45_beta_15_**

Figure S21. SDS-PAGE gel of β-hairpin-derived peptide-brush polymers. Approximate average molecular weight was estimated from analysis of the band positions relative to the positions of the molecular weight ladders (far left and right).
Figure S22. SDS-PAGE gel of β-hairpin-derived peptide-brush polymers. Approximate average molecular weight was estimated from analysis of the band positions relative to the positions of the molecular weight ladders (middle lane).

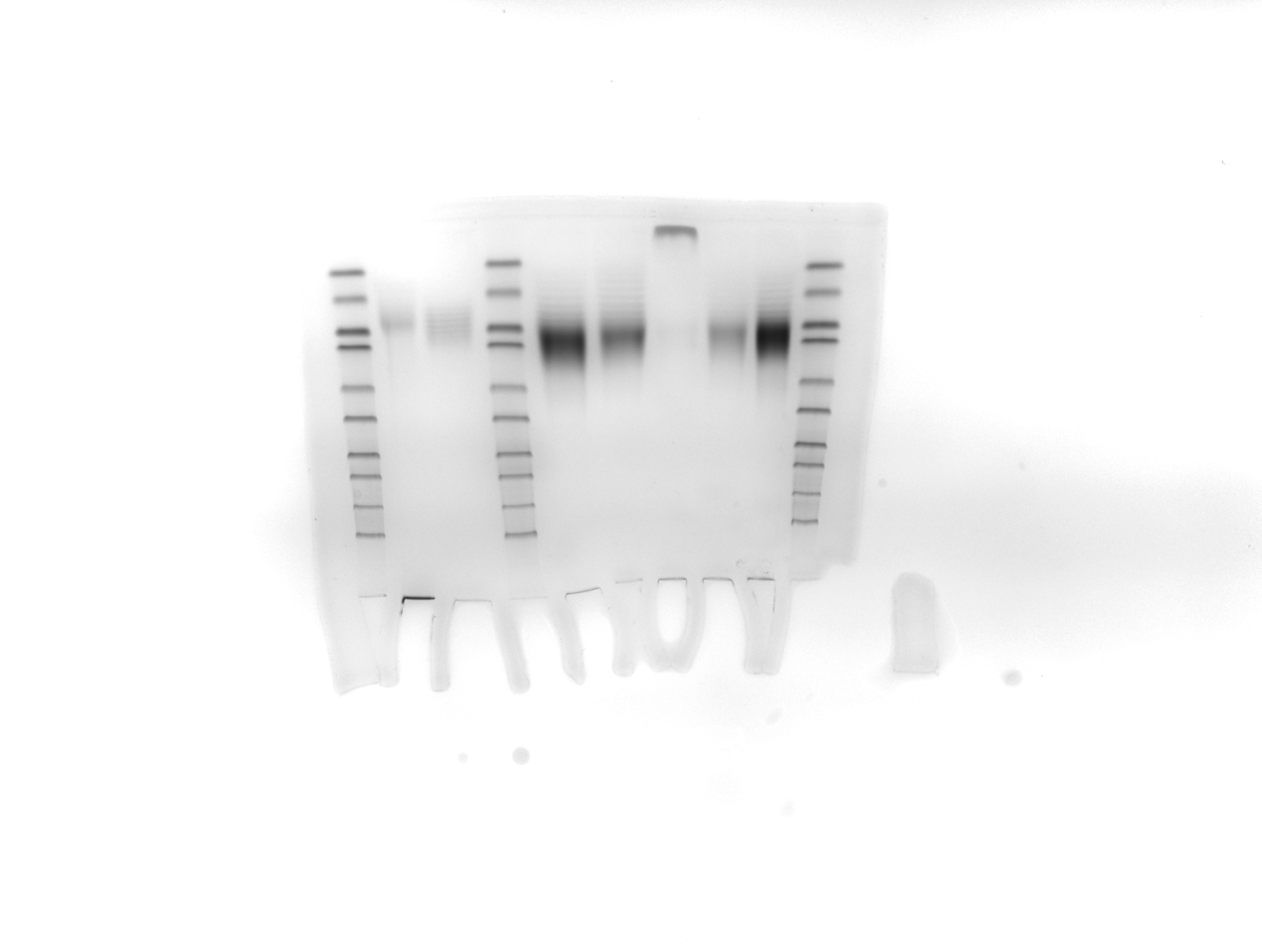

**NorVWWV_15_**

**10 kDa**

**15 kDa**

**20 kDa**

**25 kDa**

**37 kDa**

**50 kDa**

**75 kDa**

**100 kDa**

**150 kDa**

**250 kDa**

**NorWVVW_15_**

5. Circular Dichroism Spectroscopy

Circular dichroism (CD) spectra were obtained on a JASCO J-815 spectrometer equipped with a Peltier sample holder for temperature control in the Keck Biophysics Facility at Northwestern University. Samples were dissolved in 50 mM sodium acetate buffer at pH 4. These solutions were approximately 100 μM with respect to peptide monomer concentration, and placed into a 0.1 mm quartz cuvette (Hellma). Measurements were obtained using a scan speed of 10 nm/min, using a 1 nm bandwidth and 4 s response time. Background spectra of solvent were obtained prior to measurement of the sample solutions and subtracted from the sample spectra using JASCO Spectra Analysis. Spectra were not smoothed. Sample concentration was corrected using calculated extinction coefficients^11^ and absorbance data measured at 205 nm. Spectra were generally collected at 20 °C unless otherwise stated.

For temperature scans, sample temperature was ramped between 5 and 90 °C with steps of 1 °C/min while monitoring ellipticity at 229 nm. At endpoints of the temperature ramps (20 and 90 °C), full CD spectra were obtained.

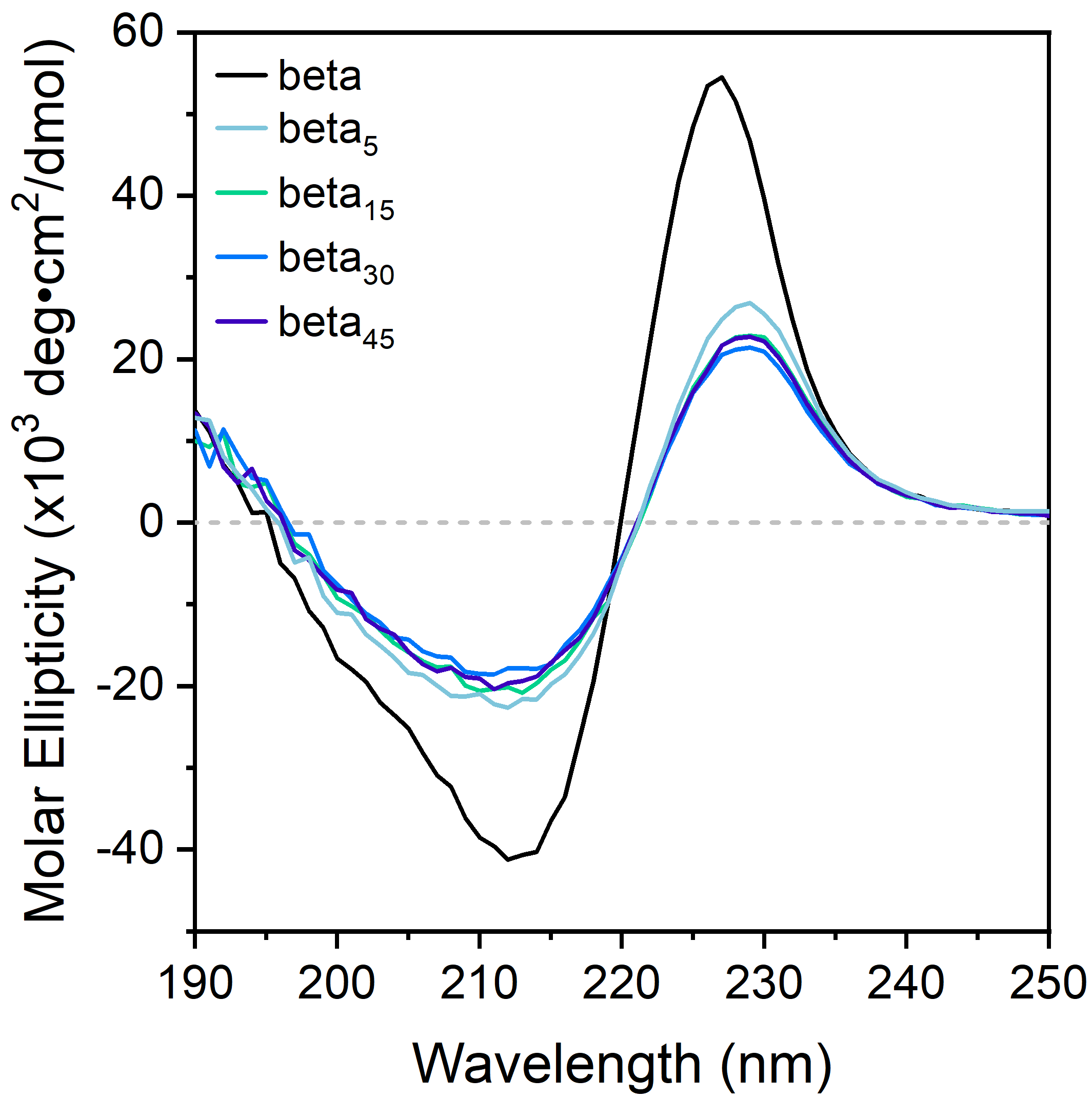

Figure S23. CD spectra of the beta monomer (black trace), beta_5_ (light blue trace), beta_15_ (green trace), beta_30_ (blue trace), and beta_45_ (dark blue trace), measured at 20 °C in 50 mM sodium acetate buffer.

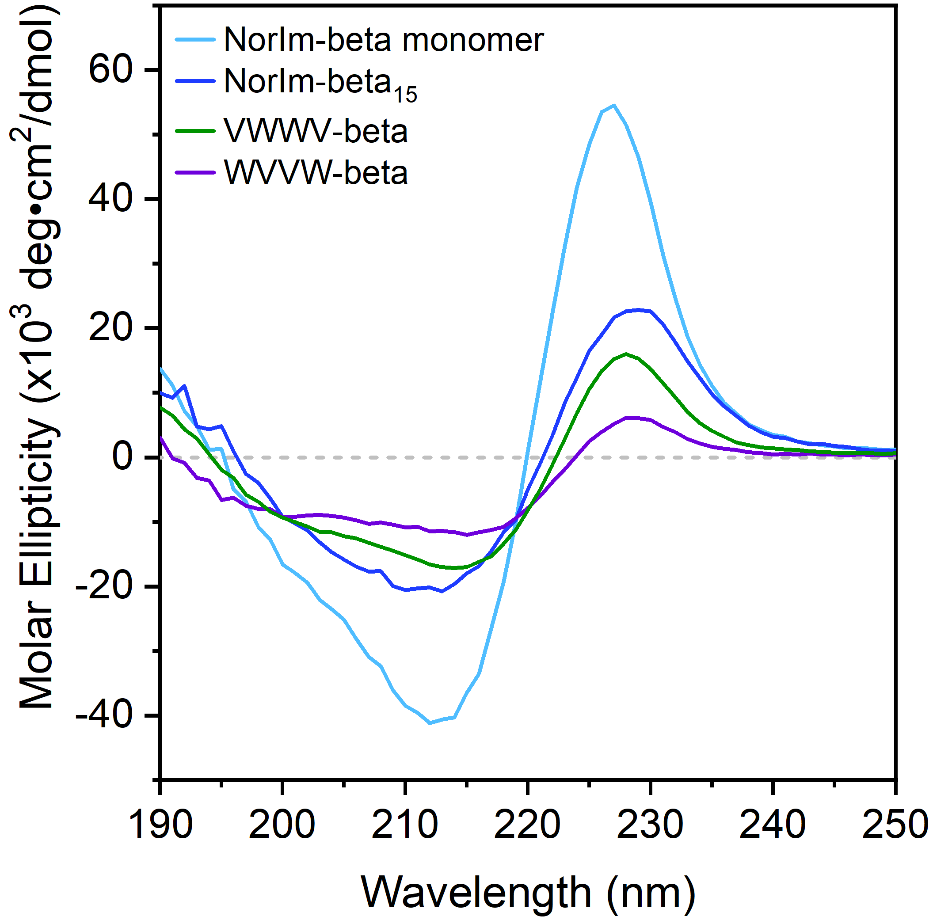

Figure S24. CD spectra of the peptide monomers beta (light blue trace), VWWV-beta (green trace), and WVVW-beta (purple trace), as well as the polymer beta_15_ (blue trace), measured at 20 °C in 50 mM sodium acetate buffer.

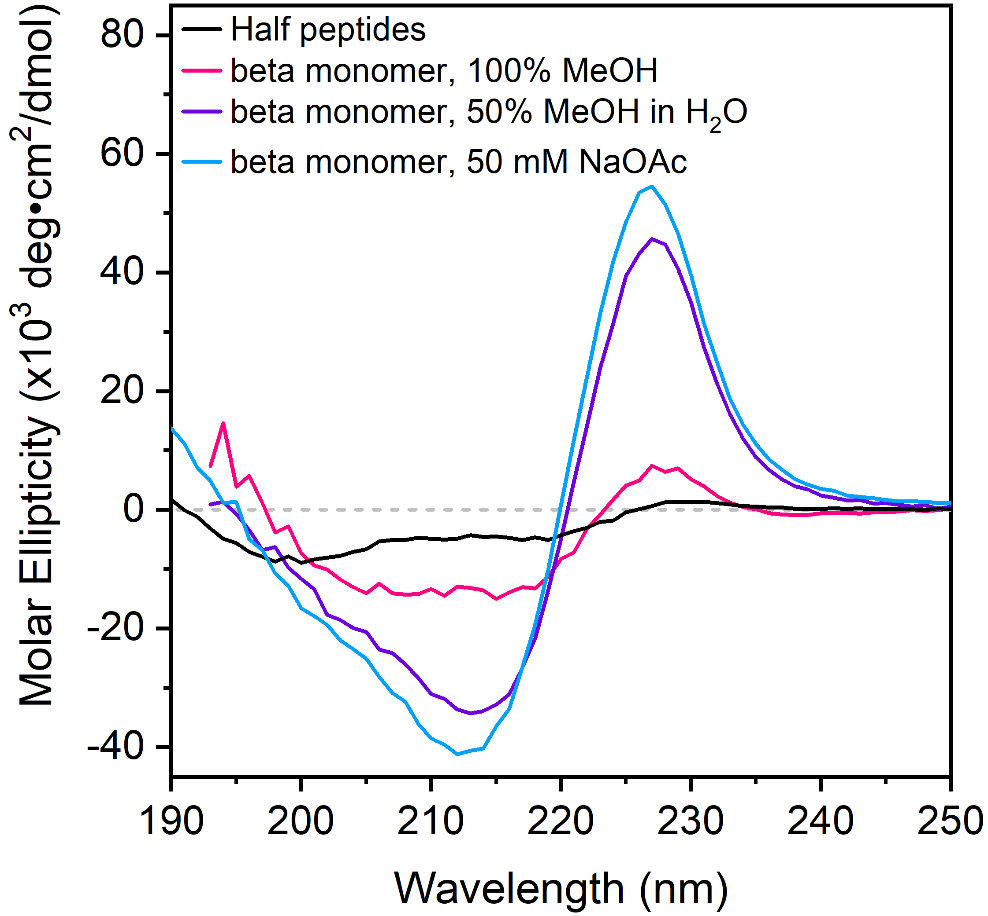

Figure S25. CD spectra of the beta monomer in 100% methanol (pink trace), 50% methanol in water (purple trace), and 50 mM sodium acetate buffer (light blue trace), measured at 20 °C in aqueous buffer. These spectra are shown plotted alongside the half peptides of the trpzip2 sequence (SWTWEN and GKWTWK; black trace).

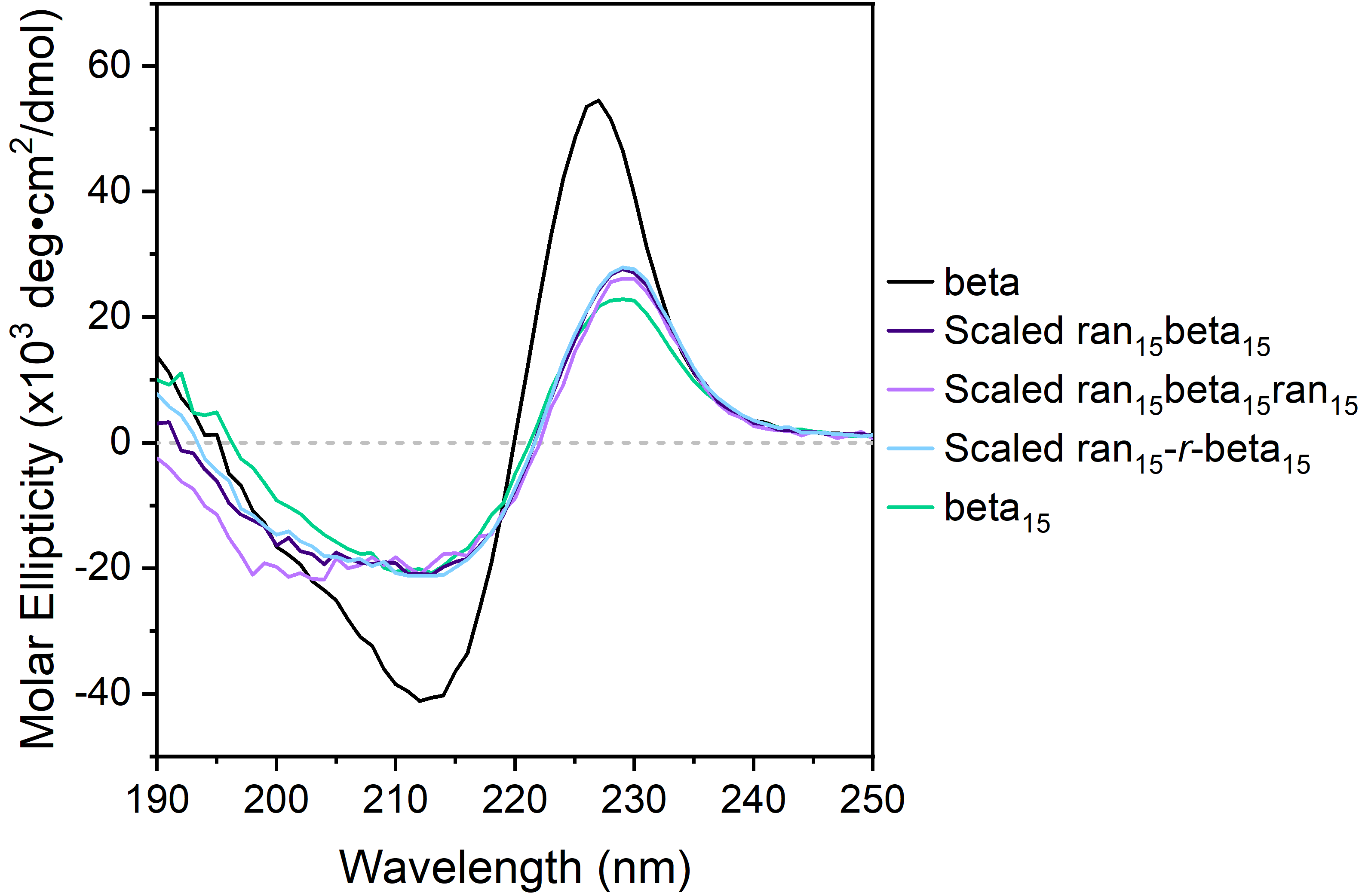

Figure S26. CD spectra of the beta monomer (black trace), beta_15_ (teal trace), scaled ran_15_beta_15_ (dark purple trace), scaled ran_15_beta_15_ran_15_ (light purple trace), and scaled ran_15_*-r-*beta_15_ran_15_ (light blue trace), measured at 20 °C in aqueous buffer. Scaling of intensity was performed according to the ratio of beta monomer in the copolymer.

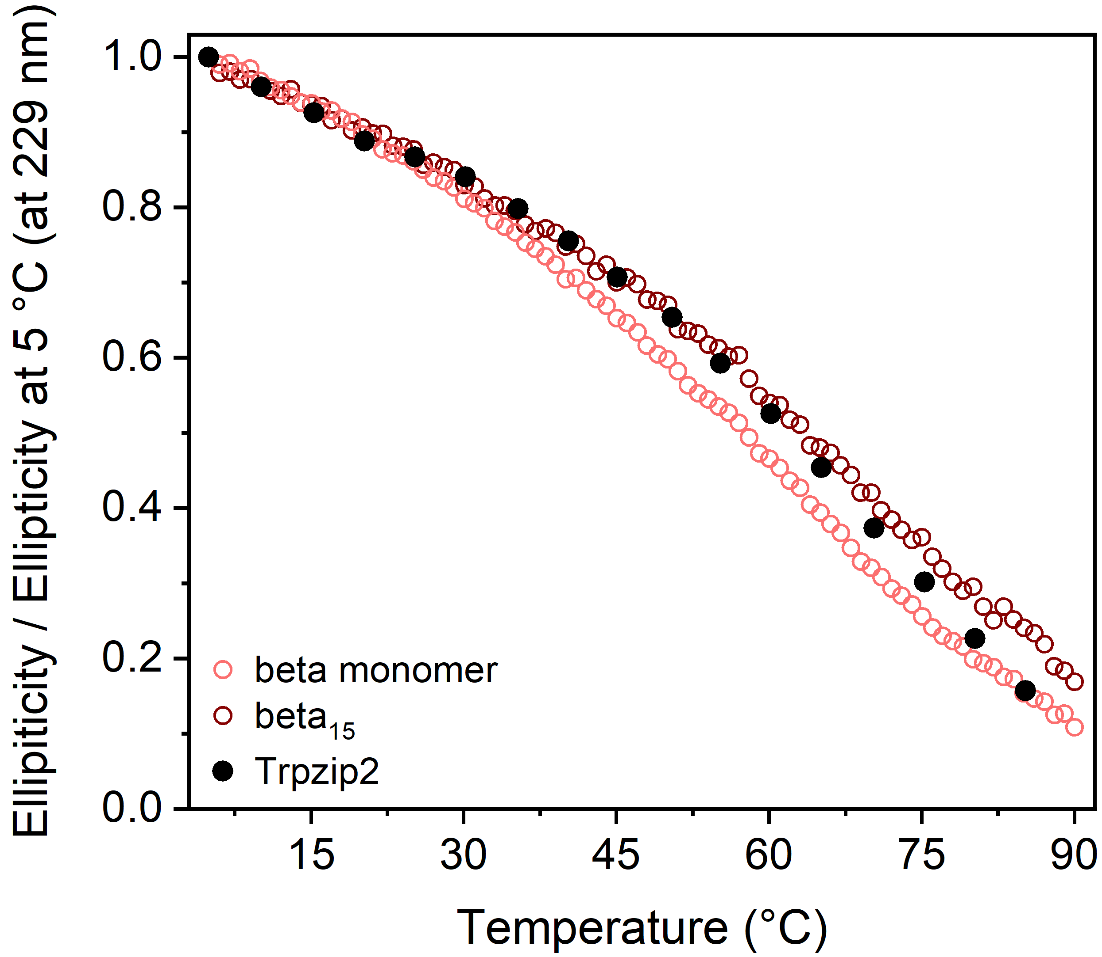

Figure S27. Ellipticity of beta monomer (pink symbols) and beta_15_ (maroon symbols) monitored at 229 nm during heating from 5 °C to 90 °C at 1 °C/min. These values were compared to those previously measured for trpzip2 (black symbols).^12^

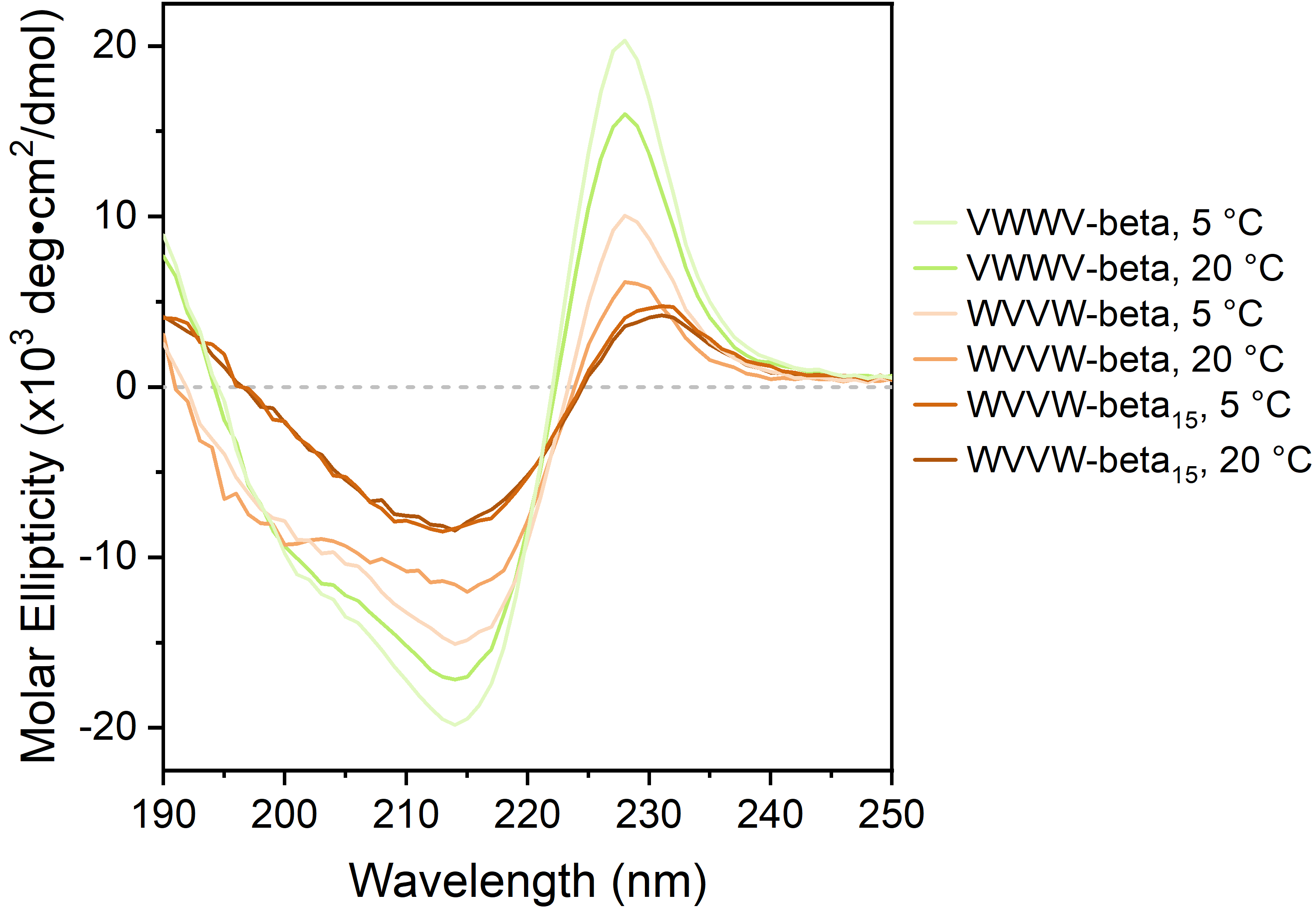

Figure S28. CD spectra of VWWV-beta at 5 °C (light green trace) and 20 °C (green trace), WVVW-beta at 5 °C (pale orange trace) and 20 °C (light orange trace), and WVVW-beta_15_ at 5 °C (orange trace) and 20 °C (darker orange trace).

6. Fourier Transform Infrared Spectroscopy

Samples of peptide monomers and polymers were dissolved in 10 mM HCl in H_2_O and lyophilized three times, to remove trace trifluoroacetic acid that might interfere with analysis of the amide IR bands.^13^ After the lyophilization process, the samples were dissolved in D_2_O (ca. 5 to 10 mg/mL) and then their Fourier transform infrared spectra were measured on a Thermo Scientific Nicolet iS50 infrared spectrometer. Solutions of samples (20 μL) were placed between two calcium fluoride plates (32 mm × 3 mm, Pike Technologies) with a 0.5 mm Teflon spacer, which were then put into a transmission cell and measured using 256 scans. Sample spectra were background-subtracted from a scan of D_2_O (20 μL) in the same setup.

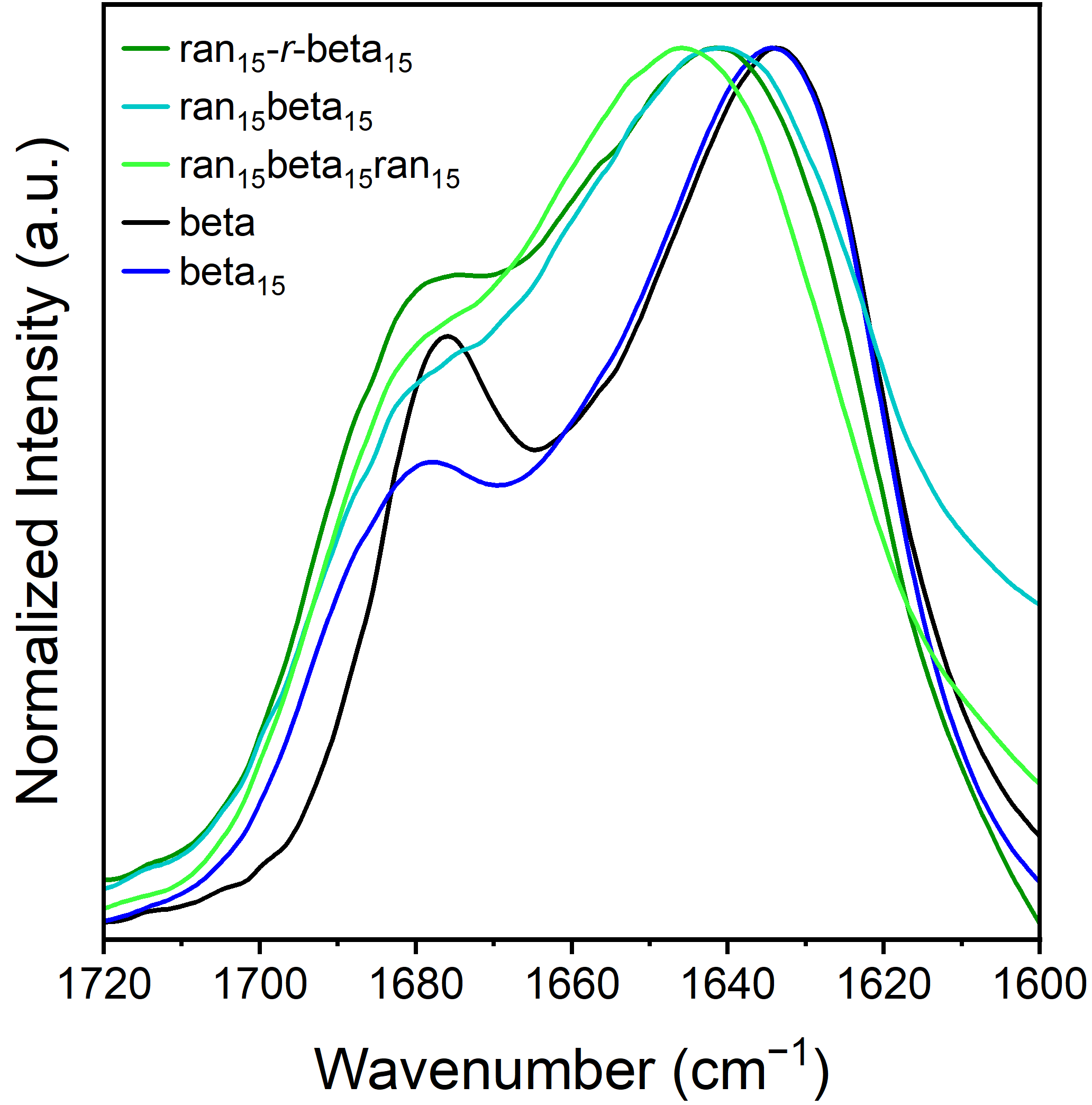

Figure S29. FTIR spectra of samples dissolved in D_2_O (5 to 10 mg/mL) of the beta monomer (black trace), beta_15_ polymer (blue trace), ran_15_-*r*-beta_15_ (green trace), ran_15_beta_15_ (teal trace), and ran_15_beta_15_ran_15_ (light green trace).

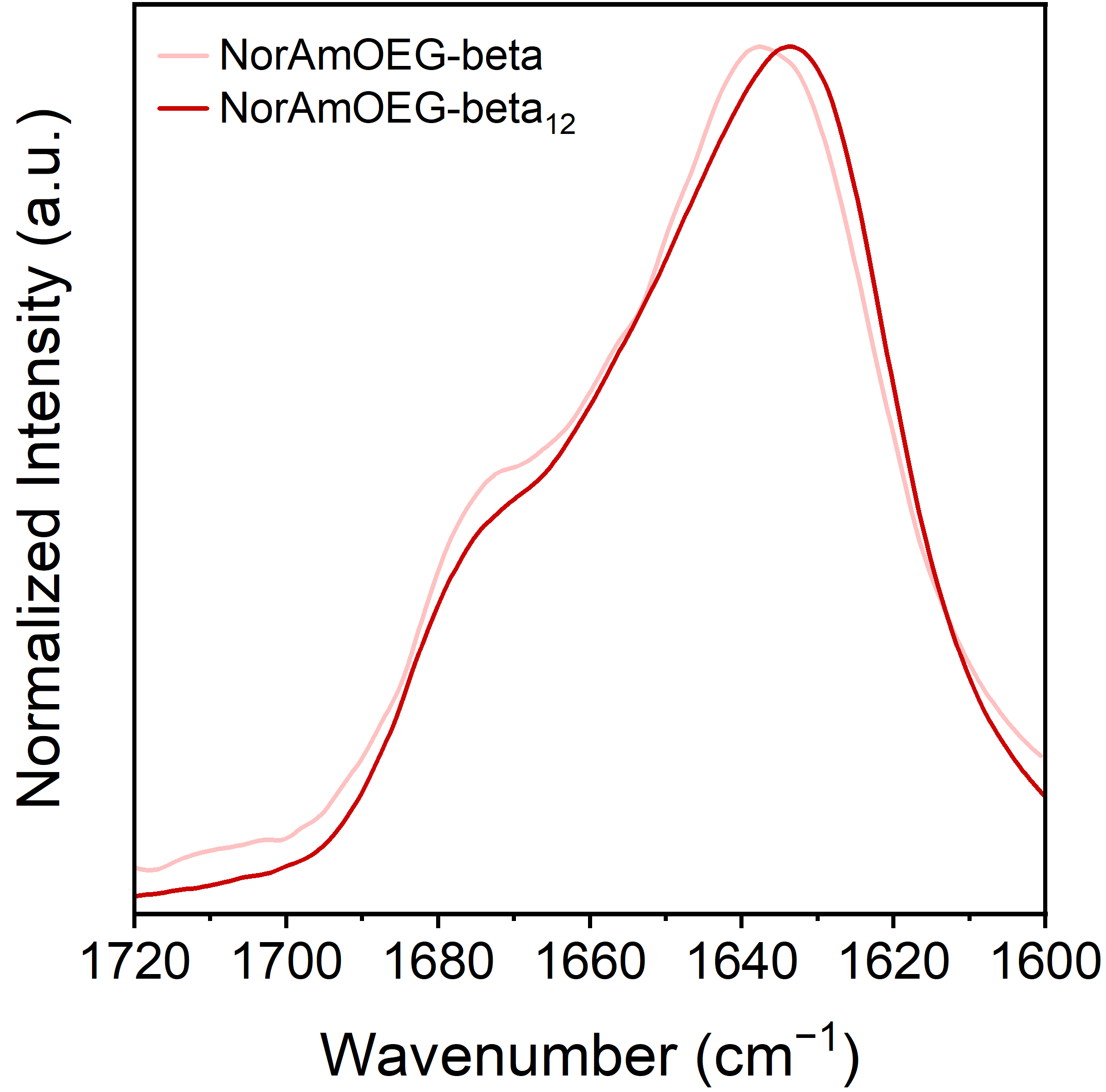

Figure S30. FTIR spectra of samples dissolved in D_2_O of NorAmOEG-beta (pink trace) and NorAmOEG-beta_12_ (red trace).

7. Simulations

**Structure and parametrization**

CHARMM36m was used for parameterization of the peptide sequence, but the NorAha group (hereon referred to as the ligand) as well as the bonds and dihedrals involving NorAha were parametrized using CGENFF. To parametrize brush polymer structures, the 3D model is first built in VMD using Molefacture, starting from an accurate ChemDraw structure. The structure is saved as both .mol2 and .pdb files. Since Molefacture assigns atom names rather than element types, atom names in the .mol2 file must be manually corrected before submitting to CHARMM-GUI’s Ligand Reader. CHARMM-GUI generates topology (lig.rtf) and parameter (lig.prm) files. A ligand-only version of the topology file is created (lig_only.rtf) by removing non-ligand atoms and irrelevant bonded terms to avoid artifacts during topology generation with PSFGEN. The structure is then cleaned and renumbered. Ligand atom names are updated to match CHARMM-GUI output, enabling separation into ligand and peptide .pdb files. A connect patch is written to define the bond between peptide and ligand, ensuring consistent atom types. After PSFGEN is run, the output is visually inspected for structural integrity. Running ParmEd on the generated .psf and .pdb typically reveals missing bonded parameters between protein and ligand atoms. These are resolved by mapping protein atom types to CGENFF equivalents and extracting the appropriate bonded terms from the CGENFF force field. Custom parameters are stored in a new parameter file (lig_self_defined.prm) and sourced during ParmEd conversion to generate final AMBER-compatible rst7 and parm7 files. Solvent (TIP3P) and ions (Na+ and Cl−, for neutralization) are added afterward to minimize errors during parametrization.

**Molecular dynamics (MD) simulations**

**Monomer system**

Molecular dynamics simulations were performed using OpenMM 7.7.0. A minimization protocol was followed by an NVT ensemble simulation performed for 2 ns. The system was heated from 5 K to 300 K. Protein backbone atoms were restrained during the NVT simulation using a spring constant of 1.0 kcal/mol·Å2. An NPT ensemble simulation was then performed for 4 ns: the first 2 ns with restrained backbone atoms, followed by 2 ns while removing restraints in 100 steps by reducing the value of the spring constant from 1.0 kcal/mol·Å2 to zero. A further 2 ps of NPT ensemble simulation was performed without restraints. This was followed by an additional equilibration step using an NPT simulation without restraints for 2 ns. Production simulations were carried out in the NPT ensemble for a cumulative simulation time from all replicates of ∼1.736 ms. Folding@Home distributed computing was used for the production simulations. Pre-production simulations and setup were performed on the NVIDIA RTX 4090 GPU.

**Polymer systems**

Molecular dynamics simulations were performed using OpenMM 7.7.0. A minimization protocol was followed by an NVT ensemble simulation performed for 10 ns. The system was heated from 5 K to 300 K. Protein backbone atoms were restrained during the NVT simulation using a spring constant of 1.0 kcal/mol·Å^2^. An NPT ensemble simulation was then performed for 20 ns: the first 10 ns with restrained backbone atoms, followed by 10 ns while removing restraints in 100 steps by reducing the value of the spring constant from 1.0 kcal/mol·Å^2^ to zero. A further 2 ps of NPT ensemble simulation was performed without restraints. This was followed by an additional equilibration step using an NPT simulation without restraints for 50 ns. Production simulations were carried out in the NPT ensemble for a cumulative simulation time from all replicates of ∼100 μs for each polymer system. Folding@Home distributed computing was used for the production simulations. Pre-production simulations and setup were performed on the NVIDIA RTX 4090 GPU.

**Simulation Parameters**

A Langevin integrator was used with a temperature of 300 K, a friction coefficient of 2.8284 ps⁻¹, and a time step of 2 fs. Constraints were applied to all bonds involving hydrogen atoms, and water molecules were kept rigid. Nonbonded interactions were treated using the Particle Mesh Ewald (PME) method with a cutoff distance of 1.2 nm and an Ewald error tolerance of 0.0005. A Monte Carlo barostat was added to maintain a pressure of 1 atm at 300 K. The constraint tolerance was set to 1 × 10−6. Trajectory data were saved every 50,000 steps (100 ps).

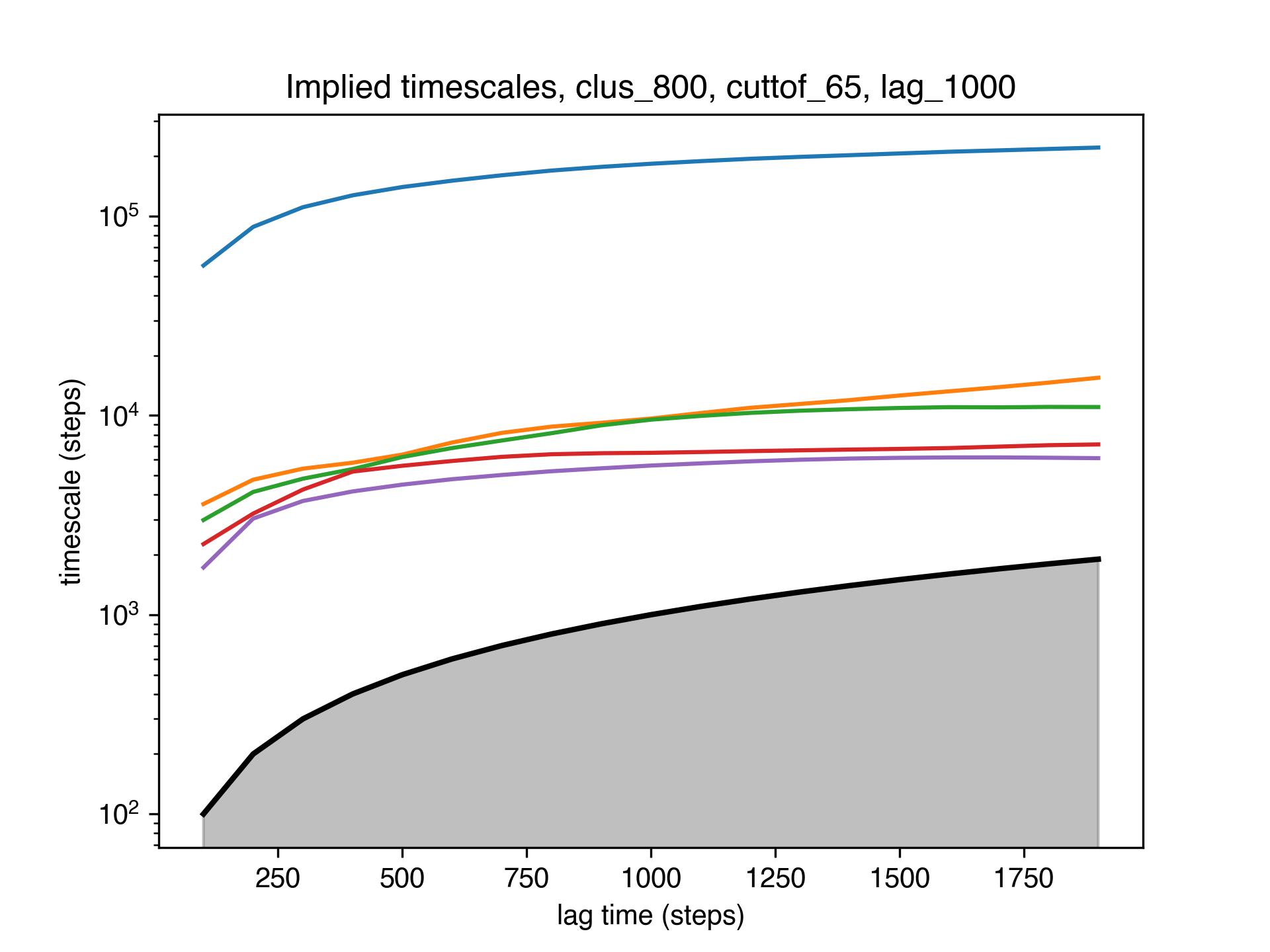

Figure S31. Implied timescale convergence plot for the monomer system, showing the slowest timescales. The Markov state model was built using 800 clusters, tICA cutoff of 65% and lag time of 100 ns (1000 steps).

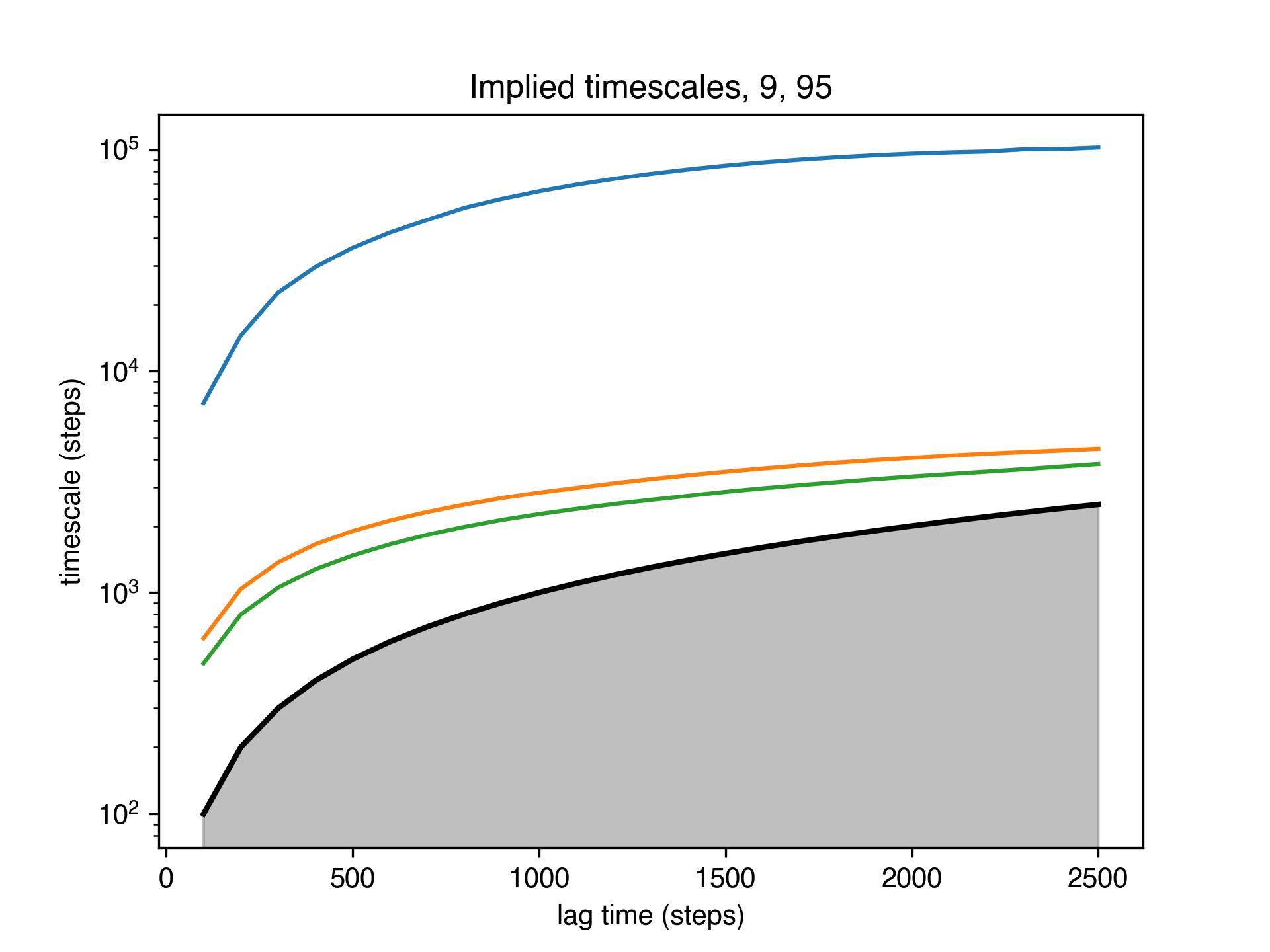

Figure S32. Implied timescale convergence plot for the asparagine-linked polymer system, showing the slowest timescales. The Markov state model was built using 9 clusters, tICA cutoff of 95% and lag time of 180 ns (1800 steps).

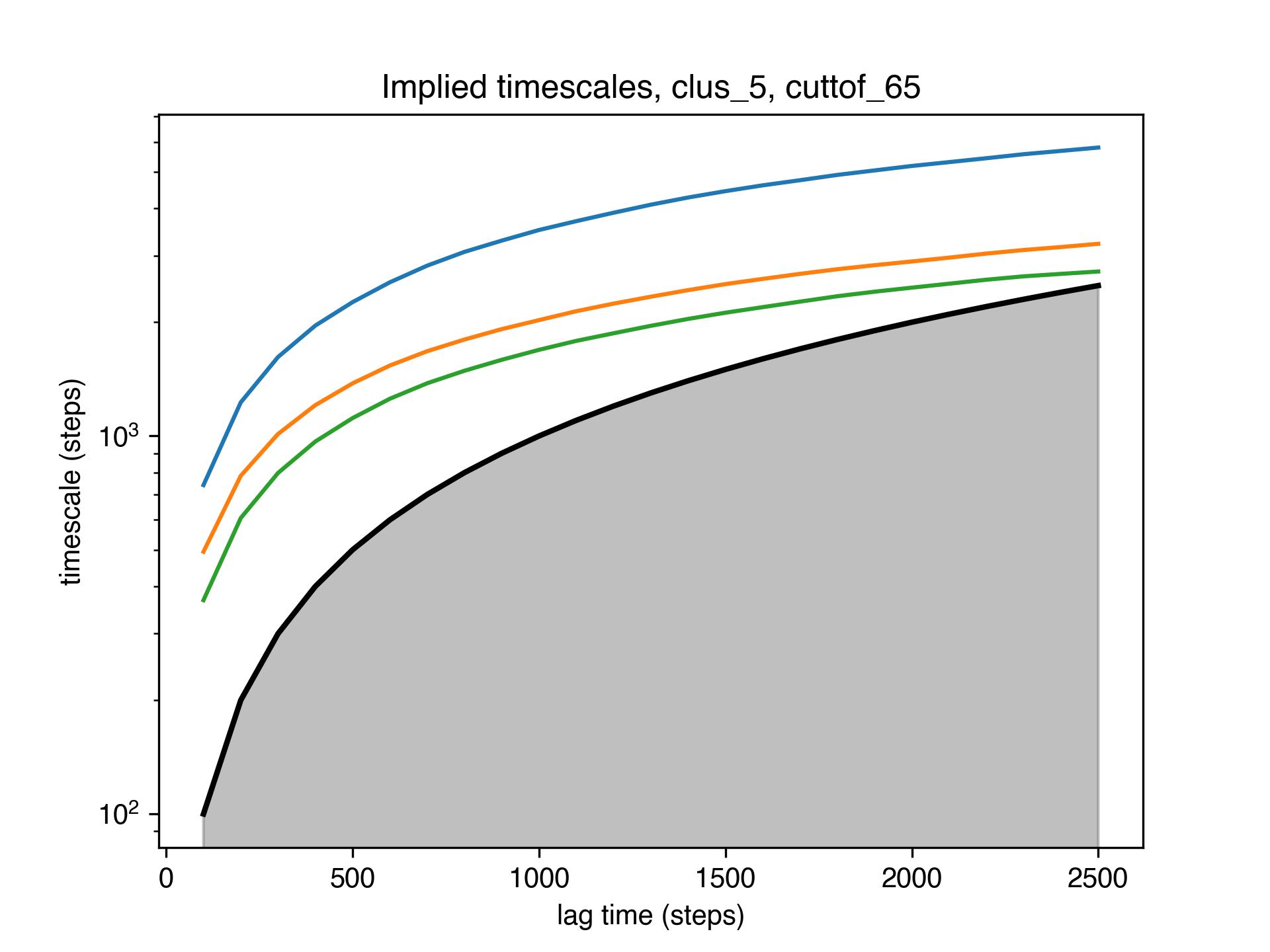

Figure S33. Implied timescale convergence plot for the serine-linked polymer system, showing the slowest timescales. The Markov state model was built using 5 clusters, tICA cutoff of 65% and lag time of 200 ns (2000 steps).

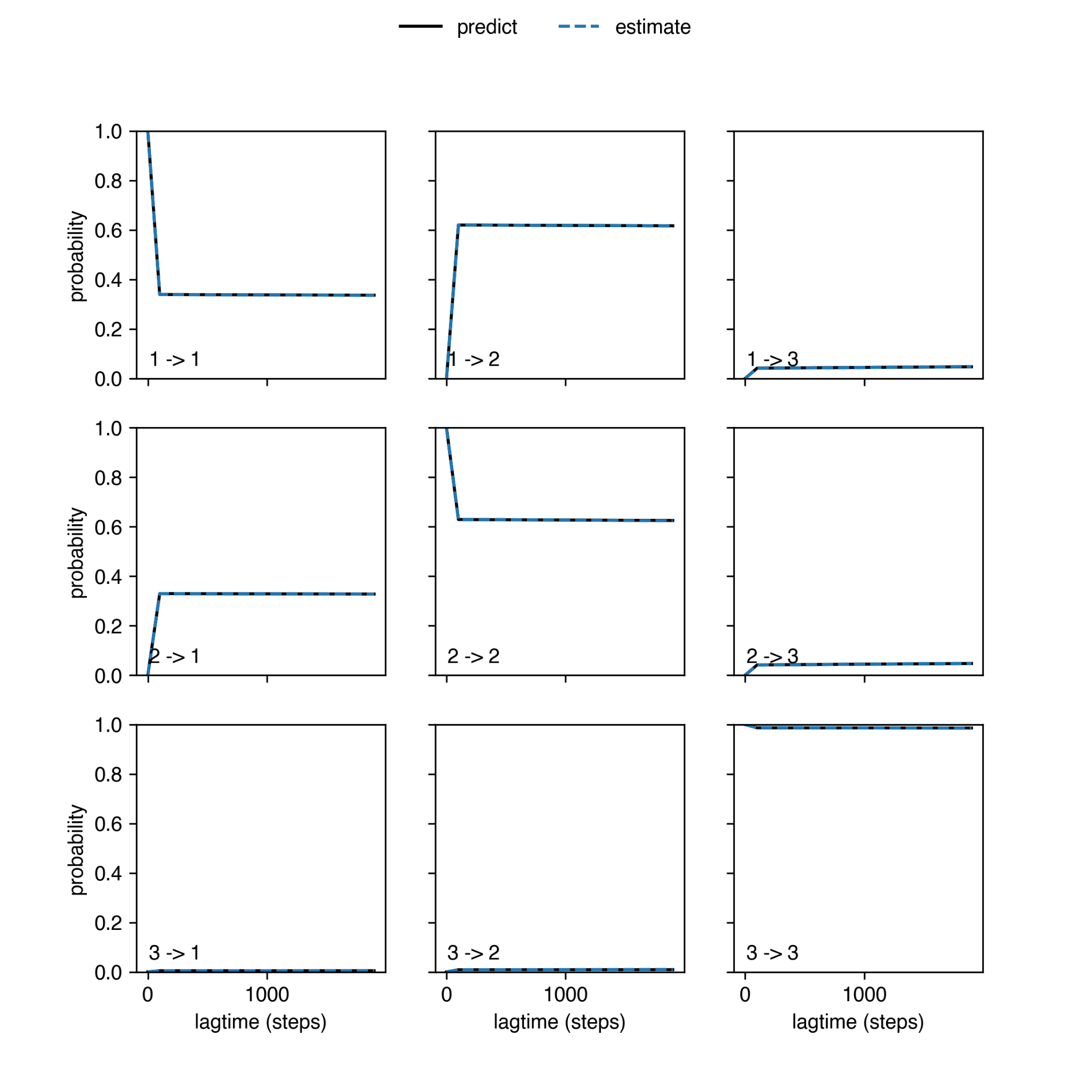

Figure S34. Chapman–Kolmogorov test for the monomer system with three metastable states.

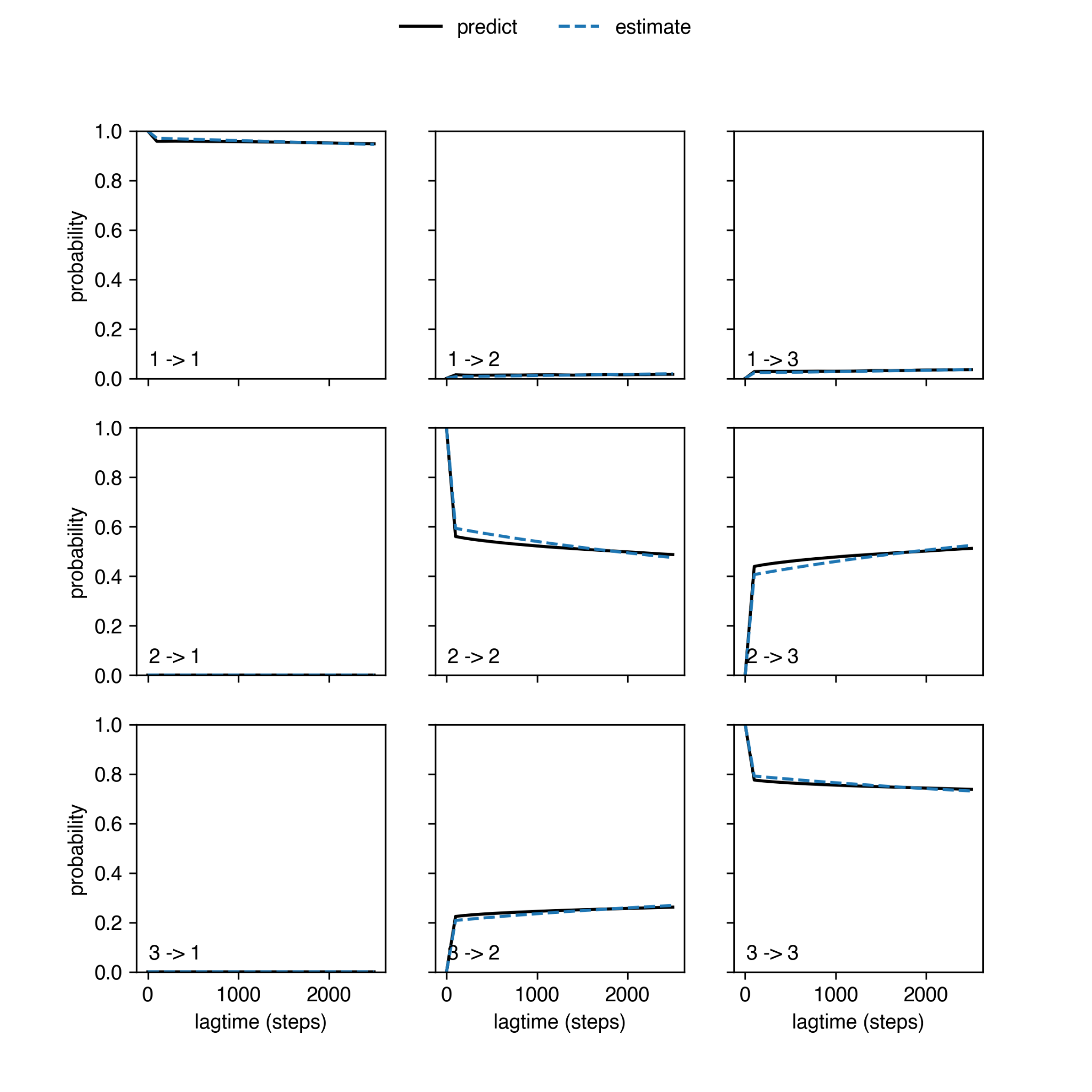

Figure S35. Chapman–Kolmogorov test for the asparagine-linked polymer system with three metastable states.

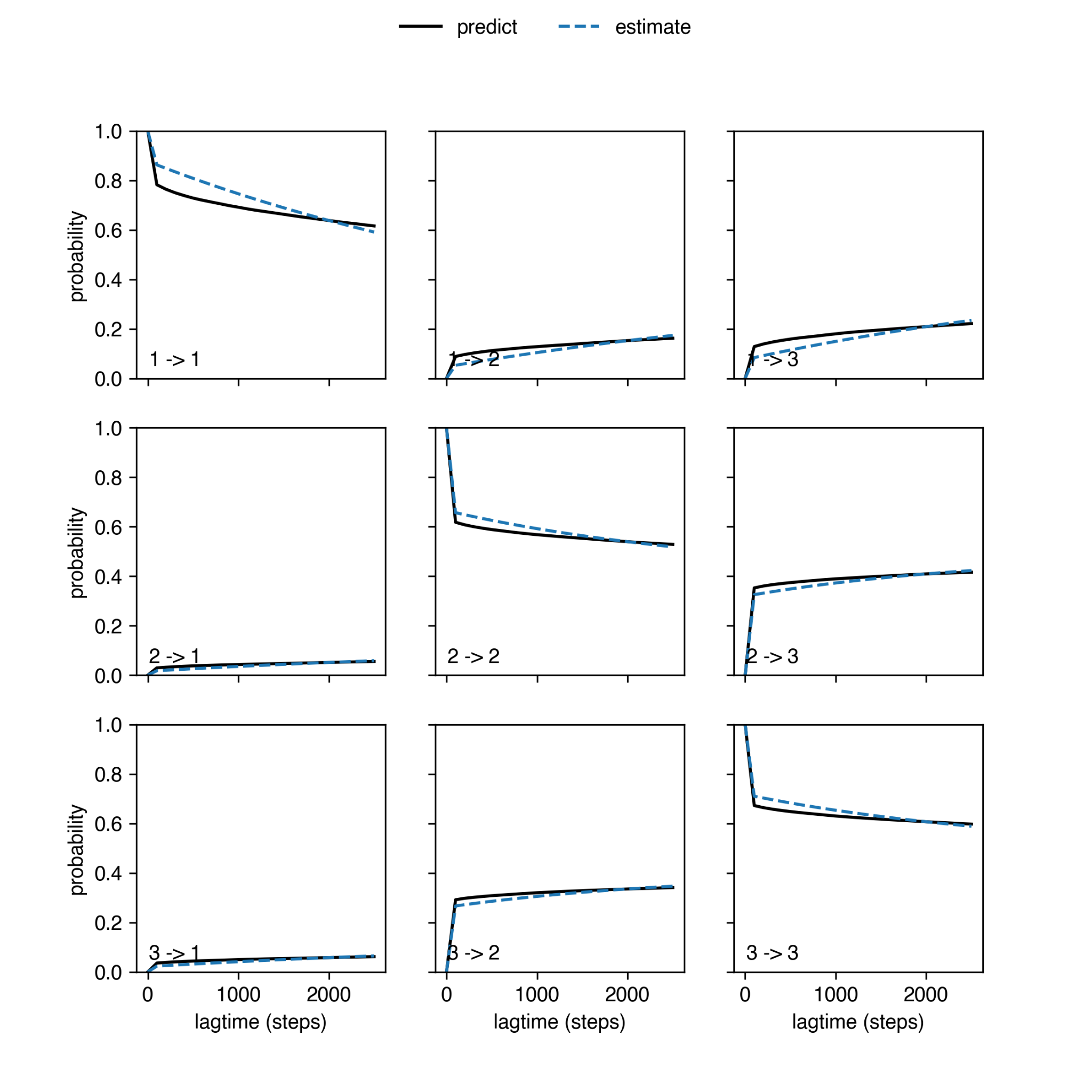

Figure S36. Chapman–Kolmogorov test for the serine-linked polymer system with three metastable states.

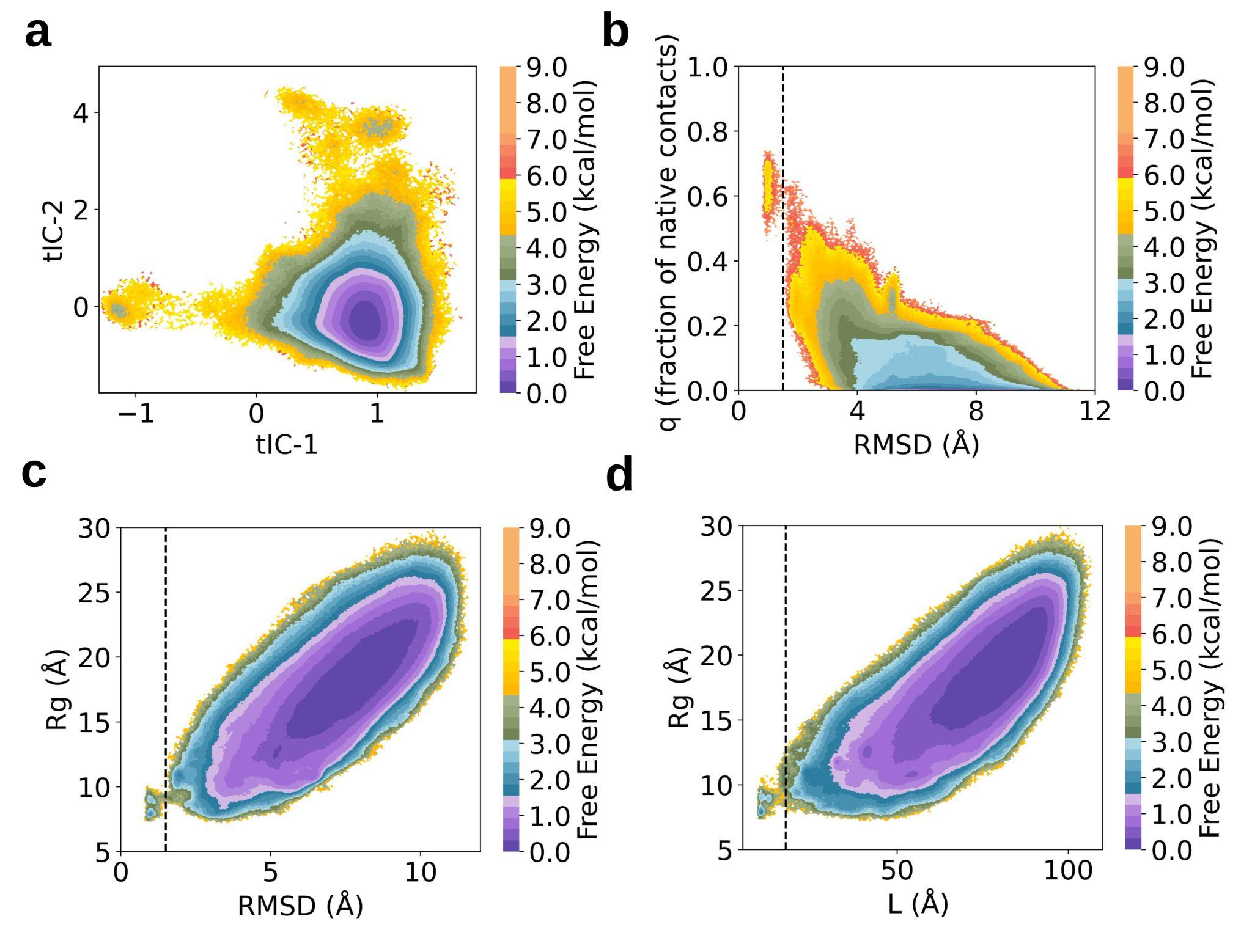

Figure S37. Free energy maps constructed from simulations of asparagine-linked polymer system, projected onto various features. **(a)** tICA projection **(b)** Root-mean-square deviation (RMSD) vs. fraction of native contacts **(c)** RMSD vs. radius of gyration of the tryptophan pairs **(d)** Sum of native hydrogen bond distances vs. radius of gyration of the tryptophan pairs.

8. Small-Angle X-ray Scattering

SAXS experiments were conducted at Beamline 16-ID of the National Synchrotron Light Source II, Brookhaven National Laboratory.^14–16^ Samples were dissolved in buffered solutions (sodium acetate buffer) in concentrations ranging from 1.25 to 5 mg/mL, dialyzed for two days against the buffer solution using 3.5 kDa molecular weight cutoff (MWCO) dialysis cassettes, filtered through 0.22 μm hydrophilic PTFE membrane filters, shipped to the beamline in sealed PCR plates, and then loaded into sample holders for measurement in a flow-through cell at the beamline station.^14,16^ The wavelength used for data collection ranges from 0.8172 Å to 0.8202 Å and is indicated in figure captions and in Table S1. Sample data were collected simultaneously on Pilatus 1M and Pilatus 900K detectors to capture an approximate *q* range of 0.006 Å^−1^ to 3.2 Å^−1^. Data processing and background subtraction were performed using Python as previously detailed.^14^

Background-subtracted patterns were used to construct Kratky and Guinier plots. Specifically, Guinier plots were used to extract experimental radii of gyration, *R*_g_, and the intensity at zero scattering angle, *I*_0_, using the program ATSAS.^17^ These values are reported in Table S1. Additionally, the values obtained from this analysis were used to generate Kratky and normalized Kratky plots of the corresponding data, which are presented in the manuscript and below (Figures 5 and S41). Fitting of the power law, Guinier–Porod,^18,19^ and flexible cylinder models^20,21^ was performed using SasView.^22^ These models are frequently used for similar bottle-brush polymers and provide detail into their conformations in solution.^23–28^ For example, shorter polymers often collapse into compact, globular conformations, which typically have a dimension (or shape) parameter *s* of 0 and a Porod exponent of about 4, whereas more anisotropic systems will yield dimension parameters closer to 1.

Data for samples of all concentrations were generally examined for signs of aggregation or interparticle repulsion, and generally, the data of the intermediate concentration, 2.5 mg/mL, was chosen due to not exhibiting such effects while maintaining high intensities and signal-to-noise.

Table S1. Table of structural parameters of all polymers. All *R*_g_ and *I*_0_ values were derived from samples of concentrations 2.5 mg polymer per mL of solvent, dissolved in 100 mM NaOAc buffer, pH 4.

| **Sample Name** | ***λ* (Å)** | ***R*_g_ (nm), from Guinier plot fit** | ***I*_0_** | **Fidelity** |
| --- | --- | --- | --- | --- |
| beta_5_ | 0.8172 | 2.45 ± 0.01 | 4.78 ± 0.02 | 1 |
| beta_15_ | 0.8189 | 3.40 ± 0.04 | 1.81 ± 0.02 | 0.78 |
| ran_15_beta_15_ | 0.8172 | 2.578 ± 0.009 | 15.30 ± 0.04 | 0.99 |
| ran_15_beta_15_ran_15_ | 0.8172 | 2.96 ± 0.01 | 23.74 ± 0.08 | 0.73 |
| ran_15_-*r*-beta_15_ | 0.8172 | 2.78 ± 0.01 | 20.27 ± 0.05 | 0.99 |
| OEG_45_-*r*-beta_15_ | 0.8172 | 3.04 ± 0.01 | 6.54 ± 0.02 | 0.99 |
| Beta_3_-OEG_45_ | 0.8189 | 3.02 ± 0.04 | 1.33 ± 0.01 | 0.94 |

Table S2. Table of structural parameters of polymers derived from the β-hairpin peptide sequence. Specifically, we report the radius of gyration *R*_g_, the dimension variable *s*, and the Porod exponent *d* determined from Guinier–Porod model^18,19^ fits performed in SasView,^22^ along with the goodness-of-fit values.

| **Sample Name** | ***R*_g_ (nm)** | ***s*** | ***d* (Porod exponent)** | **χ^2^** |
| --- | --- | --- | --- | --- |
| beta_5_ | 1.90 ± 0.02 | 0.22 ± 0.01 | 3.7 ± 0.1 | 1.65 |
| beta_15_ | 1.91 ± 0.02 | 0.25 ± 0.01 | 4.1 ± 0.1 | 1.46 |
| ran_15_beta_15_ | 2.29 ± 0.07 | 0.091 ± 0.004 | 4.21 ± 0.09 | 6.02 |
| ran_15_beta_15_ran_15_ | 2.76 ± 0.07 | 0.056 ± 0.004 | 4.27 ± 0.03 | 7.26 |
| ran_15_-*r*-beta_15_ | 2.533 ± 0.06 | 0.065 ± 0.004 | 4.28 ± 0.04 | 7.85 |
| OEG_45_-*r*-beta_15_ | 2.38 ± 0.01 | 0.204 ± 0.006 | 3.59 ± 0.03 | 4.25 |
| beta_3_OEG_45_ | 2.20 ± 0.03 | 0.26 ± 0.02 | 2.82 ± 0.07 | 1.21 |

Table S3. Table of structural parameters of β-hairpin peptide-based polymers. P1 and P2 designate power law coefficients derived from fitting the power law model as implemented in SasView, with the respective q range of the fit and goodness-of-fit value designated below the coefficient.^22^

| **Sample Name** | **P1** | **P2** |
| --- | --- | --- |
| beta_5_ | 1.029 ± 0.006  0.007 to 0.03 Å^-1^; χ^2^ 0.86 | 3.500 ± 0.004  0.1 to 0.3 Å^-1^; χ^2^ 1.97 |
| beta_15_ | 0.00002 ± 0.0000002  0.01 to 0.03 Å^-1^; χ^2^ 0.66 | 3.628 ± 0.004  0.1 to 0.3 Å^-1^; χ^2^ 1.28 |
| ran_15_beta_15_ | 0.0110 ± 0.0001  0.008 to 0.025 Å^-1^; χ^2^ 1.36 | 4.260 ± 0.002  0.1 to 0.2 Å^-1^; χ^2^ 3.76 |
| ran_15_beta_15_ran_15_ | 0.000329 ± 0.000002  0.007 to 0.025 Å^-1^; χ^2^ 2.88 | 3.53 ± 0.04  0.08 to 0.17 Å^-1^; χ^2^ 3.14 |
| ran_15_-*r*-beta_15_ | 0.000460 ± 0.000005  0.008 to 0.025 Å^-1^; χ^2^ 1.82 | 3.243 ± 0.001  0.08 to 0.18 Å^-1^; χ^2^ 7.05 |
| OEG_45_-*r*-beta_15_ | 0.000430 ± 0.000003  0.007 to 0.02 Å^-1^; χ^2^ 0.37 | 3.190 ± 0.001  0.08 to 0.22 Å^-1^; χ^2^ 3.82 |
| beta_3_OEG_45_ | 0.00415 ± 0.00002  0.007 to 0.035 Å^-1^; χ^2^ 0.84 | 3.540 ± 0.006  0.1 to 0.35 Å^-1^; χ^2^ 0.58 |

Table S4. Table of structural parameters of peptide-based polymers derived from the β-hairpin peptide monomer. Specifically, we report the parameters determined using the flexible cylinder model^20,21^ as implemented in SasView,^22^ with *L* representing the length of the flexible cylinder, the Kuhn length of the flexible cylinder, *R* as the radius of the flexible cylinder, as well as the polydispersity of *R* and the goodness-of-fit parameter χ^2^.

| **Sample Name** | ***L* (nm)** | **Kuhn Length (nm)** | ***R* (nm)** | **Polydispersity of *R*** | **χ^2^** |
| --- | --- | --- | --- | --- | --- |
| beta_5_ | 11.977 ± 0.004 | 3.0 ± 0.2 | 1.46 ± 0.02 | 0.15 ± 0.04 | 0.78 |
| beta_15_ | 17.01 ± 0.02 | 1.7 ± 0.2 | 1.46 ± 0.06 | 0.13 ± 0.04 | 0.87 |
| ran_15_beta_15_ | 10.996 ± 0.001 | 2.7 ± 0.2 | 1.78 ± 0.01 | 0.24 ± 0.02 | 1.14 |
| ran_15_beta_15_ran_15_ | 12.0 ± 0.2 | 3.002 ± 0.001 | 2.088 ± 0.009 | 0.28 ± 0.01 | 1.53 |
| ran_15_-*r*-beta_15_ | 11.5 ± 0.2 | 2.8675 ± 0.001 | 1.920 ± 0.008 | 0.25 ± 0.01 | 2.07 |
| OEG_45_-*r*-beta_15_ | 20.24 ± 0.01 | 2.0 ± 0.1 | 1.76 ± 0.05 | 0.17 ± 0.04 | 0.77 |
| beta_3_OEG_45_ | 15.689 ± 0.001 | 3.9 ± 0.3 | 1.40 ± 0.06 | 0.29 ± 0.06 | 0.68 |

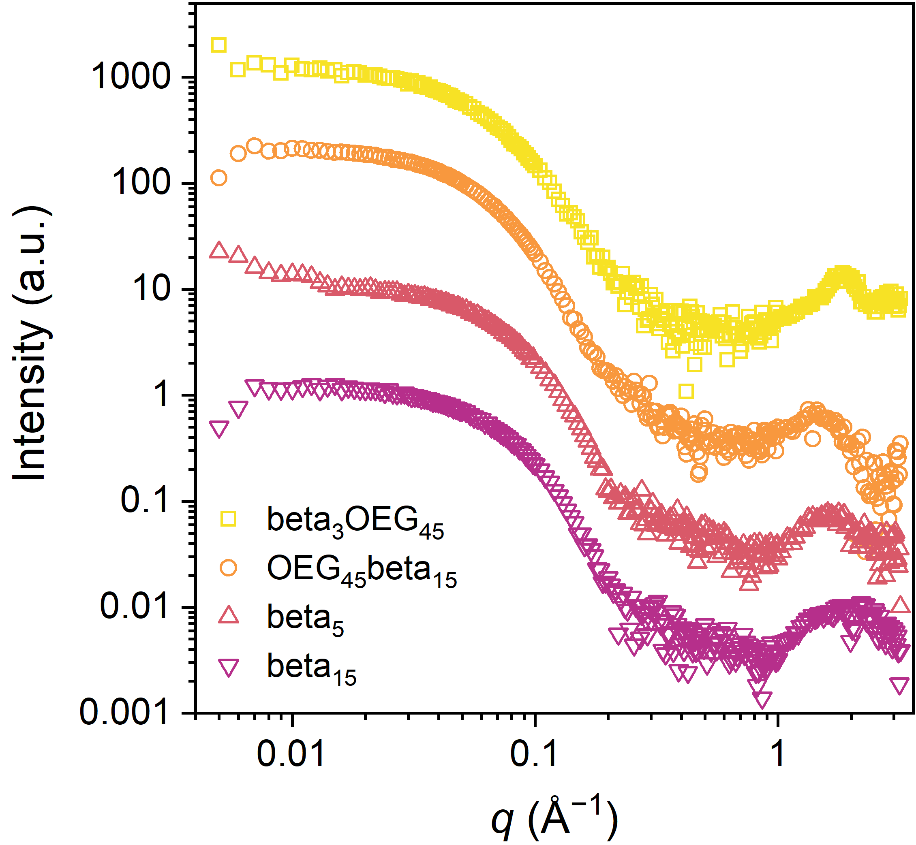

Figure S38. Background-subtracted small-angle X-ray scattering patterns of solutions of β-hairpin-based polymers with concentrations of 2.5 mg/mL in sodium acetate buffer. The polymers beta_5_, beta_15_, OEG_45_-*r*-beta_15_, and beta_3_OEG_45_ are indicated with maroon, red, orange, and yellow symbols, respectively. The wavelength of measurement is listed in Table S1. SAXS patterns besides beta_15_ are vertically offset for greater visual clarity.

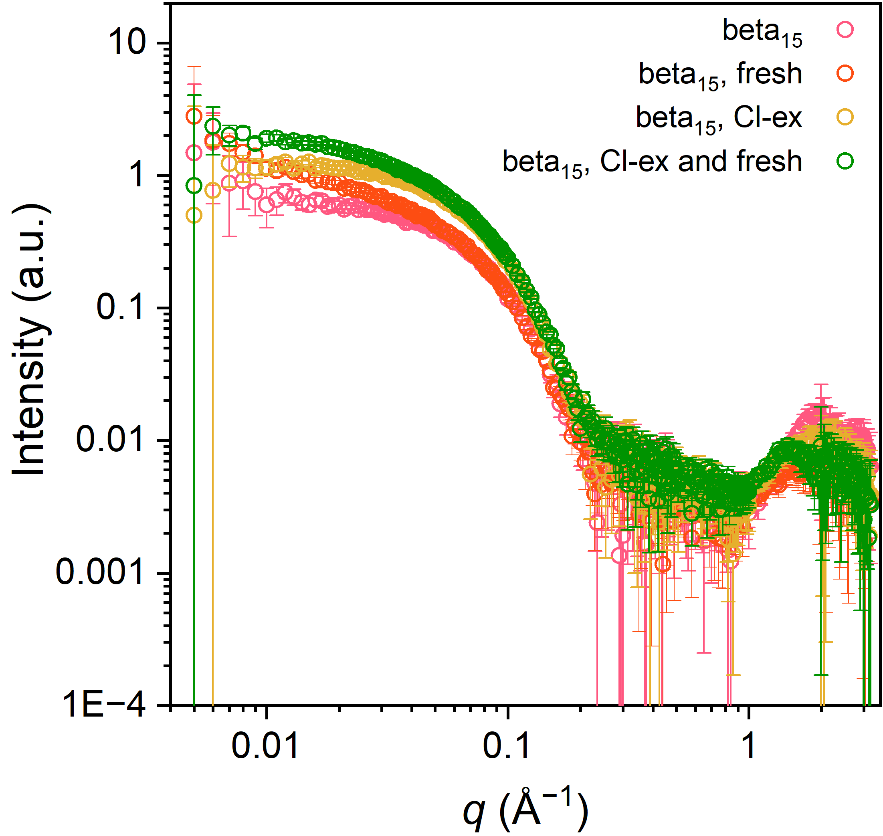

Figure S39. Background-subtracted small-angle X-ray scattering patterns of solutions of β-hairpin-based polymers with concentrations of 2.5 mg/mL in sodium acetate buffer. The polymers beta_15_, freshly prepared beta_15_, Cl-exchanged beta_15_, and freshly prepared Cl-exchanged beta_15_ are indicated with red, orange, yellow, and green symbols, respectively. Freshly prepared samples, which were not dialyzed against sample buffer, show a slight increase in slope at low *q* (<0.05 Å^−1^), which we postulate is due to insufficient equilibration with the sample buffer. The wavelength of measurement is listed in Table S1.

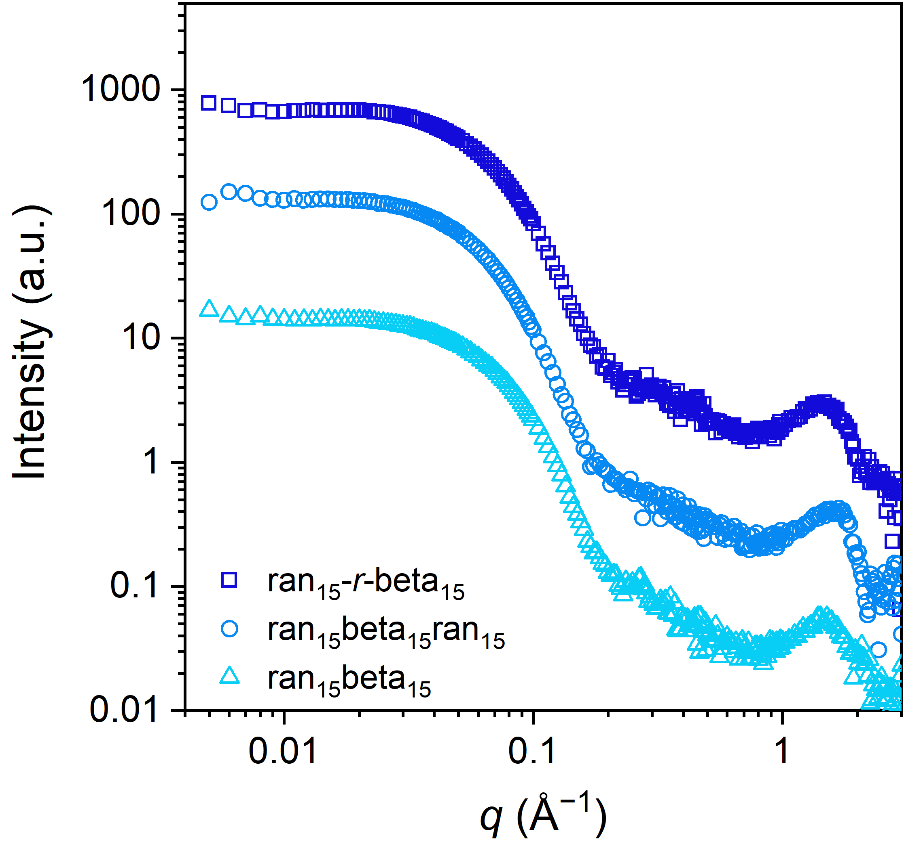

Figure S40. Background-subtracted small-angle X-ray scattering patterns of solutions of copolymers of the random and β-hairpin peptide monomers, with concentrations of 2.5 mg/mL in sodium acetate buffer. The polymers ran_15_beta_15_, ran_15_beta_15_ran_15_, and ran_15_-*r*-beta_15_ are indicated with light blue, blue, and navy symbols, respectively. The wavelength of measurement was 0.8172 Å. SAXS patterns besides ran_15_-r-beta_15_ are vertically offset for greater visual clarity.

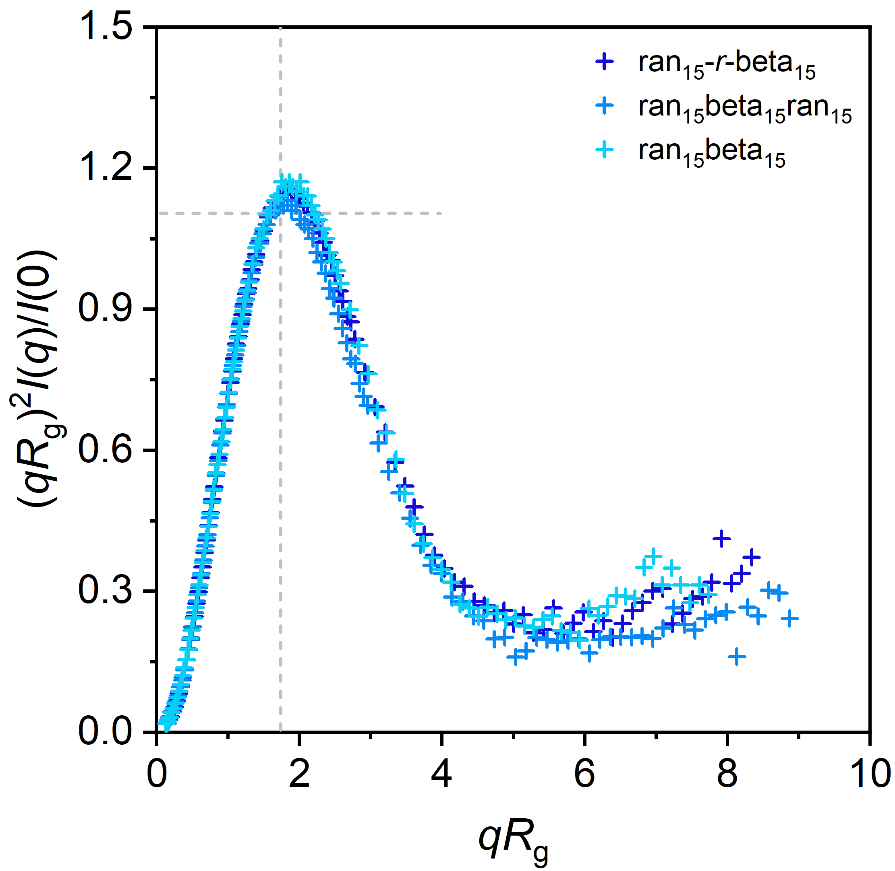

Figure S41. Normalized Kratky plots of background-subtracted SAXS data of solutions of copolymers of the random and β-hairpin peptide monomers, with concentrations of 2.5 mg/mL in sodium acetate buffer. The polymers ran_15_beta_15_, ran_15_beta_15_ran_15_, and ran_15_-*r*-beta_15_ are indicated with light blue, blue, and navy symbols, respectively. The wavelength of measurement was 0.8172 Å. Gray dashed lines indicate positions of *qR*_g_ and (*qR*_g_)2*I_(q)_*/*I*_(0)_ for an ideal compact sphere.

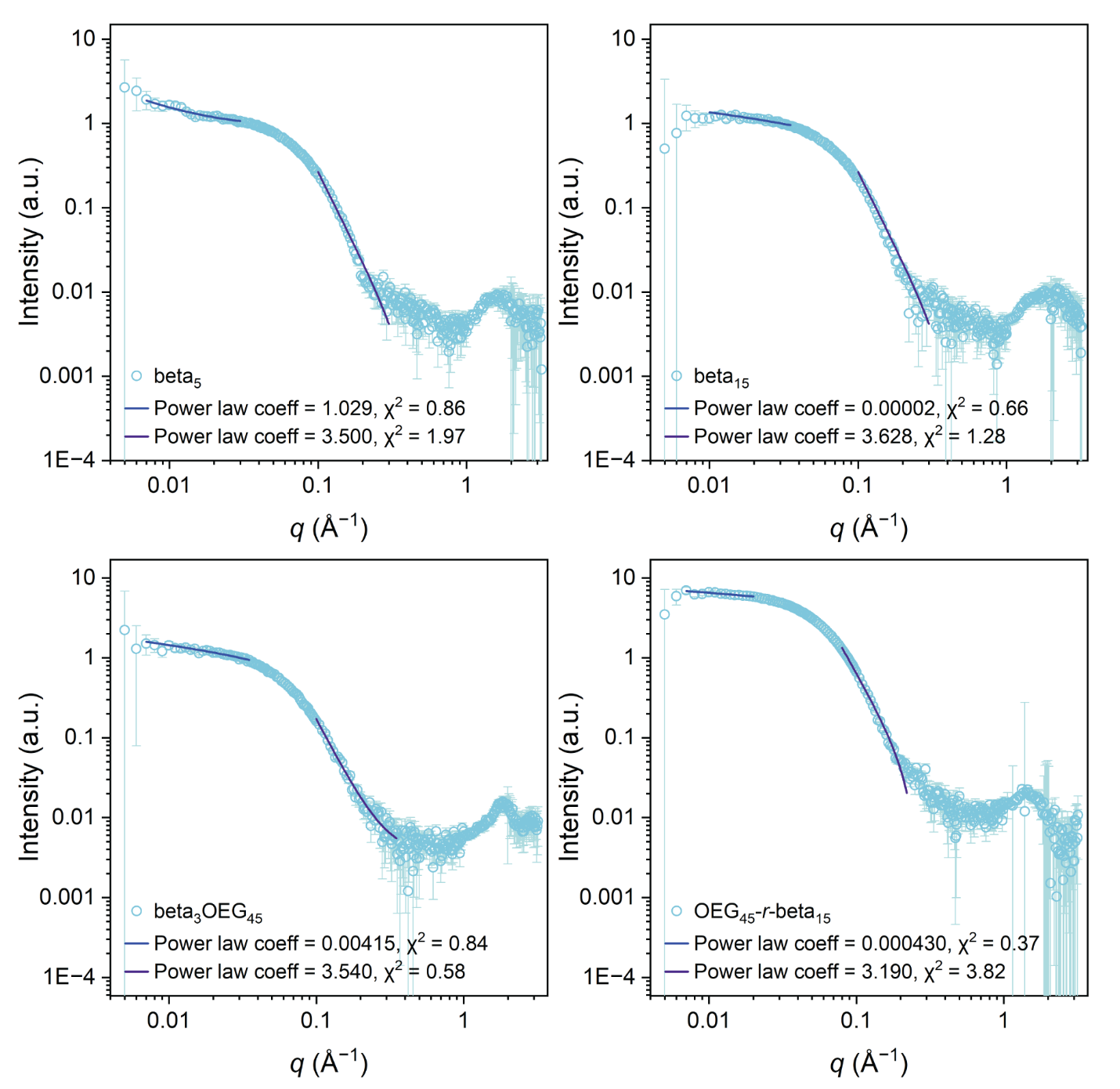

Figure S42. Background-subtracted small-angle X-ray scattering patterns of solutions of β-hairpin polymers (pale blue symbols), with concentrations of 2.5 mg/mL in sodium acetate buffer, compared to the power law model fits (blue and purple lines) as implemented in SasView.^22^ The wavelength of measurement is listed in Table S1.

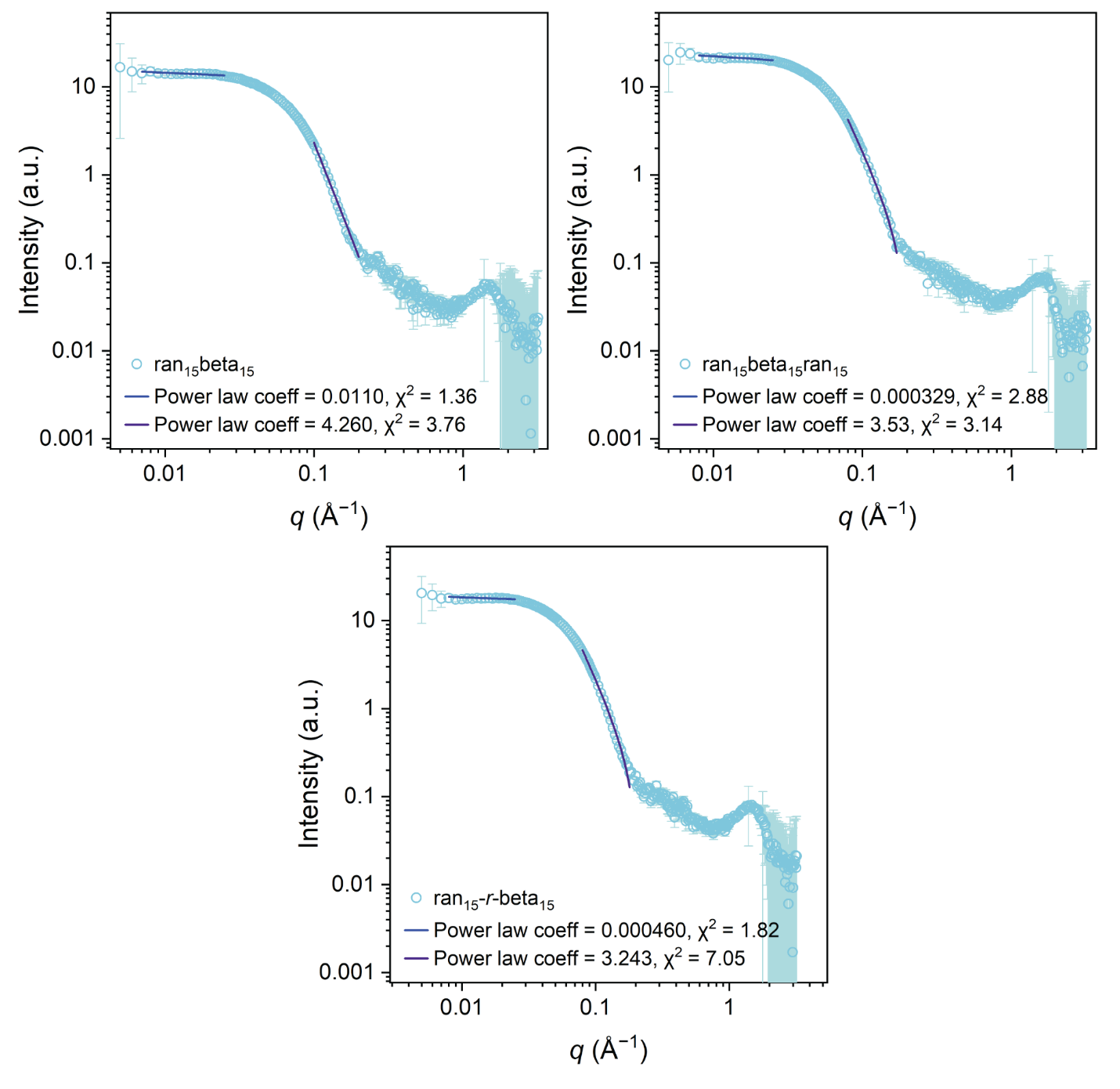

Figure S43. Background-subtracted small-angle X-ray scattering patterns of solutions of β-hairpin and random copolymers (pale blue symbols), with concentrations of 2.5 mg/mL in water, compared to the power law model fits (blue and purple lines) as implemented in SasView.^22^ The wavelength of measurement is listed in Table S1.

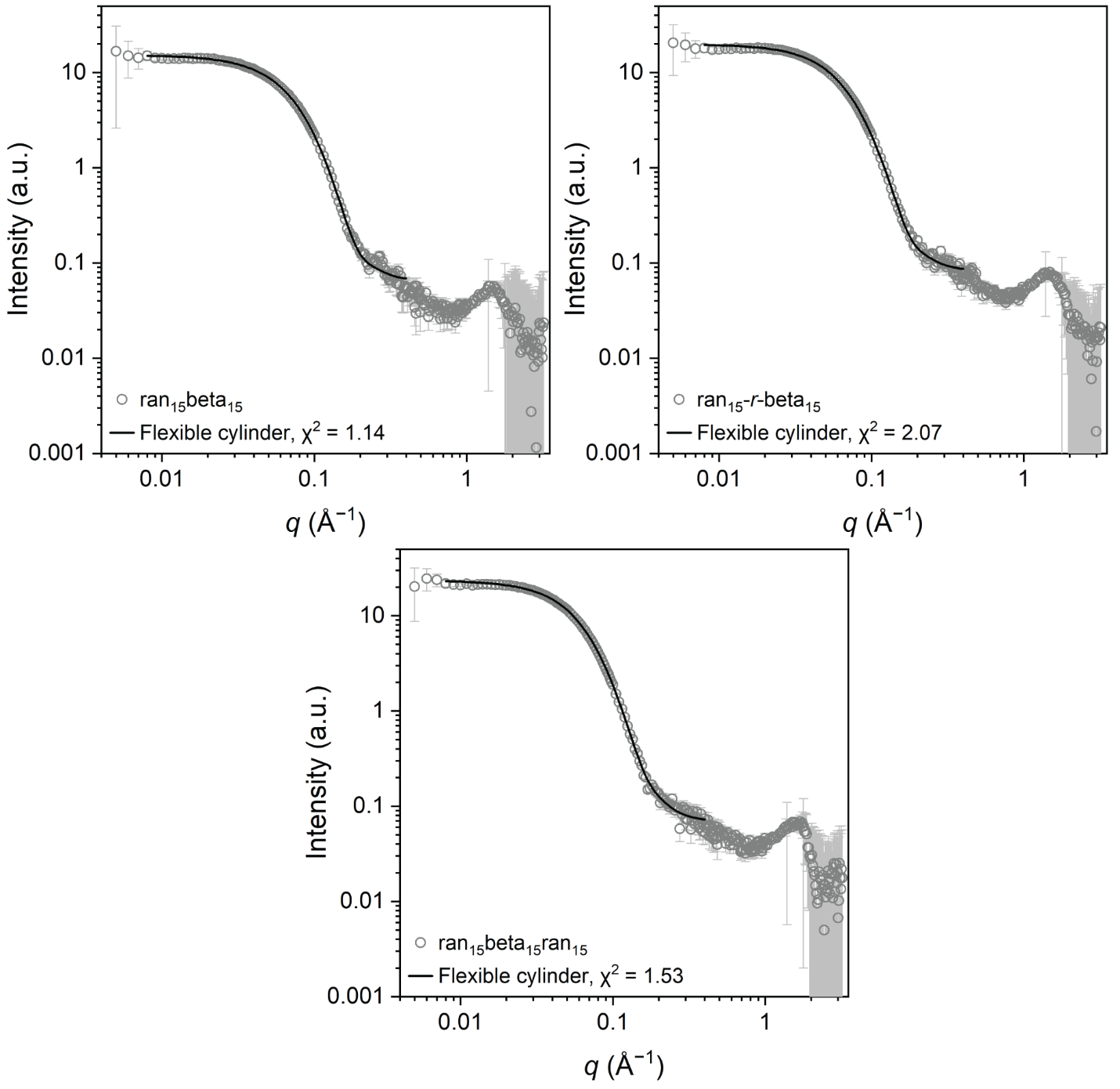

Figure S44. Background-subtracted small-angle X-ray scattering patterns of solutions of β-hairpin and random copolymers (gray symbols), with concentrations of 2.5 mg/mL in water, compared to the flexible cylinder model fits (black lines) as implemented in SasView.^22^ The wavelength of measurement is listed in Table S1.

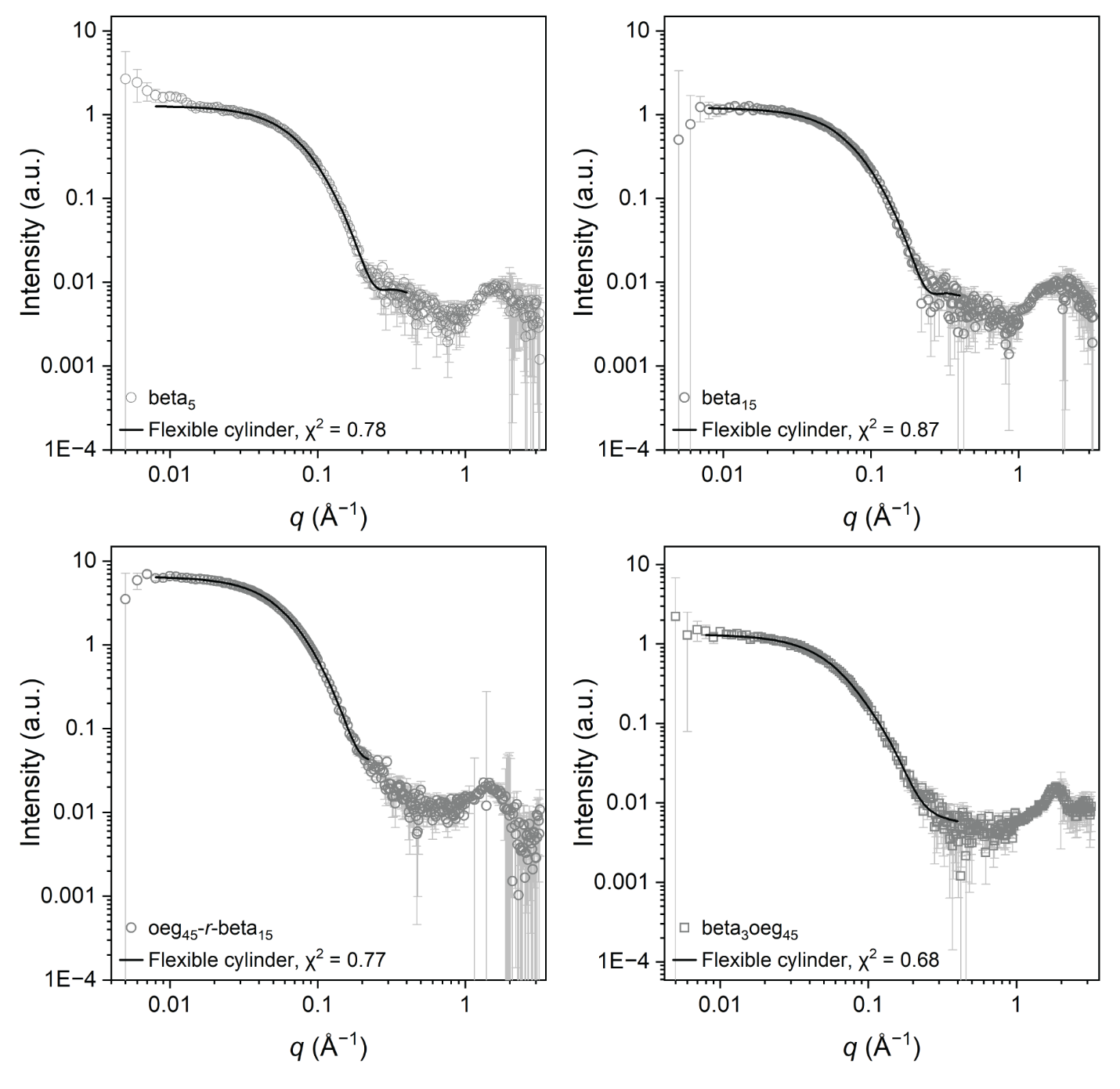

Figure S45. Background-subtracted small-angle X-ray scattering patterns of solutions of β-hairpin polymers (gray symbols), with concentrations of 2.5 mg/mL in sodium acetate buffer, compared to the flexible cylinder model fits (black lines) as implemented in SasView.^22^ The wavelength of measurement is listed in Table S1.
